# Method of loci training yields unique prefrontal representations that support effective memory encoding

**DOI:** 10.1101/2025.02.24.639840

**Authors:** Jingyuan Ren, Boris N. Konrad, Yannan Zhu, Fan Li, Michael Czisch, Martin Dresler, Isabella C. Wagner

## Abstract

The method of loci is a technique to effectively boost memory, but its impact on the underlying neural representations is poorly understood. Here, we used functional magnetic resonance imaging and representational similarity analysis to compare the neural representations of memory athletes ranked among the world’s top 50 in memory sports to those of mnemonics-naïve controls. In a second study, mnemonics-naïve individuals underwent a 6-week-long memory training, working memory training, or no intervention. Results showed distinct neural representations in the prefrontal cortex, inferior temporal, and posterior parietal regions as memory athletes and the memory training group studied novel content. Neural representations were also distinct between these experienced individuals, which was related to better memory performance after 4 months. In parallel, the data revealed increased neural pattern similarity in the anterior hippocampus and precuneus, suggesting a more generalised role of these regions in episodic memory formation and retrieval. Our findings highlight how extensive memory training affects neocortical memory engrams. We suggest that the method of loci may bolster memory uniqueness within one’s “memory palace”, setting the stage for exceptional memory performance.

## Introduction

Mnemonic techniques can substantially boost memory performance. One of the most commonly used techniques is the so-called method of loci, relying on mental navigation along well-known spatial routes (Yates, 1966). To-be-studied material is mentally placed at salient locations along the imagined route and can then be recalled by “retracing” the route, “picking up” the material where it was previously “dropped off”. Successful application of the method of loci requires extensive training and is often used by experts, or “memory athletes”, who demonstrate their superior abilities in memorising and accurately reproducing tremendous amounts of arbitrary information in competitions such as the World Memory Championships. Previously, our group showed that mnemonic-naïve individuals can also enhance their memory performance through multiple-week training (Dresler et al., 2017). The training effects were accompanied by changes in neural activation and connectivity of brain regions typically engaged in memory and visuospatial processing, including the hippocampus, posterior medial, and prefrontal regions (Wagner et al., 2021). Parallel to levels of activation and connectivity, superior memory might rely on distributed activation patterns that can be assessed with representational similarity analysis (RSA, Kriegeskorte et al., 2008). RSA quantifies the spatial alignment of activation patterns, and dissimilar patterns are thought to code for distinct informational content. For instance, Liu and colleagues (2022) demonstrated that method of loci training over the course of five days was associated with lower neural pattern similarity in hippocampal subfields during memory encoding (hence, distinct hippocampal patterns). This effect was specific to the studied material that was mentally placed at close-by locations along the imagined route and was coupled to accurate temporal order memory. Together, this highlights that short-term training with the method of loci disambiguates hippocampal representations, but the question remains of how extended, multiple-week training transforms the neural representations that underlie superior memory abilities as seen in memory athletes.

The processes that enable superior memory through method of loci training may also impact the similarity of neural patterns between individuals. This is grounded upon recent work demonstrating that the degree of neural pattern similarity to an average, group-based neural pattern during associative encoding predicted subsequent memory ability across individuals (Sheng et al., 2023). Put differently, the more the individuals resembled the group in their neural representations, the better their memory was. Increased neural pattern similarity between individuals was also reported during successful scene encoding (Koch et al., 2020) and associative retrieval (Xiao et al., 2020). Such “neural alignment” between individuals might stem from the processing of similar content. For instance, individuals display neural alignment when watching the same video (Chen et al., 2017) or when sharing beliefs about the same story (Yeshurun et al., 2017). In educational settings, neural alignment between students is tied to knowledge acquisition of abstract physics concepts (Mason & Just, 2016) and indicates effective information transmission in the classroom (Nguyen et al., 2022). Taking this a step further, Meshulam and colleagues (2021) not only showed neural alignment between learners after acquiring new conceptual knowledge but also reported neural alignment between learners and experts that was predictive of learning outcomes. Whether superior memory after method of loci training is coupled to neural alignment between individuals (and, specifically, between trainees and memory athletes, akin to learners and experts) is currently unclear. Furthermore, the notion of neural alignment appears difficult to reconcile with the finding of distinct neural representations after method of loci training (Liu et al., 2022), raising the possibility that neural alignment might be less beneficial when memory specificity and flexible order recognition are at stake.

We addressed these knowledge gaps in two separate studies (for an overview, see Fig. 1; and see also Dresler et al., 2017; Wagner et al., 2021 who used the same data sets). In the first study (the “athlete study”), we included individuals who were experts in using the method of loci (hereafter referred to as “memory athletes”) and who were ranked among the world’s top 50 in memory sports (*n* = 17). These were compared to a control group that was closely matched for age, sex, handedness, and intelligence (*n* = 16; see Table S1 for demographic details). To assess neural representations during memory encoding and retrieval, participants underwent functional magnetic resonance imaging (fMRI) while studying novel words (word list encoding task) and were subsequently tested for their ability to recognise their temporal order (temporal order recognition task). We specifically opted for these tasks as they are commonly used during memory championships, and because we reasoned that the particular strength of the method of loci lies in the learning and recalling of ordered sequences (due to mental navigation through the “memory palace”). Memory performance was assessed through a free recall test after fMRI. In the second study (the “memory training study”), we enrolled mnemonics-naïve individuals who completed a 6-week method of loci training (memory training group, *n* = 17). These individuals were compared to two control groups that completed a 6-week working memory training (active controls, *n* = 16) or no intervention (passive controls, *n* = 17). All participants completed the abovementioned memory tasks during fMRI before and after the 6-week period to determine any (training-related) changes in neural representations. Memory performance was assessed in a free recall test post-fMRI and after 24 hours, as well as in a re-test after four months.

**Fig. 1.**
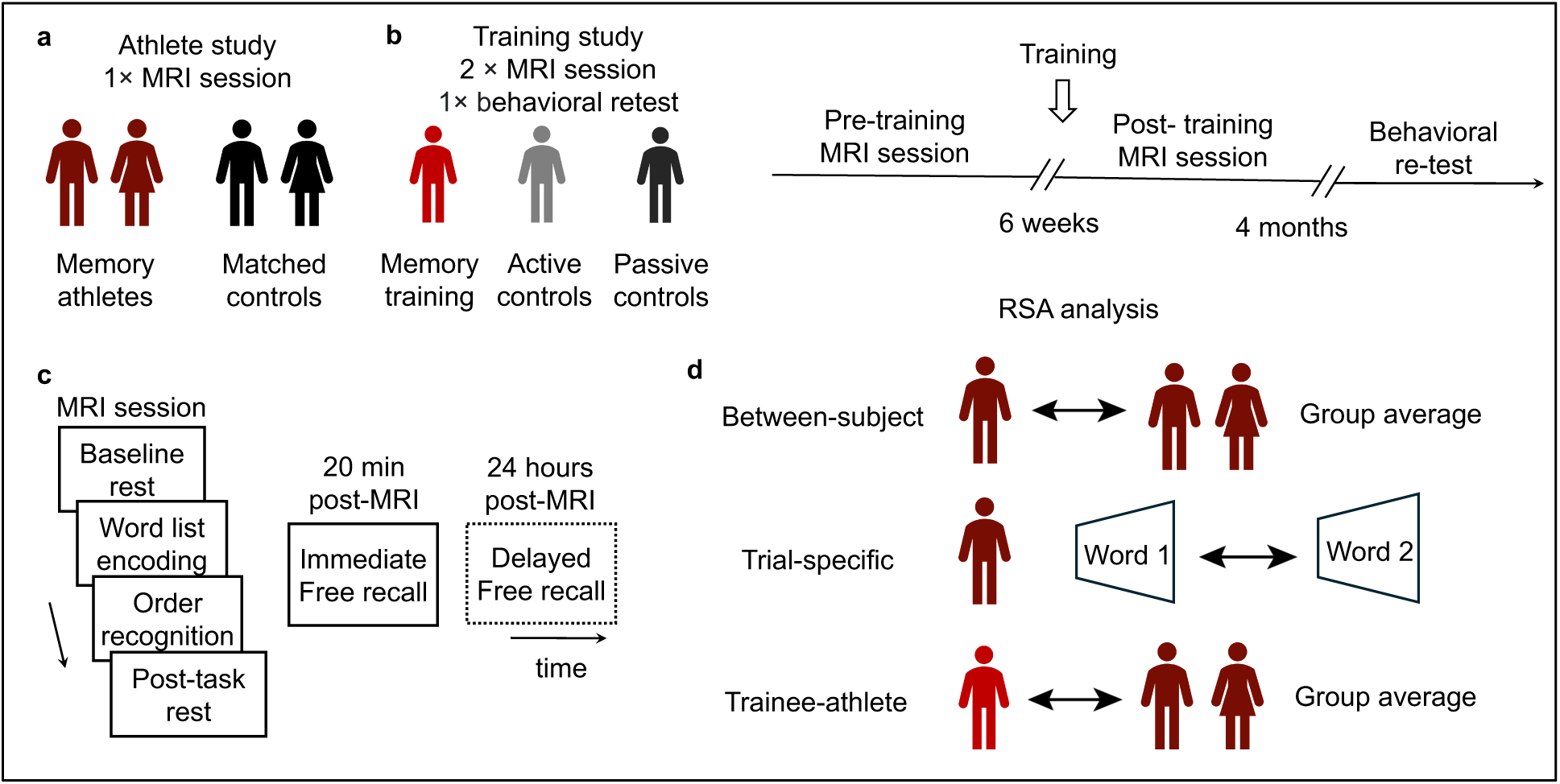
Overview of athlete and training studies, and representational similarity analysis. (**a**) We examined memory athletes (*n* = 17) and compared them to matched controls (*n* = 16) in a single MRI session (athlete study). (**b**) Individuals in the training study were assigned to one of three groups (memory training group, *n* = 17, active controls, *n* = 16, passive controls, *n* = 17). They completed two MRI sessions prior and after the 6-week-long interval (pre- and post-training sessions), and a behavioural re-test after 4 months. (**c**) General structure of all MRI sessions: baseline and post-task rest periods (not discussed further), word list encoding and temporal order recognition tasks (10 min each), immediate (20 min post-MRI) and delayed (24 hours post-MRI) free recall (dashed frame indicates that the delayed test was only included in the training study). (**d**) We conducted three types of representational similarity analysis (RSA). Between-subject RSA: this assessed the neural pattern similarity between individuals within the same experimental group. Trial-specific RSA: this examined neural pattern similarity across the studied words, but within an individual. Trainee-athlete RSA: this evaluated the degree of neural pattern similarity between trainees and memory athletes. Note that the schematic shows memory athletes and individuals of the memory training group (the “trainees”) only, but the same analysis was performed for all remaining groups as well.

Since the method of loci involves the active formation of unique, salient associations (Liu et al., 2022), we hypothesised distinct neural representations for the verbal material that was studied during the word list encoding task. This effect should be specific to individuals who completed the method of loci training (thus, memory athletes and the memory training group, or the so-called “trainees”). Using the existing data sets (Dresler et al., 2017; Wagner et al., 2021), our novel analysis (which is orthogonal to prior work) was focused on the training-related changes in neural representations between individuals. We reasoned that the distinct neural representations on the level of the studied material should impact the degree of neural alignment between those who completed the method of loci training. This allowed us to directly test whether trainee-athlete neural alignment during encoding is beneficial when memory specificity and temporal order recognition are at stake. Lastly, we leveraged the data from the recognition task. Since we were unsure how method of loci training would affect the neural representations when individuals retrieved the previously learned word-location associations, we opted for an exploratory analysis. Overall, our approach provides first insights into how extended memory training transforms the neural representations that give rise to superior memory.

## Results

### Memory training is associated with distinct neural representations between experienced individuals during word list encoding

Building on our previous work that showed increased free recall performance after method of loci training (Dresler et al., 2017) as well as enhanced durable memory formation (Wagner et al., 2021), we here revisited the fMRI data to investigate potential changes in the underlying neural representations that might arise after memory training. Because successful memory formation after shorter-term method of loci training was shown to be associated with distinct neural representations of the specific study material (Liu et al., 2022), we hypothesised that this would impact the degree of neural alignment between experienced individuals. In other words, method of loci training should lead to distinct neural representations (thus, lower neural alignment) between memory athletes and between individuals of the memory training group after training while they studied novel words (word list encoding task; Fig. 2b). During the task, memory athletes and individuals of the memory training group (post-training) were asked to use the method of loci (they were asked to mentally navigate through their memory palace and to “place” the studied words at the specific location). We quantified the degree of neural alignment by means of representational similarity analysis (RSA) and focused on the lateral prefrontal cortex and the superior frontal gyrus that we previously had found to be associated with word list encoding following method of loci training (Wagner et al., 2021). For each individual and region-of-interest (ROI), we extracted the average neural pattern during word list encoding (reflecting encoding-related activity relative to the implicit fixation cross baseline). This was then correlated with the neural patterns of all other individuals of the same experimental group (e.g., memory athletes × memory athletes, and memory training group × memory training group, separately for pre- and post-training), yielding the magnitude of neural pattern similarity per group and time point (Fig. 2a, see section “Between-subject RSA” in the Methods).

**Fig. 2.**
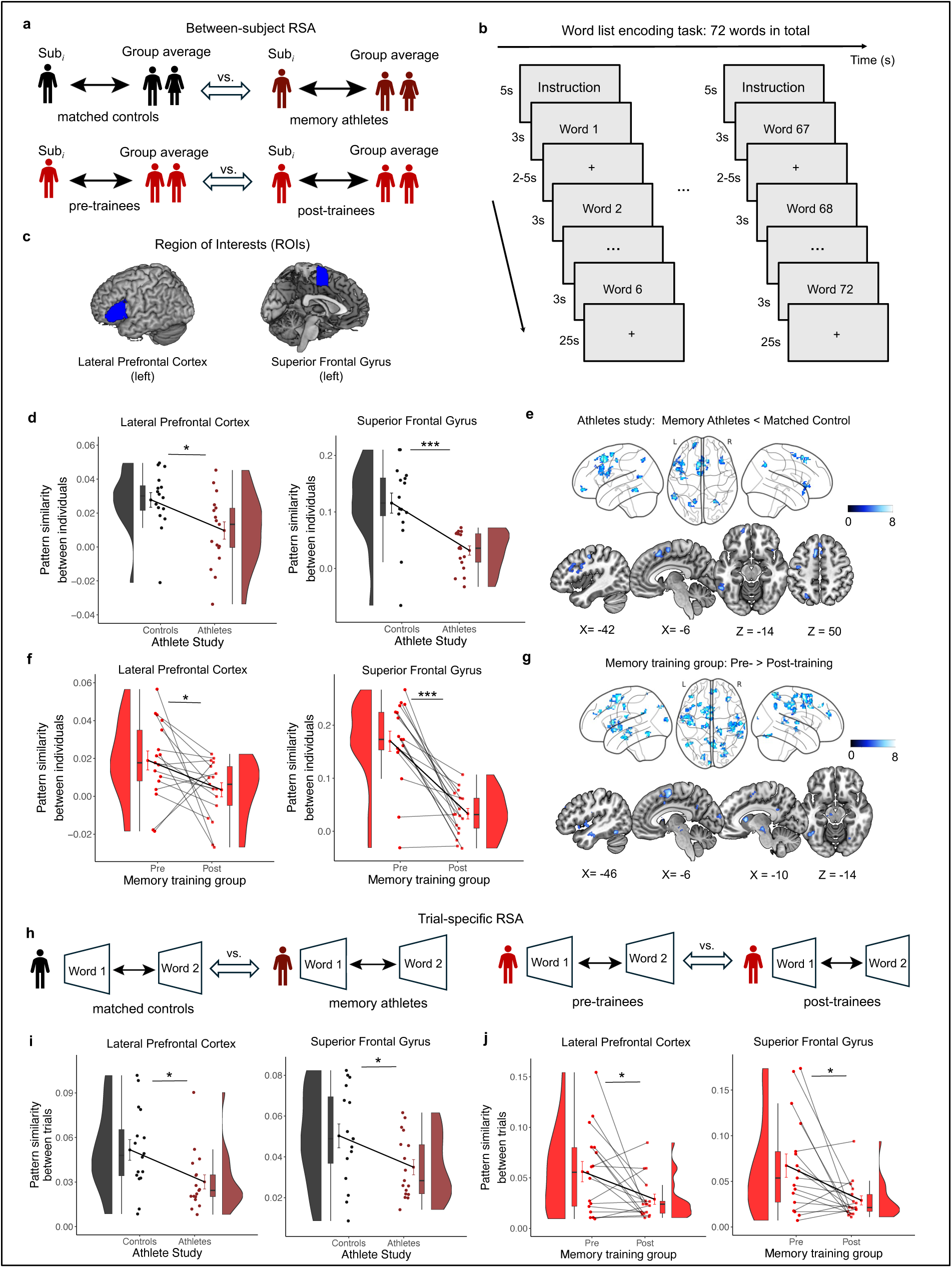
Method of loci training yields distinct neural representations between experienced individuals during word list encoding. (**a**) Detailed schematic of the between-subject RSA: We compared the individual-to-group neural pattern similarity between memory athletes (*n* = 17) and matched controls (*n* = 16) during word list encoding (relative to the implicit fixation cross baseline), and between pre- and post-training within the memory training group (*n* = 17, see Methods, and see supplementary materials for control analyses). (**b**) Schematic of the word list encoding task. A trial was defined as the presentation of one word. The 72 trials were organised into 12 blocks of 6 words each. A block began with an instruction screen presented for 5 seconds, followed by the 6 words, each presented for 3 seconds and separated by a jittered fixation cross of 2-5 seconds. After every block, a fixation cross was displayed for 25 seconds. (**c**) Regions of interest (ROIs) that were used for all analyses pertaining to word list encoding. (**d, f**) ROI results of the between-subject RSA (control analyses for the active and passive control groups are shown in Fig. S1). (**e, g**) Searchlight results of the between-subject RSA. Results were thresholded at *p* < 0.05, family-wise error (FWE)-corrected at cluster-level, using a cluster-defining threshold of *p* < 0.001 (Tables S2-3). (**h**) Detailed schematic of the trial-specific RSA: We calculated neural pattern similarity on a trial-by-trial basis (relative to the implicit fixation cross baseline) for each participant and compared memory athletes to matched controls, and pre- to post-training time points in the memory training group (same *n* as above). (**i, j**) ROI results of the trial-specific RSA (control analyses for the active and passive control groups are shown in Fig. S3). (**d, f, i, j**) Data points represent individual participants (athlete study, *n* = 33, memory training group, *n* = 17). Boxplots display the median and upper/lower quartiles, with whiskers extending to the most extreme data points within 1.5 interquartile ranges above/below the quartiles. The data points with error bars show the mean ± standard error of the mean (S.E.M.). Data distributions are based on the respective probability density function. All ROI-based results survived multiple comparison corrections using the false discovery rate (FDR) with the Benjamini-Hochberg procedure set to *q* < 0.05. \*\*\**p* < 0.001; \**p* < 0.05; ns, not significant. Pattern similarity reflects Fisher *z*-transformed Pearson correlations (*r*).

First, we investigated data from the athlete study. Results revealed significantly lower neural pattern similarity between memory athletes compared to matched controls in both the left lateral prefrontal cortex (separate independent-sample *t*-tests, memory athletes, *n* = 17, matched controls, *n* = 16, *t*(31) = 2.690, *p* = 0.011, Cohen’s *d* = 0.937; Fig. 2d) and the left superior frontal gyrus during word list encoding (*t*(31) = 4.350, *p* < 0.001, Cohen’s *d* = 1.517; Fig. 2d). To test for effects beyond these pre-defined ROIs, we repeated our analysis but instead moved a spherical searchlight throughout the entire brain volume (see section “Searchlight-based RSA” in the Methods). This confirmed the abovementioned results and further showed significantly lower neural pattern similarity in the left inferior temporal cortex, the superior parietal and precentral gyrus in memory athletes compared to matched controls (independent-sample *t*-test, same samples as above, contrasts word list encoding > baseline in memory athletes < matched controls, *p* < 0.05, family-wise error (FWE) corrected at cluster-level using a cluster-defining threshold of *p* < 0.001, critical cluster size = 36 voxels, Fig. 2e, Table S2). Thus, in line with our expectations, we found distinct neural representations (lower neural alignment) between memory athletes compared to matched controls in brain regions related to successful memory encoding and visuospatial processing when studying novel words.

Next, we turned to the data of the training study. Similar to the findings above, we observed significantly lower neural pattern similarity between the individuals of the memory training group after training (compared to the pre-training baseline) in the left lateral prefrontal cortex (separate paired-sample *t*-tests, *n* = 17, *t*(16) = 2.200, *p* = 0.043, Cohen’s *d* = 0.533, Fig. 2f) and the left superior frontal gyrus (*t*(16) = 7.240, *p* < 0.001, Cohen’s *d* = 1.756, Fig. 2f). This effect was specific to the memory training group after training but was not present in the active or passive control groups (Fig. S1). Whole-brain searchlight analysis confirmed these findings and further revealed lower neural pattern similarity in individuals of the memory training group after training (compared to the pre-training baseline) in the left inferior temporal cortex, caudate nucleus, pre- and postcentral gyri, the superior temporal and parietal gyrus, among others (paired-sample *t*-test, *n* = 17, contrast word list encoding > baseline during pre-training > post-training, *p* < 0.05, FWE-corrected, cluster-defining threshold *p* < 0.001, critical cluster size = 30 voxels, Fig. 2g, Table S3). Neural pattern similarity in the memory training group was also lower as compared to the active and passive controls (Fig. S2). Overall, we found that studying word lists while utilising the method of loci was coupled to distinct neocortical and primarily prefrontal representations between memory athletes and between individuals of the memory training group after training.

Given the relatively large size of the lateral prefrontal cortex ROI (2161 voxels) and its known functional heterogeneity (Koechlin et al., 2000, 2003; Nee & D’Esposito, 2016; Schumacher et al., 2019), we examined whether training-related effects were localised to specific subregions along the dorsal-to-ventral axis (see Fig. S9a). This analysis showed lower neural pattern similarity in the dorsal and ventral lateral prefrontal regions in memory athletes and trainees (post-training), respectively (see Supplementary Results S1, which also includes a brief discussion, Fig. S10, Table S10).

Because of the known medial temporal lobe (MTL) involvement in method of loci processing (Fellner et al., 2016; Liu et al., 2022; Maguire et al., 2003; Nyberg et al., 2003), we also conducted additional ROI analysis focused on the hippocampus and parahippocampal gyrus. Surprisingly, we found increased (rather than decreased) neural pattern similarity in both the hippocampus (memory athletes) and parahippocampal gyrus (memory training group after training; see Supplementary Results S2, Fig. S15). We reasoned that these effects were potentially driven by sequence processing (which should be common to all experienced individuals as they mentally navigated along the loci routes to memorise the novel word-location associations; note that this analysis was only performed using the training study data). By employing a sequence-geometry analysis, we hence tested for a monotonic decrease in neural pattern similarity as a function of distance between word-location associations along the loci route, but the results were not significant (Supplementary Results S6, which also includes a brief discussion, Fig. S20).

Next, we reasoned that if the MTL was engaged in the encoding of unique word-location associations, we might be able to detect decreases in neural pattern similarity by employing an alternative between-subject analysis approach that emphasised single-trial characteristics (which was not the case for the abovementioned main analysis). We thus performed a trial-wise intersubject consistency (ISC) analysis (memory athletes used their own loci routes, which should be coupled to maximally unique neural representations and decreased neural pattern similarity between them; note that this analysis was only performed using the athlete study data), but results were not significant (Supplementary Results S6, which also includes a brief discussion, Fig. S20).

### Experienced individuals hold unique neural representations associated with the studied words

To verify that the finding of distinct neural representations in the prefrontal cortex between experienced individuals was related to distinct neural representations associated with each studied word (Liu et al., 2022), we focused on the same *a priori* ROIs as above (left lateral prefrontal cortex and superior frontal gyrus) and quantified the average neural pattern similarity across all studied words per participant (see section “Trial-specific RSA” in the Methods and Fig. 2h). In contrast to the between-subject RSA, this analysis focused on neural pattern similarity *between different words* during the word list encoding task and was conducted on a per-subject basis, assessing whether method of loci training led to shared or distinct neural representations across items.

Results indeed revealed significantly lower neural pattern similarity associated with the studied words in memory athletes compared to matched controls in both the left lateral prefrontal cortex (separate independent-sample *t*-tests, memory athletes, *n =* 17, matched controls, *n* = 16, *t*(31) = 2.590, *p* = 0.014, Cohen’s *d* = 0.904, Fig. 2i) and the left superior frontal gyrus (*t*(31) = 2.230, *p* = 0.033, Cohen’s *d* = 0.778; Fig. 2i). Returning to the data from the training study, we observed significantly lower neural pattern similarity associated with the studied words in the memory training group after training (compared to the pre-training baseline) in both the left lateral prefrontal cortex (separate paired-sample *t*-tests, *n* = 17, *t*(16) = 2.530, *p* = 0.022, Cohen’s *d* = 0.613; Fig. 2j) and the left superior frontal gyrus (*t*(16) = 2.800, *p* = 0.013, Cohen’s *d* = 0.679; Fig. 2j). This effect was only present in the memory training group after training, but not in the active or passive controls (Fig. S3). Again, the effect of lower neural pattern similarity was localised to the dorsal and ventral subregions of the lateral prefrontal cortex in memory athletes and trainees (post-training), respectively (see Supplementary Results S1, Fig. S11, and Table S10; and see Supplementary Results S2, Fig. S16 for additional MTL analysis that did not reveal any significant results).

Thus, the distinct neural representations in the prefrontal cortex between individuals who were proficient in the method of loci appeared driven by distinct neural representations associated with each studied word.

### Distinct neural representations between trainees and memory athletes during word list encoding

Previous work showed that the degree of neural alignment between learners and experts was predictive of learning outcomes (Meshulam et al., 2021), and we were eager to test whether similar mechanisms are at play when it comes to expertise with the method of loci. So far, we have shown unique neural representations for the studied material, distinct neural representations between individuals of the memory training group after training (thus, in the so-called “trainees”), and between memory athletes. Based on these results, we expected distinct neural representations between trainees and memory athletes, which would indicate that superior memory relies on neural processes that are specific to those who are proficient in utilizing the method of loci. To quantify the degree of trainee-athlete similarity in neural representations, we extracted the average neural patterns during word list encoding for each participant of the memory training group. Analyses were performed using the abovementioned *a priori*-defined ROIs (left lateral prefrontal cortex and superior frontal gyrus). We then correlated the neural patterns with those of each memory athlete, and, separately, with those of each individual of the matched control group. This allowed us to compare the average neural pattern similarity between *time points* (pre-vs. post-training) and *groups* (neural pattern similarity between trainees × athletes vs. trainees × matched controls; Fig. 3a, see section “trainee-athlete RSA” in the Methods).

**Fig. 3.**
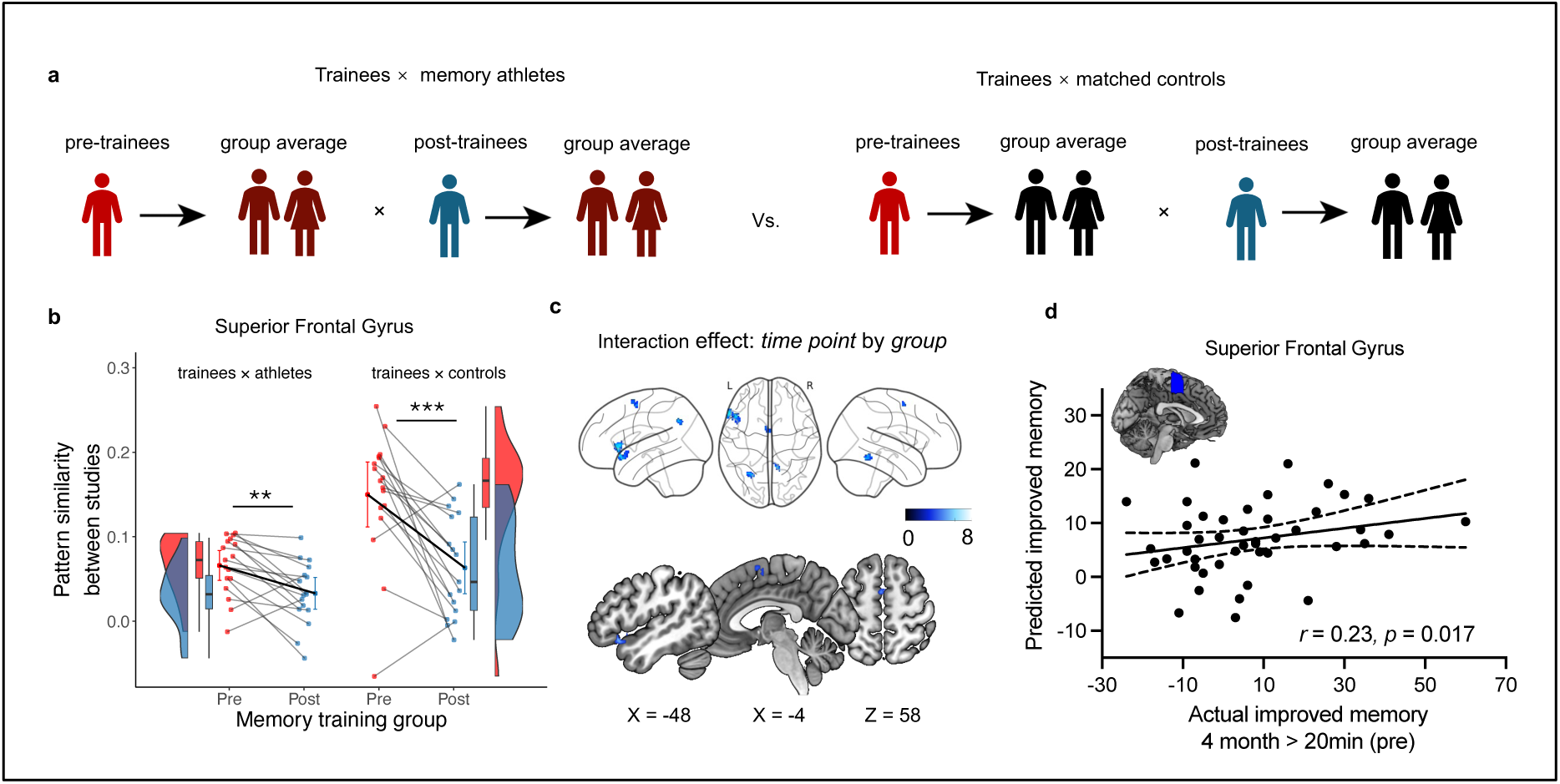
Method of loci training yields distinct neural representations between trainees and memory athletes during word list encoding. (**a**) Detailed schematic of the trainee-athlete RSA: We calculated the trainee-to-athletes (“trainees” are individuals of the memory training group) and trainee-to-controls neural pattern similarity during word list encoding (relative to the implicit fixation cross baseline) and compared between pre-(red) and post-training (blue) time points (see Methods). (**b**) ROI results (control analyses for the active and passive control groups are shown in Fig. S4). Data points represent individual participants (memory training group, *n* = 17). Boxplots display the median and upper/lower quartiles, with whiskers extending to the most extreme data points within 1.5 interquartile ranges above/below the quartiles. The data points with error bars show the mean ± S.E.M. Data distributions are based on the respective probability density function. (**c**) Searchlight results for the time point × group interaction. Results were thresholded at *p* < 0.05, FWE-corrected at cluster-level, using a cluster-defining threshold of *p* < 0.001 (Table S4). (**d**) The scatter plot depicts the correlation between memory performance at the 4-month reset (4 months > immediate free recall test during the pre-training session) and the predicted changes in neural pattern similarity between all individuals of the training study and memory athletes from pre- to post-training (*n* = 44). \*\*\**p* < 0.001; \*\**p*< 0.01; ns, not significant. Pattern similarity reflects Fisher *z*-transformed Pearson correlations (*r*).

During word list encoding, we observed significantly lower neural pattern similarity in the left superior frontal gyrus when trainees had completed the method of loci training (repeated measures ANOVA, *n* = 17, main effect of *time point*: *F* (1, 16) = 19.001, *p* < 0.001, *η_p_*^2^ = 0.54, and significantly lower neural pattern similarity between trainees × athletes as compared to between trainees × matched controls (main effect of *group*: *F* (1, 16) = 29.577, *p* < 0.001, *η_p_*^2^ = 0.65). We also found a significant *time point* × *group* interaction (*F* (1, 16) = 12.080, *p* < 0.001, *η_p_*^2^ = 0.43). Simple effects showed that neural pattern similarity between trainees × athletes (*p* = 0.002) and between trainees × matched controls (*p* = 0.001) was lower in the post-training compared to the pre-training session (Fig.3b). Whole-brain searchlight analysis confirmed these findings (flexible factorial analysis with the factors *time point* and *group*, *n* = 17, *p* < 0.05, FWE-corrected, cluster-defining threshold *p* < 0.001, critical cluster size = 38 voxels, Fig. 3c, Table S4). These effects were only present in the memory training group but not in the active or passive control groups (Fig. S4). We also addressed potentially more localized effects in lateral prefrontal subregions (see Supplementary Results S1) and MTL effects (see Supplementary Results S2, Fig. S17), but this did not reveal any significant results.

Therefore, results suggest that method of loci training leads to distinct neural representations between trainees and memory athletes when studying novel words.

### Distinct neural representations during word list encoding are associated with better free recall performance after 4 months

We then tested the behavioural relevance of the distinct neural representations between all individuals of the training study and memory athletes at the time of encoding with respect to longer-term memory performance. Individuals of the training study were re-invited to the behavioural laboratory to participate in a re-test after 4 months. During this session, they completed the word list encoding task once more, followed by a free recall test, which we leveraged to derive a measure of free recall performance (see Methods). Our behavioural data showed that only the athletes and memory training group exhibited a significant improvement in free recall performance from baseline to the 4-month re-test, while no such improvement was observed in the other groups (see memory performance per group in Table S8). To further explore the relationship between neural representations and long-term memory improvement, we included all individuals of the training study (i.e., all three groups) in the prediction analysis, rather than restricting it to the method of loci group alone. This ensured that the analysis captured both the specific effects of method of loci training and broader training-related mechanisms across all groups, while enhancing statistical power and generalizability. We next employed cross-validated machine-learning (Cohen, 2010; Supekar et al., 2013) to predict the increase in free recall performance due to method of loci training (i.e., the change in free recall performance from baseline, measured 20 minutes post-fMRI during the pre-training session, to the 4-month re-test) based on the changes in neural pattern similarity between trainees × athletes from pre- to post-training during word list encoding (see section “Prediction analysis” in the Methods and trainee × athlete effects in Fig. 3a).

Results showed that decreased neural pattern similarity between all individuals of the training study and memory athletes in the left superior frontal gyrus was significantly associated with better free recall performance after 4 months (*r*_(predicted, observed)_ = 0.23, *p* = 0.017, *n* = 44, Fig. 3d). We also repeated the analysis by defining the change in free recall performance after 24h, which gave virtually identical results (Table S9; but note that stratified, within-group analysis did not reveal any significant associations, potentially due to the small group sizes and generally high, close-to-ceiling performance in the memory training group after training; see Supplementary Results S3, Table S12). To move beyond the predefined ROIs, we also performed whole-brain, voxel-wise linear regression using individual memory improvement as a covariate of interest (Supplementary Results S3). This yielded lower neural pattern similarity in the dorsolateral prefrontal cortex during word list encoding that was associated with better free recall performance after 4 months, and that was specific to the memory training group after training (Fig. S18).

This shows that method of loci training led to distinct, encoding-related neural representations, specifically between experienced individuals, that expressed a longer-lasting benefit on memory performance.

### Memory training is associated with shared neural representations between experienced individuals during temporal order recognition

Following word list encoding, participants engaged in a temporal order recognition task where they were presented with word triplets from the previously studied word lists and were asked to indicate whether words were presented in the same or in a different order than during study (Fig. 4a; for an overview of task performance, see Supplementary Results S5, Table S13). Memory athletes and individuals from the memory training group (post-training) were instructed to employ the method of loci during this task. It is important to highlight that this analysis can yield novel insights as to whether method of loci training affected general retrieval processes during temporal order recognition, rather than focusing on the “uniqueness” of each word-location association (as we have done so far). This is because temporal order judgments were based on word triplets (rather than sequences of single words), thereby amplifying the neural processes that are shared across the different items (i.e., the analysis picks up upon common, retrieval-related aspects that are similar across different words). Nevertheless, based on our results from word list encoding, we expected distinct neural representations (thus, lower neural alignment) between memory athletes and between individuals of the memory training group after training. We focused our analysis on brain regions that we previously found to be associated with temporal order recognition following method of loci training, such as the hippocampus and precuneus (Wagner et al., 2021). Between-subject RSA was performed identically to the word list encoding task, quantifying neural pattern similarity (relative to baseline that involved the syllable-counting control task) by correlating the ROI-based (Fig.4b) neural patterns within each of the experimental groups (e.g., memory athletes × memory athletes, and memory training group × memory training group, separately for pre- and post-training; Fig. 2a, and see section “between-subject RSA” in the Methods).

**Fig. 4.**
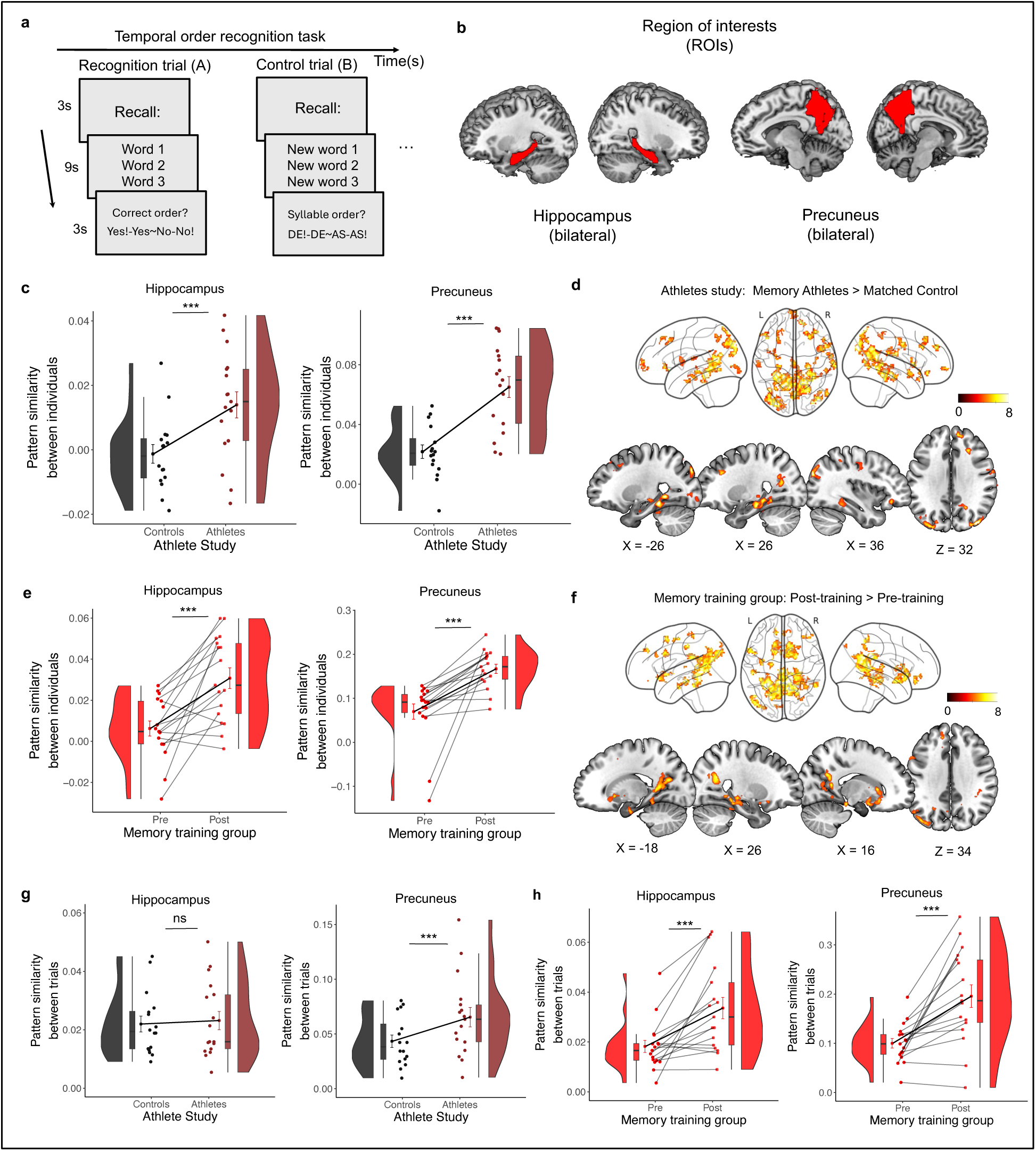
Method of loci training yields shared neural representations between experienced individuals during temporal order recognition. (**a**) Schematic of the temporal order recognition task. Participants viewed 24 triplets taken from the previously encoded word list. During recognition trials, participants judged whether the order of words in each triplet matched their original presentation during word list encoding. Main task trials were interspersed with control trials (in an alternating ABAB sequence), during which triplets of novel words were presented. Participants were asked to indicate whether these words were arranged in ascending or descending order based on the number of syllables (syllable-counting task). All trials began with a cue that signalled the start of the trial (3 s), followed by the triplet presentation (9 s), and a response screen (3 s). (**b**) ROIs that were used for all analyses pertaining to temporal order recognition (all analyses were performed relative to the control task baseline). (**c, e**) ROI results of the between-subject RSA (control analyses for the active and passive control groups are shown in Fig. S5). (**d, f**) Searchlight results of the between-subject RSA. Results were thresholded at *p* < 0.05, FWE-corrected at cluster-level, using a cluster-defining threshold of *p* < 0.001 (Tables S5-6). (**g, h**) ROI results of the trial-specific RSA pertaining to the (**g**) athlete study and (**h**) the training study (control analyses for the active and passive control groups are shown in Fig. S7). (**c, e, g, h**) Data points represent individual participants (athlete study, *n* = 33, memory training group, *n* = 17). Boxplots display the median and upper/lower quartiles, with whiskers extending to the most extreme data points within 1.5 interquartile ranges above/below the quartiles. The data points with error bars show the mean ± standard error of the mean (S.E.M.). Data distributions are based on the respective probability density function. All ROI-based results survived multiple comparison corrections using the false discovery rate (FDR) with the Benjamini-Hochberg procedure set to *q* < 0.05. \*\*\**p* < 0.001; ns, not significant. Pattern similarity reflects Fisher *z*-transformed Pearson correlations (*r*).

Contrary to what we had expected, results showed significantly increased neural pattern similarity between memory athletes compared to matched controls in the hippocampus (separate independent-sample *t*-tests, memory athlete, *n* = 17, matched controls, *n* = 16, *t*(31) = 6.430, *p* < 0.001, Cohen’s *d* = 2.240; Fig.4c) and in the precuneus during temporal order recognition (*t*(31) = 4.731, *p* < 0.001, Cohen’s *d* = 1.648; Fig.4c). Whole-brain searchlight analysis showed additional increases in neural pattern similarity in the parahippocampal-, fusiform-, and in the superior temporal gyrus, as well as in the angular gyrus (independent-sample *t*-test, same samples as above, contrast recognition > baseline, memory athletes > matched controls, *p* < 0.05, FWE-corrected, cluster-defining threshold *p* < 0.001, critical cluster size = 35 voxels, Fig. 4d, Table S5).

Similarly, we observed significantly increased neural pattern similarity between individuals of the memory training group after training (compared to the pre-training baseline) in the hippocampus (separate paired-sample *t*-tests, *n* = 17, *t*(16) = 4.842, *p* < 0.001, Cohen’s *d* = 1.174; Fig. 4e) and in the precuneus (*t*(16) = 4.852, *p* < 0.001, Cohen’s *d* = 1.177; Fig. 4e). This was not the case in the active or passive control groups (Fig. S5). Whole-brain searchlight analysis confirmed these results and further showed increased pattern similarity in the parahippocampal and fusiform gyrus, the caudate, and angular gyrus, among other regions (paired-sample *t*-test, *n* = 17, contrast recognition > baseline during post-training > pre-training session, *p* < 0.05, FWE-corrected, cluster-defining threshold *p* < 0.001, critical cluster size = 30 voxels, Fig. 4f, Table S6). These effects were larger in the memory training group compared to both the active and passive control groups (Fig. S6).

Given the relatively large size of the hippocampus ROI (2261 voxels), and given the known functional division along its longitudinal axis (Brunec et al., 2018; Collin et al., 2015; Poppenk et al., 2013), we further examined whether training-related effects were localized to specific subregions along the anterior-to-posterior hippocampal divisions (see Fig. S9b). This revealed increases in neural pattern similarity in the anterior hippocampus in both memory athletes and in the memory training group (post-training) (see Supplementary Results S1, Fig. S12, and Table S11).

We reasoned that these effects could be driven by sequence processing (which should be common to all experienced individuals as they mentally navigated along the loci routes to memorize the novel word-location associations) and performed a sequence-geometry analysis that tested for a monotonic decrease in neural pattern similarity as a function distance between word-location associations along the loci route (Supplementary Results S6, Fig. S20). This indeed showed that neural patterns in the (anterior) hippocampus were better explained by sequence geometry in the memory training group during temporal order recognition compared to passive (but not active) controls (post-training; note that this analysis was only performed using the training study data).

Still, the finding of increased neural pattern similarity in the medial temporal lobe during temporal order recognition (and partly also during word list encoding, as we found above) stands in contrast with the main result of unique neural (primarily prefrontal) representations associated with method of loci processing within and between experienced individuals. We thus repeated the trial-wise intersubject consistency (ISC) analysis during temporal order recognition, which capitalised on single-trial characteristics (memory athletes used their own loci routes during recall, which should be coupled to maximally unique neural representations and decreased neural pattern similarity between them; note that this analysis was only performed using the athlete study data). This showed increased trial-wise ISC in the precuneus between memory athletes compared to matched controls (Supplementary Results S6, Fig. S21).

In summary, we found shared neural representations in the anterior hippocampus and precuneus between memory athletes and between individuals of the memory training group (post-training) as they recognised the temporal order of previously studied material.

### Experienced individuals hold similar neural representations associated with different word triplets during temporal order recognition

Once again, we tested whether the finding of shared neural representations between experienced individuals was perhaps attributable to individuals holding similar neural representations associated with the different word triplets during temporal order recognition. We performed trial-specific RSA identical to the analysis of the word list encoding task, which yielded the average neural pattern similarity between all word triplets per participant and for each *a priori*-defined ROIs (hippocampus and precuneus), and which is complementary to the between-subject RSA (see section “Trial-specific RSA” in the Methods).

Findings showed significantly increased neural pattern similarity across word triplets in memory athletes compared to matched controls in the precuneus (separate independent-sample *t*-tests, memory athlete, *n* = 17, matched controls, *n* = 16, *t*(31) = 4.026, *p* < 0.001, Cohen’s *d* = 1.402; Fig. 4g), but not in the hippocampus during temporal order recognition (*t*(31) = 0.568, *p* = 0.574, Cohen’s *d* = 0.198; Fig. 4g). In a similar vein, data from the training study showed significantly increased neural pattern similarity across word triplets in the memory training group after training (compared to the pre-training baseline) in both the hippocampus (separate paired-sample *t*-tests, *n* = 17, *t*(16) = 4.372, *p* < 0.001, Cohen’s *d* = 1.060; Fig. 4h) and precuneus (*t*(16) = 4.890, *p* < 0.001, Cohen’s *d* = 1.186; Fig. 4h). This was not the case in the active or passive control groups (Fig. S7). Once again, we repeated the trial-specific RSA for the hippocampal anterior-to-posterior subdivisions. We found no significant increases in neural pattern similarity in any of the hippocampal subdivisions in memory athletes, but generally increased neural pattern similarity in the memory training group after training (see Supplementary Results S1, Fig. S13, Table S11).

Together, these results suggest that the shared neural representations in the hippocampus and precuneus between individuals who were proficient in the method of loci were driven by similar neural representations for the different word triplets during temporal order recognition.

### Shared neural representations between trainees and memory athletes during temporal order recognition

Thus far, we showed shared neural representations when individuals of the memory training group after training (the so-called “trainees”), or memory athletes, were judging the temporal order of previously studied words. We expected that these effects would translate to the between-group level, showing shared neural representations between trainees and memory athletes. This would indicate that the retrieval-related processes engaged in correctly judging the temporal order of the unique word-location associations are comparable between individuals who are proficient in the method of loci. We thus went on to quantify the degree of trainee-athlete similarity in neural representations by focusing on the *a priori*-defined ROIs (hippocampus and precuneus; Fig. 3a, and see section “trainee-athlete RSA” in the Methods).

Results showed significantly increased neural pattern similarity in the precuneus when trainees had completed the method of loci training (repeated measures ANOVA, *n* = 17, main effect of *time point*: *F* (1, 16) = 6.016, *p* = 0.026, *η_p_*^2^ = 0.27), and significantly higher neural pattern similarity between trainees × athletes compared to pattern similarity between trainees × matched controls (main effect of *group*: *F* (1, 16) = 9.775, *p* = 0.007, *η_p_*^2^ = 0.38). We also found a significant *time point* × *group* interaction (*F* (1, 16) = 92.430, *p* < 0.001, *η_p_*^2^ = 0.85), showing that neural pattern similarity between trainees × athletes (*p* = 0.002) but not between trainees × matched controls (*p* = 0.208) was higher in the post-compared to pre-training session (Fig.5a; results remained unchanged when removing two formal outliers in the pre-training session). Whole-brain searchlight analysis confirmed these findings (flexible factorial analysis with the factors *time point* and *group*, *n* = 17, *p* < 0.05, FWE-corrected, cluster-defining threshold *p* < 0.001, critical cluster size = 37 voxels, Fig. 5b, Table S7). Again, these effects were only present in the memory training group but not in the active or passive control groups (Fig. S8). An analysis of the anterior-to-posterior subdivisions along the hippocampal long-axis indicated a trend toward increased neural pattern similarity between trainees and memory athletes in the anterior hippocampus after training, whereas no clear effects emerged in the remaining subdivisions (see Supplementary Results S1, Fig. S14, Table S11).

Together, these results show that method of loci training led to shared neural representations between experienced individuals (i.e., trainees and memory athletes) during temporal order recognition.

### Shared neural representations during temporal order recognition are associated with better free recall performance after 4 months

In a final step, we tested the behavioural relevance of the shared neural representations between experienced individuals during temporal order recognition with respect to longer-term memory performance. As above, we employed a cross-validated machine-learning approach (Cohen, 2010; Supekar et al., 2013) to predict the increase in free recall performance due to method of loci training (i.e., the change in free recall performance from baseline to the 4-month re-test; Table S8) based on the changes in neural representations between all individuals of the training study × athletes from pre- to post-training during temporal order recognition (see section “Prediction analysis” in the Methods and trainee × athlete effects in Fig. 3a).

Results showed that increased neural pattern similarity between all individuals of the training study and athletes in the precuneus was significantly associated with better free recall performance after 4 months (*r*_(predicted, observed)_ = 0.44, *p* < 0.001, *n* = 44, Fig. 5c). We also repeated the analysis by defining the change in free recall performance after 24h, which gave virtually identical results (Table S9, but note that stratified within-group analyses did not reveal any significant associations for the ROI-based or whole-brain, voxel-wise regression analyses; see Supplementary Results S3, Table S12).

**Fig. 5.**
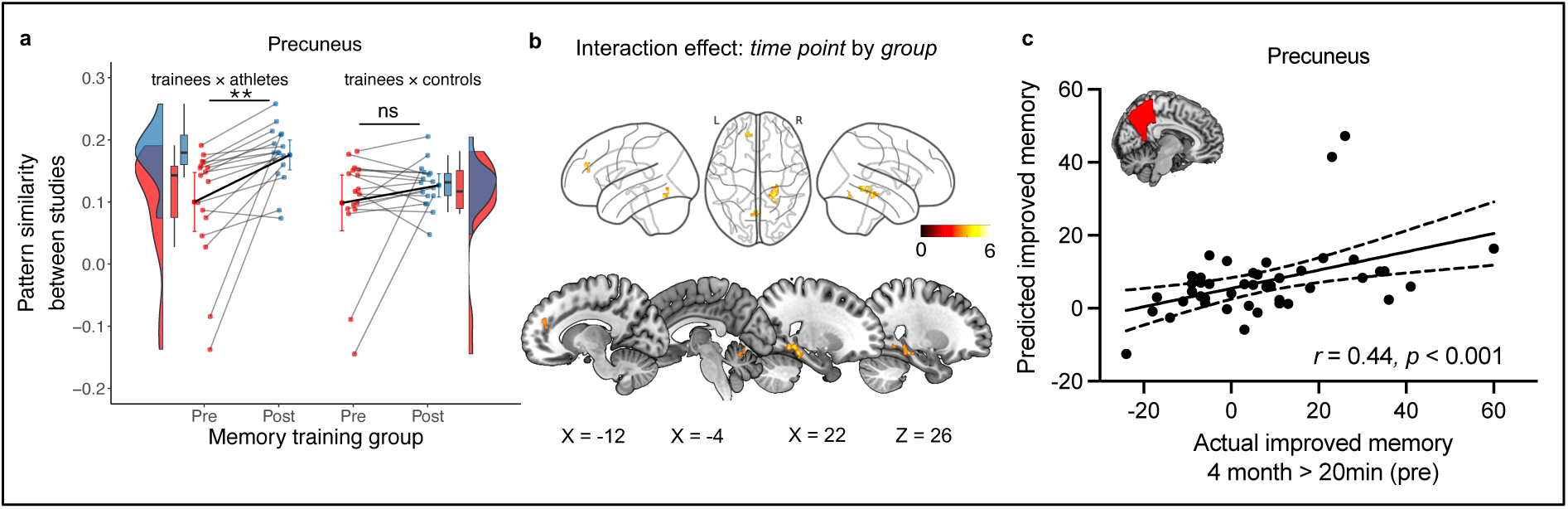
Method of loci training yields shared neural representations between trainees and memory athletes during temporal order recognition. (**a**) ROI results of the trainee-athlete RSA (control analyses for the active and passive control groups are shown in Fig. S8; analysis was performed relative to the control task baseline). Data points represent individual participants (memory training group, *n* = 17). Boxplots display the median and upper/lower quartiles, with whiskers extending to the most extreme data points within 1.5 interquartile ranges above/below the quartiles. The data points with error bars show the mean ± S.E.M. Data distributions are based on the respective probability density function. (**b**) Searchlight results for the time point × group interaction. Results were thresholded at *p* < 0.05, FWE-corrected at cluster-level, using a cluster-defining threshold of *p* < 0.001 (Table S7). (**c**). The scatter plot depicts the correlation between memory performance at the 4-months reset (4 months > immediate free recall test during the pre-training session) and the predicted changes in neural pattern similarity between all individuals of the training study and athletes from post- to pre-training (*n* = 44, see Methods). \*\**p*< 0.01; ns, not significant. Pattern similarity reflects Fisher *z*-transformed Pearson correlations (*r*).

In summary, we found shared, retrieval-related neural representations between all individuals of the training study that predicted a longer-lasting benefit on memory performance. These associations were not specific to the memory training group, which does not surprise given that recognition performance was generally less affected by method of loci training (while performance in the memory training group was generally higher compared to controls, it did not reach the levels of memory athletes with decades of experience; see Supplementary Results S5).

## Discussion

We investigated the impact of a 6-week-long method of loci training on the neural representations that support superior memory. To characterize whether the effects of such multiple-week training would be comparable to existing expertise with mnemonic techniques, we performed two separate studies that involved initially mnemonics-naïve individuals who completed a method of loci training, and experienced memory athletes. Several key results emerged from our novel RSA analyses that substantially expand our previous work (Dresler et al., 2017; Wagner et al., 2021): We found that method of loci training was associated with distinct neural representations primarily in the prefrontal cortex between experienced individuals as they studied lists of novel words. Put differently, the memory training group (post-training) and memory athletes appeared less similar in their encoding-related neural representations. This was also reflected at the level of the studied material, showing that individuals formed distinct neural representations for each word-location association. Focusing on the trainee-athlete similarity during encoding, we showed that more distinct neural representations exerted a lasting benefit on memory as they predicted free recall performance after 4 months. This may be related to the formation of unique word-location associations in an individual’s “memory palace”. We also took the opportunity to explore the neural representations during temporal order recognition. There, memory training led to shared (rather than distinct) neural representations in the hippocampus and precuneus between experienced individuals, suggesting a more generalised role of these regions in episodic memory. Taken together, our findings provide first insights into how extended memory training transforms the neocortical representations that enable effective, superior memory.

Our focus revolved around the potential impact of method of loci training on the encoding-related neural representations in individuals of the memory training group after they completed their training, as well as in memory athletes. Using a between-subject analysis, we found distinct neural representations between these experienced individuals in the left lateral prefrontal cortex and superior frontal gyrus during word list encoding. This was mirrored at the level of the studied material (using a within-subject, between-item analysis, complementary to the abovementioned between-subject RSA), revealing that they held unique neural representations for the specific word-location associations. In line with these results, Liu and colleagues (2022) reported that 5-day method of loci training was associated with lower neural pattern similarity in hippocampal subfields during memory encoding (thus, distinct hippocampal patterns). This effect was specific to the studied material that was mentally placed at close-by locations along the imagined route and was coupled to accurate temporal order memory. Bellmund and colleagues (2022) investigated neural pattern similarity during the encoding of event sequences (but their study did not involve memory training). The authors observed distinct hippocampal patterns when events appeared closer in time. More evidence comes from LaRocque and co-authors (2013), who reported that successful recognition memory was predicted by distinct hippocampal representations at encoding (but note that we could not confirm these hippocampal effects; see Supplementary Results S2 along with a brief discussion). Here, we found that memory training led to distinct neocortical representations in lateral and superior prefrontal regions. The lateral prefrontal cortex is known as a key player when it comes to reducing memory interference during learning (Ghazizadeh et al., 2018; Sakai et al., 2002). Using electrophysiological recordings in non-human primates, Ghazizadeh and colleagues (2018) showed that lateral prefrontal neurons coded for object-reward associations that persisted over several months and that were resistant to extinction. Adjacent inferior prefrontal regions were shown to support the retrieval of detailed episodic information (Badre & Wagner, 2007; Öztekin et al., 2009) and to differentiate (temporal) contexts in both humans (Sakai et al., 2002; Schlichting et al., 2015) and non-human primates (Mante et al., 2013). Prefrontal regions may support the longer-term stabilisation of memories via top-down projections (Blumenfeld & Ranganath, 2007), resonating with theoretical accounts of pattern separation that highlight the role of distinct neural representations in orthogonalizing overlapping memory content and minimising interference (Amer & Davachi, 2023; Nash et al., 2021; Stevenson et al., 2020; Yassa & Stark, 2011).

Strategy customisation might be a relevant factor when it comes to prefrontal cortex engagement. Experienced individuals were accustomed to forming unique word-location associations during encoding and, depending on the characteristics of the to-be-encoded word, individuals may have flexibly modulated their mnemonic strategy (e.g., relying on vivid, visual imagery only or mentally interacting with the items and/or loci, such as moving heavy boxes out of the way when the item “box” needed to be remembered). Some participants might have also segmented the mental route by creating event boundaries between adjacent associations to differentiate contexts (Amer & Davachi, 2023; Nash et al., 2021; Stevenson et al., 2020; Yassa & Stark, 2011). This shows that utilising the method of loci is tied to highly idiosyncratic mental imagery, amplifying memory “uniqueness” during encoding by reducing the spatiotemporal similarity between events. As such, it is applicable to any other instance that requires memorising large amounts of (arbitrary) information, knowledge about the order of items (e.g., numbers, shopping lists), or study material (e.g., law texts, anatomical nomenclature; but note that it generalises poorly to other tasks that do not tap into this domain; Ramon et al., 2016).

Mentally placing to-be-remembered information at salient locations along the imagined path likely produces relatively bizarre associations, thereby triggering neural mechanisms related to novelty (Fernández & Morris, 2018). This is known to crank up dopamine and noradrenaline release from the brainstem and ventral striatum, which boosts memory persistence (Duszkiewicz et al., 2019). In addition, the lateral prefrontal cortex appears to be associated with the processing of novel information as well (Ranganath & Rainer, 2003). Prior work using invasive recordings in non-human primates showed that novel stimuli increase the activity of neurons in the lateral prefrontal cortex (Matsumoto et al., 2007), and lesion studies highlighted that damage to the lateral prefrontal cortex abolishes the memory advantage for novel compared to non-novel stimuli in both non-human primates (Parker et al., 1998) and human patients (Kishiyama et al., 2009). Here, we found distinct neural representations in the lateral prefrontal cortex between experienced individuals during encoding, which might be related to the processing of novel word-location associations. Our results further included the superior frontal gyrus. Human patients with lesions in this area exhibit marked deficits in working memory and spatially oriented attention (Boisgueheneuc et al., 2006). Other reports discussed this region’s role in monitoring and executive processing (Owen, 2000; Petrides, 2000), as well as its part within a prefrontal hierarchy that processes multi-feature and stimulus-response associations along an inferior-to-superior gradient (Koechlin et al., 2003). Taken together, we speculate that the prefrontal involvement during word list encoding might stem from the formation of complex, multi-feature associations (lateral prefrontal cortex) that require monitoring and spatial attention (superior frontal gyrus). Since the memory training group and memory athletes received extensive training in utilising the method of loci, the process of associative memory formation might become comparable to stimulus-response learning (Müller et al., 2018), allowing for the rapid integration of novel information into the well-established “memory palace”.

We also demonstrated distinct neural representations between experienced individuals in posterior representational regions during encoding, including the left inferior temporal cortex and bilateral posterior parietal areas. The inferior temporal cortex was shown to code for object categories during perception (Op de Beeck et al., 2008) and mental imagery (Dijkstra et al., 2019). Previous work that employed invasive recordings in monkeys and fMRI-based recordings in humans demonstrated that the representational geometry in the inferior temporal cortex (e.g., how distinct the neural representations of two objects are) depends on an object’s visual features (Kriegeskorte, Mur, Ruff, et al., 2008) and is highly specific to an individual’s perceptual judgments (Charest et al., 2014). Our results also extend toward the (left) visual word form area and to posterior parietal regions, which are known to selectively process written letter strings (Dehaene & Cohen, 2011) and to code for (egocentric) spatial context (Baddeley et al., 2001; Sereno et al., 2001; Takashima et al., 2007), respectively. Naturally, these effects are likely specific to the stimulus material at hand and might be different for other sensory modalities and task setups. One limitation is that we lack information on the specific mental routes that participants chose during the memory tasks in the post-training MRI session, their specific strategies, or information on the vividness of their imagination. This would have allowed us to decode the reinstatement of, e.g., landmarks during temporal order recognition, elucidating how vivid imagination contributes to training-related changes in neural pattern similarity. We encourage future research to address this with an adapted task design that better lends itself to such a decoding-based approach (e.g., Wagner et al. 2015, 2016, as well as Haynes, 2015).

Our findings partly stand in contrast to previous reports that highlighted the beneficial effect of interindividual similarity on cognitive performance. For instance, Sheng and colleagues (2023) showed that the degree of neural pattern similarity to an average, group-based neural pattern during associative encoding predicted subsequent memory ability across individuals (the more individuals resembled the group in their neural representations, the better their memory was). Such “neural alignment” might stem from the processing of similar content, and it follows logically that the processing of unique memories should be tied to distinct neural representations. Meshulam and colleagues (2021) not only showed neural alignment between learners after acquiring new conceptual knowledge but also reported neural alignment between learners and experts that was predictive of learning outcomes. The authors interpreted their effects in terms of group knowledge, or an ideal canonical representation, comparable to memories for shared experiences (Chen et al., 2017). Here, we also found that trainees and memory athletes were less similar in their encoding-related prefrontal representations than individuals who did not undergo memory training (but note that neural representations in the hippocampus were more similar; see the Supplementary Materials, where we also provide an extended discussion). While the acquisition of a shared representation might certainly be useful when conceptual knowledge structures are acquired, we suggest that distinct neural representations may be favourable when it comes to memory specificity. For instance, unique memories were shown to be retrieved with increased visual detail compared to repeated events or facts (Tanguay et al., 2023). The distinct trainee-athlete neural representations also predicted free recall performance after 4 months, emphasising the beneficial impact of memory “uniqueness” on memory persistence (Burgess et al., 2002; Ranganath & Ritchey, 2012a).

Following the word list encoding task, participants completed a temporal order recognition task that is commonly used during memory championships. Results showed that memory training led to shared (rather than distinct) neural representations between experienced individuals in the hippocampus, adjacent medial temporal lobe, posterior medial regions, and in the angular gyrus. Methodologically, a contributing factor might be the task design. Temporal order judgments were based on word triplets (not single words) that required participants to mentally retrieve (i.e., navigate between) three word-loci associations and to evaluate their order (thus, the analysis boosted common aspects between them). Conceptually, the task should have facilitated mental route reinstatement (and thus, shared cognitive processes), resulting in increased between-subject neural pattern similarity (note that we also analysed reinstatement-related processes, see Supplementary Results S4, along with a brief discussion). In line with this, we found that neural patterns in the (anterior) hippocampus were explained by sequence order (see Supplementary Results S6), which was previously linked to hippocampal-entorhinal circuits that encode abstract relational and temporal information (Constantinescu et al., 2016; Eichenbaum, 2013; Garvert et al., 2017; Hsieh et al., 2014; Schapiro et al., 2013). We also analysed whether word-location associations were uniquely represented in individuals (or whether they were similar between them). Results showed increased trial-wise neural pattern similarity in the precuneus between memory athletes compared to matched controls (see Supplementary Results S6), fitting with theoretical accounts of scene construction and context reinstatement in this brain region (Diana et al., 2007; Ranganath & Ritchey, 2012b). Together, these data highlight a potentially more generalised role of the (anterior) hippocampus and precuneus in episodic memory during method of loci processing.

On a broader scale, associative retrieval using the method of loci is comparable to the retrieval of prior knowledge structures, or schemas (Kesteren et al., 2012), that are associated with neural alignment of retrieval-related brain activity between individuals (Chen et al., 2017). Likewise, associative memory retrieval (such as is the case in our data set) engaged similar levels of neural activity and connectivity between experienced individuals (Dresler et al., 2017; Wagner et al., 2021). Regarding the specific set of brain regions, the hippocampus is involved in comparing incoming information with prior knowledge (Hindy et al., 2016; Horner et al., 2015), while posterior medial regions are known to be engaged in self-referential (Lyu et al., 2023) and episodic memory processes (Sestieri et al., 2017), and in the recall of temporal sequences. Furthermore, Xiao and co-authors (2020) reported shared neural representations between individuals during associate retrieval in the angular gyrus. We do not think that the shared neural representations during temporal order recognition (and partly also during encoding) are tied to common encoding traces (i.e., the retrieval of the same word-location associations between individuals), as participants could freely use which and how many of the trained mental routes they would utilise, and because the order of words was random. Increased neural pattern similarity within this ensemble of hippocampal and posterior regions may thus be tied to shared processes that facilitate the vivid retrieval of the “memory palace” when evaluating temporal order. However, we fully acknowledge that our additional analyses cannot disentangle the precise underlying mechanisms of method of loci processing (also keeping in mind the methodological limitations of fMRI-based research; Ramsey et al., 2010). Future work could build on our findings by employing methods with higher temporal resolution or direct neural recordings to further characterise the neural mechanisms of mental reinstatement during method of loci processing.

In conclusion, we provide the first evidence of how multiple-week memory training transforms the neural signatures that underlie superior memory. We found distinct neural representations between memory athletes and between individuals who were trained to utilise the method of loci over several weeks (the so-called “trainees”) while they studied novel material. The distinct neural representations were also evident between trainees and memory athletes, and more distinct representations predicted better memory performance after 4 months. We suggest that these effects are linked to pattern separation and novelty processes that bolster individual memory “uniqueness”. In parallel, the data revealed increased neural pattern similarity in the anterior hippocampus and precuneus, suggesting a more generalised role of these regions in episodic memory formation and retrieval. Collectively, we suggest that method of loci training serves to disambiguate memory content upon encoding, creating unique prefrontal memory engrams that are tied to one’s “memory palace”, subserving extraordinary memory ability.

## Acknowledgments

We thank S. Weisig and P. Schuster for their help with data collection and the International Association of Memory (IAM; www.iam-memory.org) for support in recruiting memory athletes. This article is based on work from a Veni project supported by the Dutch Research Council (NWO); and the European Platform for Life Sciences, Mind Sciences, and the Humanities supported by the Volkswagen Foundation. This research was funded in part by the Austrian Science Fund (FWF, 10.55776/P34775), awarded to I.C.W. For open access purposes, the author has applied a CC BY public copyright license to any author accepted manuscript version arising from this submission.

## Author contributions

M.D., B.N.K. and M.C. proposed the original study design. B.N.K. and M.D. acquired the original data. J.R., I.C.W. and M.D. conceived the specific research question at hand. J.R. and I.C.W. contributed to methods development. J.R. and I.C.W. analysed the data. J.R., I.C.W., Y.Z., F.L. and M.D. contributed to the interpretation of the results. J.R. and I.C.W. drafted the paper. All authors contributed to the critical revision of the paper.

## Methods

### Participants of the athlete and training studies

In the athlete study, we enrolled 23 memory athletes (mean age ± s.d., 28 ± 8.6 years, 9 females) who were among the top 50 participants in the World Memory Championships at the time of study participation (www.world-memory-statistics.com). Of these 23 athletes, 17 took part in a word list encoding and temporal order recognition task inside the MRI scanner. Memory athletes were matched to a control group that was selected from gifted students of academic foundations and members of the high-IQ society “Mensa” (https://www.mensa.org/). Matching criteria included IQ, age, sex, handedness, and smoking status. Six participants of the matched control group were selected from the training study based on their cognitive abilities, as determined during the screening session (details of the screening session are provided below). They were evenly sampled from the three training groups. All participants of the matched control group underwent a standardized memory test (Bäumler, 1974) to prevent the inclusion of “natural” superior memorizers (none of the participants met this criterion). Additionally, they completed a test for fluid reasoning (Weiß, 2006). Any prior experience with systematic memory training was an exclusion criterion. Together, these participant groups were part of the so-called “athlete study” (memory athletes: *n* = 17, mean age ± s.d., 24.6 ± 4.3 years, 8 females; matched controls: one participant was excluded due to excessive head motion during fMRI, *n* = 16, mean age ± s.d., 25.4 ± 3.9 years, 7 females).

To assess the behavioural and neural effects of mnemonic training in participants without prior experience with the method of loci, we enrolled 51 healthy, right-handed individuals (mean age ± s.d., 24 ± 3 years) from the University of Munich. This constituted the training study. We only included male participants due to the known influence of the menstrual cycle on memory (Genzel et al., 2012; Sattari et al., 2017) and hippocampal morphology (Zsido et al., 2023) and because the longitudinal study design would not have allowed for a systematic control of this factor. None of the participants had any reported experience with memory training, and none exhibited superior memory abilities. Participants were assigned to one of the three groups based on their cognitive performance as determined during the initial screening session (using assessments from Bäumler, 1974; Weiß, 2006). This allowed us to ensure that any potential changes in memory performance would be attributable to the method of loci training (see Table S1). The memory training group underwent a 6-week training in the method of loci (*n* = 17, mean age ± s.d., 24 ± 3 years). These participants were directly compared to a group that completed a working memory training (active controls: one participant was excluded because of technical issues with the MR recordings, *n* = 16, mean age ± s.d., 24 ± 3 years) and to a group that did not undergo any intervention (passive controls: n = 17, mean age ± s.d., 24 ± 4 years).

All participants were native German speakers with normal or corrected-to-normal vision and without a history of psychiatric or neurological diseases. Written informed consent was obtained before study enrolment, and the study protocol was reviewed and approved by the ethics committee of the Medical Faculty of the University of Munich, Germany.

### Study overview and cognitive training

Participants of the athlete study completed a single MRI session (Fig. 1a). Participants of the training study completed two identical MRI sessions that were separated by a 6-week interval, as well as a behavioural re-test four months later (Fig. 1b).

#### Method of loci training

The memory training group completed systematic memory training using the method of loci across 6 weeks. Participants were introduced to the method of loci at the Max Planck Institute of Psychiatry (Munich, Germany). They were guided in constructing their first loci route and received supervision while using the route in an initial memory task. This basic route was the same for all participants and included 50 locations in and around the Max Planck Institute (e.g., blackboard, balcony, fire extinguisher, staircase, lecture hall entrance, elevator, printer, exit, tree, pond, rock, parking space, house, street, hedge, and so forth). Subsequently, participants were acquainted with the online platform that was utilized for home-based training, for monitoring training progress, and compliance (https://memocamp.com). They were also provided with a training plan for the upcoming week.

After that, participants underwent a 40-day home-based training (30 minutes per day). The training plan included instructions for selecting locations to reduce interference from previously memorized word lists and to ensure an equal distribution of training time across different routes. During the first two weeks of training, participants were instructed to individually construct and memorize three additional loci routes according to the instructions. They were motivated to incorporate locations from their private environment as well. This resulted in a total of four routes, including the initially taught route. These routes were employed for memorizing random words. The task difficulty, which was determined by the number of words to be memorized, was adjusted dynamically based on individual performance. For example, at the beginning of each daily training session, participants were presented with five words during the first run. If participants successfully recalled all the words in any given run, the number of presented words increased by +5 in subsequent runs. The speed of training success was defined as the average number of runs needed to progress to the next difficulty level, and 40 words needed to be recalled successfully (equivalent to eight runs). While most participants in the memory training group reached this final level (16 out of 17), it is hardly achievable for mnemonics-naïve participants.

Log files from the daily training sessions were monitored daily to ensure participants’ compliance. In cases where a participant missed a training session or did not complete their daily routine, they were contacted the following morning. They then received instructions to increase their training time on the subsequent day to compensate for the missed session. Additionally, participants were required to visit the laboratory once a week for supervised training in small groups of 2-3 participants. During these sessions, they were interviewed about their training regime and received a training plan for the upcoming week.

#### Working memory training and passive controls

Participants of the active control group were introduced to the dual *n*-back task that required them to monitor and update a series of visually presented locations and auditorily presented letters (Jaeggi et al., 2008). The value of n changed between sets of trials, continuously adjusting based on performance. This adaptive process resulted in the task demand changing according to each participant’s individual performance. They engaged in a 40-day home-based training (30 minutes per day, using an online working memory training program). To ensure compliance, their progress was monitored daily. If participants missed or did not complete their daily training routine, they were notified the following morning, with instructions to increase their training time on the subsequent day to compensate for the missed session. Additionally, participants visited the laboratory once a week in small groups of 2-3 participants for an interview regarding any potential training issues and received training under direct supervision. Participants were instructed to perform to the best of their ability but were advised to avoid any systematic long-term memory training. The passive control group did not receive any training between the two sessions and were given no specific instructions beyond refraining from systematic memory training while being enrolled in the study.

### General structure of MRI sessions

An MRI session of both the athlete and training studies, during pre- and post-training, began with the acquisition of a structural brain image, followed by a (baseline) resting-state fMRI period, the word list encoding and temporal order recognition tasks (as shown in Fig. 1c), and a post-task resting-state fMRI (resting-state periods are not further discussed in this paper). The MRI sequence was stopped and started after each task/resting-state period. Subsequently, participants completed a free recall test in the behavioural laboratory 20 minutes after leaving the MR scanner (i.e., the immediate free recall test), and another free recall test via a phone interview 24 hours later (i.e., the delayed free recall test). Participants of the athlete study only performed the immediate free recall test, but not the delayed free recall test.

#### Word list encoding task

We utilized this task (along with the temporal order recognition task described below), because we reasoned that the particular strength of the method of loci lies in learning and recalling ordered sequences, facilitated by the mental navigation through an imagined “memory palace”. The task involved concrete nouns that were divided into two separate lists (1× during pre-training MRI session, 1× during post-training MRI session) that each contained 72 concrete nouns (e.g., table, hammer, chair, cup; see Table S14 for an overview of all used words). The words of each list were presented in random order during MRI, such that the order of words was shuffled per participant and session. The order of the word lists (i.e., which one was used pre-/post-training) was counterbalanced across individuals.

A trial corresponded to the presentation of a single word. The 72 trials were organised into 12 blocks of 6 words each. A block began with an instruction (5 seconds), followed by the 6 words (each presented for 3 seconds), separated by a jittered fixation cross of 2-5 seconds (mean = 3.5 seconds). After each block, a fixation cross was displayed for 25 seconds, which was a deliberate choice to space the learning and to allow for the blood-oxygen level dependent (BOLD) signal to return to baseline (Buuren et al., 2014; Wagner et al., 2016). Participants were explicitly instructed not to rehearse the studied material during these fixation periods but to keep their thoughts unoccupied. The total duration of the task was 10 minutes. All stimulus material was presented with Presentation (Neurobehavioral Systems, Albany, CA, USA).

During the tasks, participants were presented only with words (word list encoding task) or word triplets (temporal order recognition task; thus, no information or visual cues were given regarding the mental routes). Memory athletes and the memory training group (post-training) were instructed to employ the method of loci to memorise the word lists. Specifically, they were asked to mentally navigate through their “memory palace” and place each word at a specific location. Participants were free to choose which route and how many they would use to anchor the 72 items. Memory athletes used their own, pre-established routes.

#### Temporal order recognition task

We developed this task to form an MR-compatible measure of recall performance. Participants viewed 24 triplets of words based on material from the previously encoded word list (using adjacent words that were shuffled within triplets). Hence, each word triplet consisted of three words that were previously presented in direct (0-distance) and close (1-distance) proximity.

A trial commenced with a cue signalling the upcoming word presentation (3 seconds), followed by the presentation of a word triplet (9 seconds). Subsequently, participants were asked to determine whether the word order in the triplet was the same as during the previously encoded word list. They indicated their choice with a button press (answer options “same, sure”, “same, maybe”, “different, maybe”, and “different, sure”; 3 seconds). Triplet presentations were interleaved with a control task during which participants were asked to determine whether novel word triplets were displayed in ascending or descending order based on the number of syllables in each word (using the same timing and response options as described above). The word order recognition trials (A) and the control task (B) were presented in an ABAB fashion. The total duration of the task was 10 minutes. Memory athletes and the memory training group (post-training) were instructed to use the method of loci to judge whether the presented triplets were presented in the correct order. In other words, participants were asked to mentally navigate through their “memory palace”, retrieving individual words to assess whether they were presented in the correct order. No specific instructions were given to the other groups.

#### Free recall

The free recall test served as our primary outcome measure of memory performance, as it closely resembles tasks used in memory championships. Following MR scanning (approximately 20 minutes later), participants were instructed to freely recall and write down the 72 concrete words they had learned during the preceding word list encoding task (immediate free recall test). After 5 minutes, participants were asked if they needed more time, and if so, the free recall was terminated after an additional 5 minutes. The delayed free recall test was conducted 24 hours later via phone, following the same 5+5-minute format. Memory performance was assessed by counting the number of words correctly recalled, ignoring word order and spelling mistakes. The delayed free recall test was announced to all participants, as our intention was to maintain consistency between pre- and post-training sessions. Notably, participants in the athlete study completed only the immediate free recall test and not the delayed version.

#### Re-test after four months

Participants in the training study also completed a behavioural re-test four months after the post-training session. During this re-test, we performed the word list encoding task, followed by a delay period that included a reasoning task (15 minutes) and a free recall task (5+5 minutes, serving as our primary outcome measure). The task material during the word list encoding task was identical to that used during the initial pre-training session because the 4-month re-test was added after the word lists had been constructed. Importantly, participants were assigned to the different training groups only after the pre-training session. As a result, any material from the first session that might have been remembered during the 4-month re-test should be unaffected by training. As mentioned earlier, participants in the memory training group (post-training) were instructed to use the method of loci during word list encoding. Five participants (2 from the memory training group, 2 from the active control group, and 1 from the passive control group) were not available for the re-test.

### Analysis of free recall performance

To quantify memory performance, we calculated the change in freely recalled words from pre- to post-training sessions (post-minus pre-training). This was done separately for the immediate free recall test (20 min post-minus pre-training), the delayed free recall test (24 hours post-minus pre-training), as well as for the re-test after four months (4-month re-test minus pre-training test 20 min). Memory athletes only completed the immediate free recall test (20 min post-MRI), and their performance was quantified as the number of recalled words.

### MRI data acquisition parameters

All imaging data were collected at the Max Planck Institute of Psychiatry (Munich, Germany) using a 3T scanner (GE Discovery MR750, General Electric, USA) with a 12-channel head coil. We obtained a time series of 292 T2*-weighted BOLD images with the following EPI sequence: TR, 2.5 s; TE, 30 ms; flip angle, 90°; 42 ascending axial slices; FOV, 240 × 240 mm; 64 × 64 matrix; slice thickness, 2 mm, slice gap, 0.5 mm. The structural image was acquired with the following parameters: TR, 7.1 ms; TE, 2.2 ms; flip angle, 12°; in-plane FOV, 240 mm; 320 × 320 × 128 matrix; slice thickness, 1.3 mm.

### MRI data preprocessing

The MRI data were processed using Statistical Parametric Mapping (SPM version 12; www.fil.ion.ucl.ac.uk/spm/) in combination with Matlab (The Mathworks, Natick, MA, USA). The first four functional volumes were discarded to allow for T1 equilibrium. The remaining images were corrected for acquisition delay and were realigned to the mean functional image of each session. Processing was done separately for the word list encoding and temporal order recognition tasks. The high-resolution structural image was first segmented into grey matter, white matter, and cerebrospinal fluid, generating a deformation field for spatial normalisation. The functional scans were then co-registered to the structural image to ensure alignment. This deformation field, derived from T1 segmentation, was subsequently applied to normalise both the structural and functional images to the standard stereotactic Montreal Neurological Institute (MNI) space. All images were resampled to a 2-mm isotropic voxel resolution.

### Modelling of fMRI data

#### Conventional modelling across trials

To track the alignment of neural representations within and between groups, we employed two separate General Linear Models (GLMs) that captured BOLD activity changes during the word list encoding and temporal order recognition tasks. Using data from the word list encoding task, the BOLD response to all trials was modelled with a single task regressor, time-locked to the onset of each trial (boxcar function with a duration of 3 seconds). Instructions were collapsed into a regressor-of-no-interest (boxcar function with a duration of 5 seconds). Both the inter-trial intervals (ITIs, 2-5 s) and the 25 s fixation cross period after each 6th word were treated as implicit baseline (therefore, contrasts against baseline reflected neural activity changes during word list encoding relative to the implicit fixation cross baseline). Regressors were convolved with the canonical hemodynamic response function (HRF), and the six realignment parameters were included in the model to account for rigid-body head motion. This resulted in a single beta image (capturing average BOLD activity changes during word list encoding) per participant and session.

Using data from the temporal order recognition task, we distinguished between old (recognition trials) versus novel (control trials) words in our GLM. We defined a regressor that included all word triplets (with words that were studied during the previous word list encoding task), time-locked to the onset of each trial and modelled as a boxcar function that lasted until a button press occurred (thus, collapsing across the triplet presentations and response periods). We merged these periods because they were not separated by an inter-trial interval (ITI), making it difficult to dissociate their BOLD responses (Friston et al., 1995), and because we reasoned that participants would mentally reinstate the loci route and hold this mental representation active until providing a response. Instructions were incorporated into a regressor-of-no-interest (boxcar function with a duration of 3 seconds). Word triplets and response periods from the control task (syllable counting of novel words) were not explicitly modelled and thus formed the implicit baseline (therefore, contrasts against baseline reflected neural activity changes during temporal order recognition relative to the implicit control task baseline). The remaining steps were carried out identically to the above. This resulted in a single beta image (capturing average BOLD activity changes during temporal order recognition) per participant and session. For participants of the training study, separate GLM models were defined for the pre- and post-training sessions.

#### Trial-specific modelling

To track the alignment of neural representations across single words or word triplets, we employed two separate GLMs that captured trial-specific BOLD activity changes during the word list encoding and temporal order recognition tasks. That is, using data from the word list encoding task, each presented word was modelled as a separate regressor (following Mumford et al., 2014), time-locked to the onset of each word (boxcar function with a duration of 3 seconds). Instructions were collapsed into a regressor-of-no-interest (boxcar function with a duration of 5 seconds). All regressors were convolved with the canonical HRF, and the six realignment parameters were included to account for head motion. This resulted in 72 trial-specific beta images (capturing trial-specific BOLD activity changes during word list encoding) per participant and session. To track the alignment of neural representations across word triplets during the temporal order recognition task, each presented word triplet was modelled as a separate regressor, time-locked to trial onset (boxcar function, duration modelled until a button press occurred). Instructions were grouped into a separate regressor-of-no-interest (boxcar function with a duration of 3 seconds, all remaining details identical to above). This resulted in 24 beta images (capturing trial-specific BOLD activity changes during temporal order recognition) per participant and session. As mentioned above, pre- and post-training sessions were modelled separately for participants of the training study.

### Definition of regions-of-interest (ROIs)

We selected four regions-of-interest (ROIs) based on our previous findings of training-related changes in neural activity (Wagner et al., 2021). All ROIs were anatomically defined using the Brainnetome Atlas (https://atlas.brainnetome.org/). For the investigation of changes in neural representations during the word list encoding task, we selected the left superior frontal gyrus (anatomical, left, A6m, medial area 6, 758 voxels) and the left lateral prefrontal cortex (anatomical, left, inferior frontal gyrus, 2161 voxels). For the investigation of changes in neural representations during the temporal order recognition task, we focused on the bilateral hippocampus (2261 voxels) and precuneus (5843 voxels). We included all voxels within the anatomical mask.

### Representational Similarity Analysis (RSA)

We employed representational similarity analysis (RSA) to assess the impact of method of loci training on neural representations (Fig.1d). Specifically, we examined (i) the degree of neural alignment between individuals of the same experimental group (i.e., between-subject RSA), (ii) across the studied material (i.e., trial-specific RSA), and (iii) the degree of neural alignment between trainees and memory athletes (i.e., trainee-athlete RSA).

#### Between-subject RSA: degree of neural alignment between individuals of the same group

This analysis was based on the beta-maps obtained from conventional across-trial modelling (thus, capturing average BOLD activity changes during word list encoding and temporal order recognition). For each participant, neural activity was extracted from the pre-defined ROIs and the data were reshaped into a pattern vector for each ROI. The data were then *z*-scored across voxels to emphasise the multivoxel pattern structure rather than the absolute activation values (Anderson et al., 2016; Walther et al., 2016a). The pattern vectors of each ROI and each participant were then correlated to the pattern vectors of all other participants of the same group, using pairwise Pearson’s correlation (*r*). The resulting correlations were Fisher’s *z*-transformed (using [*z* = 0.5*log(1+*r*)/(1-*r*)]) to stabilise variance and approximate a normal distribution (Fisher, 1915; Walther et al., 2016b). Between-subject neural pattern similarity for each subject was then computed as the average pattern similarity of each participant to all other participants within the same group (that is, the average correlation of voxel patterns from participant 1 with those of participants 2∼*N* of a given group, Fig 2a). This procedure was repeated for both the pre- and post-training sessions, and separately for the word list encoding and temporal order recognition tasks, as well as separately for the different groups in the athlete and training studies. The following equations were used to quantify the between-subject pattern similarity of individuals in the different experimental groups of each study. In brief, we calculated the pattern similarity of each participant (here denoted as sub_1_; to provide an example, we chose a participant of the memory training group, LOC) to the group average of that same group during the pre-(Equation 1) and post-training (Equation 2) sessions, for example, during word list encoding. Equations 3 and 4 follow the same rationale to assess the pattern similarity of a different sub_1_ in the matched control (CON) and the memory athlete groups (ATH) to their respective group averages. In these equations, *n* denotes the total number of participants in the respective groups, and j represents the participant index for participants in athletes or matched control groups other than subject 1, starting from 2 to ensure the exclusion of self-comparison (For details on statistical analysis, see the section “Statistical analyses and thresholding”).

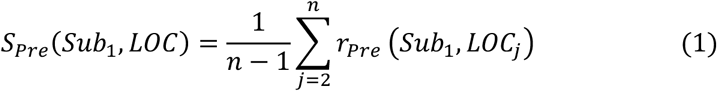

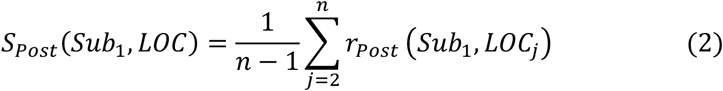

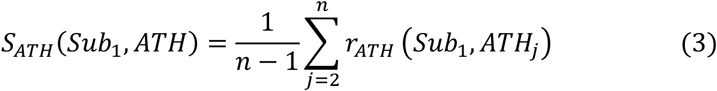

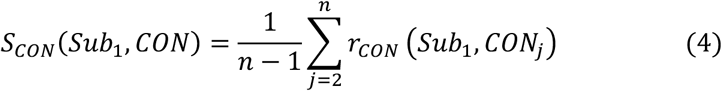

The between-subject RSA examined the degree to which an individual’s neural activity pattern during word list encoding (or temporal order recognition) aligned with the group-averaged neural representation (thus, an individual of the memory training group was compared to the group-averaged neural representation of the remaining memory training group). This approach captured neural alignment between participants of the same group.

#### Trial-specific RSA: degree of neural alignment across studied material

To investigate whether method of loci training was associated with changes in the neural representation of the studied material, we repeated the abovementioned between-subject RSA procedure but instead based the analysis on the beta-maps that were obtained from the trial-specific modelling (thus, capturing trial-specific BOLD activity changes during word list encoding and temporal order recognition that pertained to the single words and word triplets, respectively). Again, voxel activity patterns were extracted for the pre-defined ROIs for all trials, and data were then *z*-scored across voxels. The pattern vectors of each ROI and each trial were then correlated to the pattern vectors of all other trials of the same participant, using pairwise Pearson’s correlation (*r*). The resulting correlations were Fisher’s *z*-transformed. Trial-specific neural representation for each subject was computed as the average pattern similarities of each trial with all other trials (see Fig. 2h). This procedure was repeated for both the pre- and post-training sessions, and separately for the word list encoding and temporal order recognition tasks, as well as separately for the different groups in the athlete and training studies. The following statistics are identical to the above (For details on statistical analysis, see the section “Statistical analyses and thresholding”).

The trial-specific RSA focused on the neural pattern similarity between different words during the word list encoding task (or between word triplets during the temporal order recognition task) and was conducted on a per-subject basis. This approach captured the neural alignment between single words (or word triplets), assessing whether method of loci training led to shared or distinct neural representations for each item. Theoretically, this approach is linked to processes such as pattern separation and pattern completion, which support episodic memory by minimising interference between overlapping events or by reinstating cued memory content, respectively (Amer & Davachi, 2023; Yassa & Stark, 2011). The between-item RSA is hence orthogonal to the between-subject RSA, with both approaches yielding complementary results on the level of single items (per subject) and the neural alignment between subjects.

#### Trainee-athlete RSA: degree of neural alignment between groups

To investigate whether method of loci training was associated with changes in the neural pattern similarity between the training study participants and memory athletes (or matched controls), we based our analysis on the beta-maps obtained from conventional across-trial modelling (thus, capturing average BOLD activity changes during word list encoding and temporal order recognition). For each participant, neural activity was extracted from the pre-defined ROIs and the data were reshaped into a pattern vector for each ROI. The data were then *z*-scored across voxels, and the pattern vectors of each ROI and each participant were correlated to the pattern vectors of all other participants of the same group, using pairwise Pearson’s correlation (*r*). The resulting correlations were Fisher’s *z*-transformed. Trainee-athlete neural pattern similarity for each subject was then computed as the average pattern similarity of each participant in training group with those of each memory athlete, and, separately, with those of each individual of the matched control group (that is, the average correlation of voxel patterns from participant 1 with those of participants 1∼*N* of athlete group or matched control group, separately). This procedure was repeated for both the pre- and post-training sessions, and separately for the word list encoding and temporal order recognition tasks, as well as separately for the different groups in the training study. This allowed us to compare the average neural pattern similarity between *time points* (pre-vs. post-training) and *groups* (neural pattern similarity between trainees × athletes vs. trainees × matched controls; Fig. 3a). The following equations were used to quantify the trainee-athlete RSA pattern similarity of individuals in the different experimental conditions. In brief, we calculated the pattern similarity of each participant in the training study (here denoted as sub_1_; to provide an example, we chose a participant of the memory training group, LOC) to the group average of that athlete group during the pre-(Equation 1) and post-training (Equation 2) sessions, for example, during word list encoding. Equations 3 and 4 follow the same rationale to assess the pattern similarity of the same sub_1_ to the group average of that matched control (CON). In these equations, *n* denotes the total number of participants in the respective groups, and j represents the participant index for participants in athletes or matched control groups (For details on statistical analysis, see the section “Statistical analyses and thresholding”).

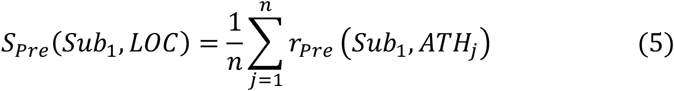

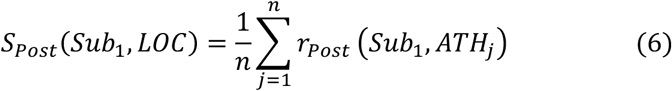

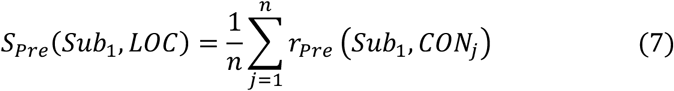

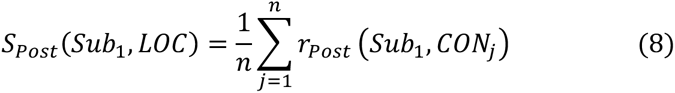

The trainee-athlete RSA hence examined the extent to which each trainee’s neural activity pattern during word list encoding (or temporal order recognition) aligned with the group-averaged neural representation of memory athletes (or matched controls). This approach allowed us to assess the degree of neural alignment between trainees and memory athletes.

#### Searchlight-based RSA

To go beyond the pre-defined ROIs and to obtain whole-brain pattern similarity results, we repeated the above-described RSA analyses but employed a searchlight with a radius of 6 mm (93 voxels) that was moved across the brain volume of each participant (Wagner, 2016; Wagner et al., 2016, 2020). Searchlights were considered only if they contained a minimum of 30 grey matter voxels. The average pattern similarity was calculated for each searchlight, and the resulting value was assigned to the centre voxel of the respective searchlight. This process was then repeated until each voxel had served as the searchlight centre once. The searchlight procedure was repeated for both pre- and post-training sessions, resulting in 3-dimensional whole-brain pattern similarity maps (2 for per task, 2 per session) per participant.

In the between-subject and trial-specific RSA analyses, we first examined the athlete study. Searchlight maps were subjected to independent-samples *t*-tests in a second-level group analysis to identify expertise-related differences in neural representations, comparing memory athletes to matched controls. For the training study, paired-sample *t*-tests were conducted to assess training-related changes in neural representations between pre- and post-training sessions. Additionally, a one-way ANOVA was performed in the training study to directly compare neural representations across the three experimental groups (memory training group, active control group, and passive control group. Finally, in the trainee-athlete RSA analysis, searchlight maps were analysed using a flexible 2 × 2 ANOVA design to investigate training-related changes in neural representations across the training study and the memory athlete study. The resulting *t*-statistic maps were then converted into *Z*-statistic maps using SPM’s implementation of Gaussian random field theory (GRFT). This transformation takes into account both the degrees of freedom and the spatial smoothness of the data, allowing for valid voxel- and cluster-level inference (Friston et al., 1996).

### Prediction analysis

We then tested the behavioural relevance of changes in neural representations between all individuals of the training study and memory athletes with respect to long-term memory performance. A balanced 4-fold cross-validation procedure (Cohen, 2010; Supekar et al., 2013) was employed to predict the improvement in free recall performance resulting from method of loci training. Specifically, we modelled the change in free recall performance from baseline to two long-term re-test intervals: 24 hours and 4 months. These predictions were based on changes in neural pattern similarity between all individuals of the training study and athletes from pre- to post-training. For each model, we included one dependent variable, such as free recall performance improvement at the 24-hour re-test (performance at the 24-hour post-training session minus the 24-hour pre-training session), or the 4-month re-test (performance at 4 months minus the 20-minute pre-training session). The independent variables differed by task: i) word list encoding: the increased distinct neural representations between all individuals of the training study and memory athletes (i.e., neural pattern similarity pre-training minus post-training) were used to test the behavioural relevance of distinct neural representations with respect to long-term memory performance. ii) temporal order recognition: the increased shared neural representations between all individuals of the training study and memory athletes (i.e., neural pattern similarity post-training minus pre-training) were used to evaluate the behavioural relevance of shared neural representations among experienced individuals in relation to long-term memory performance.

The relationship between the independent and dependent variables was quantified as *r*_(predicted,observed)_, measuring the degree to which the independent variables predicted the dependent variable. This measure was derived using the balanced 4-fold cross-validation procedure, in which the data for all individuals in the training study were divided into four equal subsets while ensuring that the distributions of dependent and independent variables were balanced across folds. This means that to validate the robustness of the relationship between training-related changes in participants’ neural representations and individual memory performance, we tested the model’s generalizability to out-of-sample individuals. In each iteration, the linear regression model was trained on three folds, and the held-out fold was used for prediction. This process was repeated until each fold served as the prediction sample once. The final *r*_(predicted, observed)_ value represented the average correlation between predicted and observed data across all folds.

To assess statistical significance, we employed a nonparametric approach. For instance, under the null hypothesis, no significant relationship would exist between free recall performance after 4 months (dependent variable) and trainee-athlete neural representation alterations (independent variable) in the specified ROIs. We generated 1000 surrogate datasets with randomly shuffled participant labels that were denoted as D_i_. These surrogate datasets were created by permuting the labels assigned to the observed data points. The *r*_(predicted, observed)*i*_ was computed by using the true labels of D*i* and the predicted labels generated through the 4-fold cross-validation procedure. This yielded a null distribution of *r*_(predicted, observed)_ (Cohen, 2010). The *p-*value was determined by counting the number of times when *r*_(predicted, observed)*i*_ was greater than r_(predicted, observed),_ divided that count by the number of D*i* datasets ([*r*_(predicted, observed)*i*_ > *r*_(predicted, observed)_]/D*i,* D*i* is 1000 in our case)(Qin et al., 2014; Zhu et al., 2022).

### Statistical analyses and thresholding

#### Statistical analysis of behavioural data and ROI-based RSA results

Analyses of ROI-based RSA results (including between-subject, trial-specific and trainee-athlete RSA) were conducted using jamovi (version 2.3.13.0; https://www.jamovi.org/). For the between-subject and trial-specific RSA, data from the athlete study (comparing memory athletes to matched controls) were subjected to independent-samples *t*-tests to evaluate expertise-related differences in neural representations. In the training study, paired-sample *t*-tests were used to examine training-related changes in neural representation between pre- and post-training sessions. For the trainee-athlete RSA, the average neural pattern similarity was compared across time points (pre-vs. post-training) and groups (neural pattern similarity between trainees × athletes vs. trainees × matched controls). These comparisons were further analysed using a 2 × 2 repeated-measures ANOVA to assess training-related changes in neural representations between the training study and the athlete study. Post-hoc pairwise comparisons following significant ANOVA effects were adjusted for multiple comparisons using the Bonferroni correction. Effect sizes were reported as Cohen’s d for *t*-tests and eta squared (*η²*) for ANOVA models (Lakens, 2013). To account for multiple comparisons across the four selected regions of interest (ROIs), false discovery rate (FDR) correction using the Benjamini–Hochberg procedure was applied at a threshold of *q* < 0.05. This correction was conducted separately for the word list encoding task and the temporal order recognition task. For all analyses, the significance level (*α*) was set at 0.05 (two-tailed).

#### Statistical thresholding of whole-brain RSA results and anatomical labelling

Unless stated otherwise, significance for all whole-brain fMRI analyses was assessed using cluster-based inference with a cluster-defining threshold of *p* < 0.001 and a cluster probability of *p* < 0.05 family-wise error (FWE) corrected for multiple comparisons. The corrected cluster size (i.e., the spatial extent of a cluster that is required to be labelled as significant) was calculated using the SPM extension “CorrClusTh.m” and the Newton-Raphson search method (script provided by Thomas Nichols, University of Warwick, United Kingdom, and Marko Wilke, University of Tübingen, Germany; http://www2.warwick.ac.uk/fac/sci/statistics/staff/academic-research/nichols/scripts/spm/). Anatomical nomenclature for all tables was obtained from the Laboratory for Neuro Imaging (LONI) Brain Atlas (LBPA40, www.loni.usc.edu/atlases/).

## Data availability

Raw, anonymised fMRI data are available upon reasonable request to the corresponding authors. At present, participant informed consent does not allow for depositing the full data set. All analyses are based on openly available software (see section “Code availability” below). Source data to reproduce all graphs (behavioural performance and ROI-based results of all analyses), as well as (un-)thresholded statistical whole-brain fMRI maps and their corresponding results tables, are openly available at the Open Science Framework (https://osf.io/7hb53/).

## Code availability

All analyses are based on openly available software or custom code that is provided on the Open Science Framework (https://osf.io/7hb53/).

## Supplementary Materials

### Supplementary Results S1: Gradient analysis, lateral prefrontal cortex (LPFC) and hippocampus

#### Word list encoding, analysis of LPFC gradients

Based on prior evidence of functional specialisation within the left LPFC (e.g., Nee & D’Esposito, 2016; Schumacher et al., 2019), we divided the LPFC along the dorsal-to-ventral axis (*z*-axis) into three functionally distinct subregions: dorsal, middle, and ventral LPFC. This tripartite division is based on accumulating evidence of the subregions supporting different cognitive functions (Koechlin et al., 2000, 2003; Nee & D’Esposito, 2016; Schumacher et al., 2019). The dorsal LPFC has been implicated in higher-order executive control, including abstract rule representation and hierarchical cognitive control. The middle LPFC is commonly associated with working memory maintenance and manipulation, playing a central role in information integration during complex tasks. The ventral LPFC appears more closely linked to semantic processing, response inhibition, and controlled retrieval from long-term memory.

##### Methodology

Identical to our main analysis, we used the Brainnetome Atlas (Fan et al., 2016) in MNI152 standard space to define the left PFC. The MNI-space coordinates of the resulting voxels were divided into three subregions along the dorsal-to-ventral axis based on the 33rd and 66th percentiles, spanning from *z* = -10 to *z* = 32. The LPFC mask yielded the dorsal (*z* = 14 to *z* = -32, 685 voxels), middle (*z* = 5 to *z* = 13, 615 voxels), and ventral LPFC (*z* = -10 to *z* = 4, 861 voxels; Fig. S9a). We then extracted the voxel activity patterns of each ROI during word list encoding and conducted representational similarity analysis (RSA) using three approaches (identical to our main analyses): (1) between-subject RSA, (2) trial-specific RSA, and (3) trainee–athlete RSA.

##### Results, between-subject RSA

In line with our initial results, we found significantly decreased neural pattern similarity in the dorsal LPFC in memory athletes compared to controls during the word list encoding task (separate independent-sample *t*-tests, memory athletes, *n* = 17, matched controls, *n* = 16, *t*(31) = -4.61, *p* < 0.001, Cohen’s *d* = -1.605; Fig. S10a). Individuals of the memory training group showed significantly decreased neural pattern similarity in the ventral LPFC after (compared to before) training (separate paired-sample *t*-tests, *n* = 17, *t*(16) = -2.691, *p* = 0.016, Cohen’s *d* = -0.653; Fig. S10b). This effect was specific to the memory training group, as neither the active nor the passive controls showed significant decreases in neural pattern similarity in any of the LPFC subregions (all *p* > 0.05, Fig. R8c, d, only active control group showed increased neural pattern similarity in the dorsal LPFC, Fig. S10c). There were no significant effects for the middle LPFC.

##### Results, trial-specific RSA

Similar to the above, the trial-specific RSA indicated significantly decreased neural pattern similarity in the dorsal LPFC in memory athletes compared to controls during the word list encoding task (separate independent-sample *t*-tests, memory athletes, *n* = 17, matched controls, *n* = 16, *t*(31) = -2.89, *p* = 0.007, Cohen’s *d* = -1.005; Fig. S11a). The memory training group showed a trend toward decreases in trial-specific neural pattern similarity in the middle (*p* = 0.034) and ventral LPFC (*p* = 0.022) post-training (note that these results did not survive multiple comparison correction but exhibited a trend toward significance, Fig. S11b). Again, these effects were specific to the memory training group, as neither the active control nor the passive control groups demonstrated comparable training-related decreases across any LPFC subregions (all *p* > 0.05, Fig. S11c, d).

##### Results, trainee-athlete RSA

After training, the memory training group exhibited no significant change in their neural pattern similarity to athletes in the LPFC in any of the subregions, and no significant effects were found for the control groups (all *p* > 0.05, see results summary in Table S10). This may suggest a synergistic effect of different subfields that is not captured when partitioning the ROI along its dorsal-to-ventral axis.

##### Summary & conclusions

Memory athletes showed decreased neural pattern similarity between individuals compared with matched controls, predominantly in the dorsal LPFC. The memory training group exhibited reduced neural pattern similarity between individuals, mainly in the ventral LPFC, after training. These findings suggest that distinct LPFC subregions are differentially engaged in method of loci processing in athletes and trainees. This pattern may reflect differences in cognitive strategies, with experts potentially relying more on executive functions (dorsal), while trainees may still recruit semantic or integrative processes during learning (ventral).

#### Temporal order recognition, analysis of hippocampal gradients

Based on prior evidence of hippocampal functional specialisation along its long axis (Brunec et al., 2018; Collin et al., 2015; Poppenk et al., 2013), we divided the hippocampus into anterior, middle, and posterior segments as they were shown to be functionally distinct (Collin et al., 2015): The anterior hippocampus is involved in affective processing and supports the abstraction of gist-like, schematic memory representations (Poppenk et al., 2013). The middle hippocampus is thought to facilitate memory integration, including the binding of episodic details (Collin et al., 2015). The posterior hippocampus appears primarily engaged in processing spatial and contextual information and plays a role in the retrieval of episodic memories (Maguire et al., 2000; Poppenk et al., 2013).

##### Methodology

Identical to our main analysis, we used the Brainnetome Atlas (Fan et al., 2016) in MNI152 standard space to define the bilateral hippocampus. We retained the bilateral hippocampus as this made the novel results comparable to our main analysis, to our previous work (Wagner et al., 2021), and to previous work that showed no lateralisation of hippocampal event representations (Collin et al., 2015). The MNI-space coordinates of the resulting voxels ranged from *y* = -42 to *y* = -2 (posterior-to-anterior). We then divided the hippocampal mask into three equal-length segments that yielded the posterior (*y* = -42 to *y* = - 32, 1306 voxels), middle (*y* = -31 to *y* = -21, 472 voxels), and anterior hippocampus (*y* = -20 to *y* = -2, 483 voxels; Fig. S9b). We then extracted the voxel activity patterns of each ROI during temporal order recognition and conducted representational similarity analysis (RSA) using three approaches (identical to our main analyses): (1) between-subject RSA, (2) trial-specific RSA, and (3) trainee–athlete RSA.

##### Results, between-subject RSA

Findings revealed significantly increased neural pattern similarity in the anterior and middle hippocampus in memory athletes compared to controls during temporal order recognition (separate independent-sample *t*-tests, memory athletes, *n* = 17, matched controls, *n* = 16, *t*(31)_anterior_ = 7.105, *p* < 0.001, Cohen’s *d* = 2.475; *t*(31)_middle_ = 3.546, *p* = 0.001, Cohen’s *d* = 1.235, Fig. S12a). Similarly, individuals of the memory training group showed significantly increased neural pattern similarity in the anterior and posterior subregions after (compared to before) training (separate paired-sample *t*-tests, *n* = 17, *t*(16)_anterior_ = 3.300, *p* = 0.005, Cohen’s *d* = 0.799; *t*(16)_posterior_ = 3.100, *p* = 0.007, Cohen’s *d* = 0.751, Fig. S12b). This effect was specific to the memory training group, as neither the active nor the passive controls showed significant increases in neural pattern similarity in any of the hippocampal subregions (all *p* > 0.05, Fig. R12c, d).

##### Results, trial-specific RSA

The trial-specific RSA indicated no significant differences between memory athletes and controls (all *p* > 0.05, Fig. S13a). The memory training group showed significant increases in trial-specific neural pattern similarity in the anterior, middle, and posterior hippocampus post-training (memory training group, *n* = 17, *t*(16)_anterior_ = 4.140, *p* < 0.001, Cohen’s d = 1.004; *t*(16)_middle_ = 3.150, *p* = 0.006, Cohen’s *d* = 0.764; *t*(16)_posterior_ = 3.440, *p* = 0.003, Cohen’s *d* = 0.835; Fig.S13b). Again, this effect was specific to the memory training group, as neither the active control nor the passive control groups demonstrated comparable training-related increases across any hippocampal subregions (Fig. S13c, d).

##### Results, trainee-athlete RSA

The trainee-athlete RSA showed a trend toward increased neural pattern similarity in the posterior hippocampus when trainees had completed the method of loci training (separate repeated measures ANOVA, *n* = 17, main effect of time point: F (1,16)_posterior_ = 3.973, *p* = 0.064, *η_p_*^2^ = 0.199), and significantly lower neural pattern similarity between trainees × athletes compared to pattern similarity between trainees × matched controls in all hippocampus subregions (main effect of group: F (1, 16)_anterior_= 6.412, *p* = 0.022, *η_p_*^2^ = 0.286; F (1, 16)_middle_= 6.282, *p* = 0.023, *η_p_*^2^ = 0.282; F (1,16)_posterior_ = 6.601, *p* = 0.021, *η_p_*^2^ = 0.292). We also found a trend toward a significant time point × group interaction (F (1, 16) = 3.959, *p* = 0.064, *η_p_*^2^ = 0.198) in the anterior hippocampus, suggesting that neural pattern similarity between trainees × athletes (*p* = 0.065), but not between trainees × matched controls (*p* = 0.672), was higher in the post-compared to pre-training session(Fig.S14). No such effects were found for other group comparisons or hippocampal subregions (see results summary in Table S11).

##### Summary & conclusions

These findings underscore a functional dissociation along the hippocampal long axis. First, between-subject RSA showed that memory athletes exhibited higher neural pattern similarity between individuals in the anterior and middle hippocampus compared to matched controls, whereas the memory training group, after training, showed increased neural pattern similarity between individuals in both the anterior and posterior hippocampus. Second, trial-specific RSA revealed no significant differences in neural pattern similarity between trials across any hippocampal subregion when comparing memory athletes to matched controls. However, the memory training group demonstrated increased neural pattern similarity between trials in all hippocampal subregions after training, relative to their pre-training. Finally, trainee-athlete RSA indicated a trend toward increased neural pattern similarity between the memory training group and memory athletes in the anterior hippocampus after training.

Together, these results may suggest that, following training, individuals in the memory training group engaged both detailed perceptual and abstract mnemonic processes, as reflected by increased pattern similarity in the posterior and anterior hippocampus, respectively. In contrast, memory athletes appeared to have relied more on integrative and schematic processing, particularly in the middle and anterior hippocampus (Brunec et al., 2018; Collin et al., 2015; Poppenk et al., 2013).

### Supplementary Results S2: Region-of-interest (ROI) analysis of the medial temporal lobe (MTL) during word list encoding

We focused on the hippocampus and parahippocampal gyrus during word list encoding due to their involvement in the method of loci (Fellner et al., 2016; Liu et al., 2022; Maguire et al., 2003; Nyberg et al., 2003), visuospatial processing, and navigation (Epstein, 2008; Hartley et al., 2014; Hassabis & Maguire, 2009; Pearson, 2019). Bilateral ROI masks were defined using the Brainnetome atlas (Fan et al., 2016; https://atlas.brainnetome.org). For each ROI, we examined the neural representations during word list encoding using the following representational similarity analysis (RSA) approaches: (1) between-subject RSA, (2) trial-specific RSA, and (3) trainee-athlete RSA.

#### Results

Interestingly, results from the between-subject RSA showed significantly increased neural pattern similarity in the hippocampus between memory athletes compared to matched controls during word list encoding (separate independent-sample *t-*tests, memory athletes, *n* = 17, matched controls, *n* = 16, *t*(31) = 3.028, *p* = 0.005, Cohen’s *d* = 1.055, Fig. S15a). There was no significant difference in neural pattern similarity in the parahippocampal gyrus between these groups (*p* = 0.571, Fig. S15a). Turning toward the data of the training study, neural pattern similarity was significantly increased in the parahippocampal gyrus (separate paired-sample *t*-tests, *n* = 17, *t* (16) = 2.965, *p* = 0.009, Cohen’s *d* = 0.719, Fig. S15b) but not in the hippocampus (but note the clear trend, *p* = 0.058, Fig. S15b) between the individuals of the memory training group after (compared to before) training. There were no significant effects in the active or passive control groups (all *p* > 0.05, Fig. S15c, d), and neither the trial-specific RSA nor the trainee-athlete RSA revealed significant effects of method of loci expertise on MTL pattern similarity (all *p* > 0.05; Fig. S16 and Fig.S17).

#### Summary & conclusion

Contrary to what we had expected, we found that method of loci training was related to increased (rather than decreased) MTL pattern similarity in experienced individuals. These results are difficult to align with those of Liu and colleagues (2022), but it is possible that an analysis of hippocampal subfields might reveal more nuanced effects. One prior study showed increased MTL pattern similarity during associative memory encoding (Wagner et al., 2016; but note that their approach corresponded to the trials-specific RSA; no analysis was conducted between-subjects). Increased neural pattern similarity could thus reflect the build-up of durable, longer-term memory associations, and we know that the experienced individuals in our sample formed a substantial proportion of durable memories (Wagner et al., 2021).

### Supplementary Results S3: Per-group for correlations between neural pattern similarity and free recall performance after 4 months

#### ROI-based correlations per group

We performed stratified within-group analyses across all primary ROIs. These analyses revealed no significant correlations within any of the groups (Table S12). We speculate that the lack of significance likely reflects the limited sample sizes in each group (memory training group *n* = 14, active control group *n* = 14, passive control group *n* = 16), reducing power. Moreover, the memory training group reached close-to-ceiling performance post-training (yielding low interindividual variability), potentially limiting our ability to detect significant correlations within this group.

#### Whole-brain, voxel-wise regression analysis

To go beyond the ROI-based analysis, and because the ROIs were quite large (making it possible that averaging signals across entire ROIs may have obscure more localized neural effects), we conducted a voxel-wise, whole-brain multiple regression analysis to further investigate the relationship between training-related neural changes and individual differences in memory performance.

#### Methodology

Using the trainee-athlete RSA, we first computed the contrast between post-training and pre-training sessions to capture overall training-related changes in neural similarity [Post(trainee × athlete) > Pre(trainee × athlete)]. Subsequently, within each group (memory training, active control, and passive control), we performed a voxel-wise linear regression analysis using individual memory improvement as a covariate of interest (calculated as the difference in free recall performance between the 4-month retest and the immediate free recall test during the pre-training session). A positive contrast weight (+1) on the covariate identified brain regions where increased neural similarity to memory athletes was associated with better memory performance, whereas a negative weight (-1) indicated regions where higher memory performance was related to reduced neural similarity.

#### Results

In the memory training group, we identified a significant cluster in the left lateral frontal cortex (peak MNI coordinates: *x* = −34, *y* = 26, *z* = 36; cluster size = 30 voxels; Fig. S18a), where lower neural pattern similarity between the memory training group (post-training) and memory athletes was associated with higher memory improvement (Fig. S18b, *p* < 0.05, FWE-corrected at the cluster level, with a cluster-defining threshold of *p* < 0.001 and a minimum cluster size of 27 voxels). This effect appeared to be specific to the memory training group, as no significant associations were identified in the active or passive control groups (Fig. S18c, d).

We also repeated the same whole-brain, voxel-wise linear regression analysis during temporal order recognition, but there were no significant associations between neural pattern similarity and memory performance (initial, ROI-based results were significant when we collapsed across all subjects of the memory training group). This is not surprising, as recognition performance was generally less affected by method of loci training (performance was numerically higher in the memory training group after training, but was not significantly different from the control groups, and did not reach the high levels as seen in memory athletes who had decades of experience).

### Supplementary Results S4: Encoding-Retrieval analysis of neural pattern similarity

We conducted two additional analyses aimed at investigating whether retrieval-related activity during the temporal order recognition task reflected the “reactivation” of encoding-related patterns (DuBrow & Davachi, 2014; Nyberg et al., 2000), and whether such similarity was modulated by memory training or expertise.

#### Trial-specific reactivation analysis

We assessed whether neural patterns associated with word triplets during retrieval resembled the neural patterns during encoding. For each participant, we extracted the trial-wise beta estimates from the word list encoding task (72 trials) and the temporal order recognition task (24 trials) within the main ROIs (left lateral prefrontal cortex, LPFC, left superior frontal gyrus, SFG, bilateral hippocampus, HIP, and bilateral precuneus, Prec). Because each retrieval trial involved a triplet of previously encoded words, we averaged the beta values of the three corresponding encoding trials to construct 24 encoding regressors, thereby matching the 24 retrieval trials. We then computed Pearson correlations between encoding and retrieval activity patterns across these trials. Correlation values were categorised into paired trials (matching triplets between encoding and retrieval) and unpaired trials (non-matching triplets), allowing us to test for trial-specific reactivation (matching triplet vs. non-matching triplet).

#### Modulation by memory training and expertise

To examine whether encoding-retrieval similarity was influenced by memory training or expertise, we compared the similarity values of paired trials across pre- and post-training sessions in the training group, and between memory athletes and matched controls.

#### Results

In both the memory athlete and the memory training group (post-training), we observed no significant difference in neural pattern similarity between paired and unpaired trials (all *p > 0.05*, Fig. S19). These results could indicate that reactivation likely occurred both during matching and non-matching triplets (because referring to the mental route is necessary to solve the task and to decide on the match). However, there were also no significant differences in encoding-retrieval similarity of paired trials across time points (pre-vs. post-training) in the memory training group, or between memory athletes and matched controls.

#### Conclusions

It is important to highlight the difference in task structure between the encoding and retrieval tasks. While encoding involved single-item imagery of word-loci pairs, retrieval was based on temporal sequence judgments of triplets. Even though we averaged across the corresponding trials during the word list encoding task, a clean analysis would require a different task design (e.g., showing word triplets also during encoding). Prior work has shown that encoding-reactivation effects are more robust when the same trial types are used (e.g., Xiao et al., 2017; Xue et al., 2010). Thus, while we did not observe significant encoding-retrieval reactivation effects in the current paradigm, this does not necessarily imply an absence of effects but rather highlights the importance of *a priori* choices in terms of task design that should be tailored to specific questions.

### Supplementary Results S5: Performance in the temporal order recognition task

Please note that we had already presented this data previously (Wagner et al., 2021), but we added it here once more for full transparency.

#### Performance during temporal order recognition (*d*-prime)

Recognition performance was quantified using *d*-prime scores. To accommodate hit rates of 1 and false alarm rates of 0 in memory athletes and the memory training group (post-training), we adjust the individual hit and false alarm rates (*z*-scored) of all participants by adding 0.5 to the raw counts of individual hit and false alarm rates (Stanislaw & Todorov, 1999). *D*-prime was calculated as the difference between these adjusted hit and false alarm rates [z(hits) − z(false alarms)], collapsing across the different confidence levels (“sure,” “maybe”), as memory athletes and participants of the memory training group (post-training) had very few “maybe” responses [number of “maybe” triplets, means ± SEM; athlete study, memory athletes: 0.47 ± 0.1, matched controls: 4 ± 0.43; training study (pre-/post-training), memory training group: 6.1 ± 0.61/1.53 ± 0.54, active controls: 4.47 ± 0.85/2.36 ± 0.61, passive controls: 3.35 ± 0.74/2.65 ± 0.85]. Also, there were very few missed responses, and we collapsed those together with incorrect triplets [number of missed triplets, means ± SEM; athlete study, memory athletes: 0.18 ± 0.1, matched controls: 0.29 ± 0.14; training study (pre−/post-training), memory training group: 0.59 ± 0.19/0.65 ± 0.17, active controls: 0.47 ± 0.17/0.29 ± 0.14, passive controls: 0.35 ± 0.12/0.35 ± 0.15].

#### Results

In brief, we found significantly higher recognition performance (*d*-prime) in the memory athletes compared to the matched controls (Wilcoxon signed-rank test; one matched pair excluded from analysis; memory athletes, *n* = 16, median = 3.54; matched controls, *n* = 16, median = 2.38; V = 131.5, *p* = 0.001; Table S13). However, despite numerically increased *d*-prime scores in the memory training group after training, there was no significant difference in recognition performance between the three groups (change in *d*-prime; one-way ANOVA; memory training group, *n* = 17; active controls, *n* = 16; passive controls, *n* = 17; main effect of group, *p* = 0.133, pairwise comparisons: memory training > active controls: *p* = 0.135, effect size *d* = 0.592; memory training > passive controls: *p* = 0.294, *d* = 0.569; active > passive controls: *p* = 0.898, *d* = −0.161; see results Table S13 for a general increase in *d*-prime from pre- and post-training). Thus, expertise in the method of loci positively affected recognition performance in memory athletes. While the effect of memory training on recognition performance in initially mnemonics-naïve participants appeared to be positive as well, the results were not significant. This may indicate an advantage of memory athletes, given their extensive experience that goes far beyond a mere 6-week training. Please see our previous work for an in-depth discussion of this finding (Wagner et al., 2021).

### Supplementary Results S6: Characterising the information represented in the MTL and precuneus during word list encoding and temporal order recognition

#### We performed two additional analyses to determine the specific type of information represented in the MTL and precuneus during word list encoding and temporal order recognition

First, if increased neural pattern similarity was indeed related to shared cognitive processes between experienced individuals, one would expect evidence for sequence memory (in other words, experienced individuals should have mentally navigated from one word-location association to the next). We performed a sequence-geometry analysis that quantified neural pattern similarity for word-location associations that were closer along the mental route and compared to those that were further apart. While this does not tackle *specific* stimulus content, it helps to understand whether experienced individuals may have mentally reinstated the loci routes.

Second, if experienced individuals formed and retrieved unique word-location associations during the tasks, one would expect specific neural representations (for the single words or triplets) between experienced individuals (in other words, experienced individuals used their own loci routes, which should be coupled to decreased neural pattern similarity). Complementary to our main analysis approach, we performed a trial-wise Intersubject Consistency (ISC) analysis that quantified neural pattern similarity for each word (or triplet) between individuals.

### Sequence-geometry analysis

#### Methodology

To quantify whether experienced individuals used sequential memory during the tasks (which would be an indication of method of loci processing), we focused on the individuals in the training study. This is because the memory training group was instructed to use loci routes that comprised 50 loci each (whereas memory athletes could use their own loci routes that likely varied in length). To provide a complete overview of the results, we analysed both the word list encoding as well as the temporal order recognition tasks, along with the relevant regions-of-interest (ROIs). During word list encoding, only the first 50 word-location associations were analysed (out of 72) because we were able to reliably map these onto one entire loci route, avoiding transitions between routes. During temporal order recognition, we restricted the analysis to the first 16 triplets (out of 24, thus 16*3=48 word-location associations to make sure we captured almost one entire loci route).

For both tasks, multivoxel patterns were extracted from the trial-wise beta estimates for each ROI (word list encoding: hippocampus, parahippocampal gyrus; temporal order recognition: hippocampus and its anterior/middle/posterior subdivisions, precuneus). For each participant, neural pattern similarity was computed separately for word list encoding (50 × 50 trials) and temporal order recognition (16 × 16 trials) by correlating across trials (Fig. 20a). Predicted similarity was modelled as a monotonic decrease with route distance (Fig. 20b), which belongs to a class of models that captures similarity decay in graph- and sequence-based representational spaces (Constantinescu et al., 2016; Garvert et al., 2017). Our primary focus was on the reciprocal distance function (Fig. 20c), but we repeated the analysis with Gaussian similarity kernels (Kriegeskorte et al., 2008; Shepard, 1987) to perform a robustness check, confirming that the effect of decreasing neural pattern similarity with increasing word-location distance did not depend on the specific choice of analysis parameter. Model fits were estimated by correlating the neural and theoretical matrices (Pearson’s *r*, Fisher *z*-transformed), yielding one model-alignment score per participant and ROI. Group-level statistics were performed with one-sample *t*-tests and one-way ANOVAs with the between-subject factor Group (memory training group, active controls, passive controls), followed by Tukey’s HSD post-hoc tests.

#### Results

During word list encoding, neural pattern similarity in the hippocampus and parahippocampal gyrus showed a significantly positive alignment with the sequence-geometry model in all participants (post-training session; one-sample *t*-tests, all *p* < 0.05, Fig. 20d shows an example model fit in the hippocampus). There were no significant group differences (all *p* > 0.05, Fig.20f), suggesting that sequence information was similarly represented across individuals, unaffected by method of loci training. As expected, no group differences were detected before training (all *p* > 0.05; not shown).

During temporal order recognition, neural pattern similarity in the MTL and precuneus showed a significantly positive alignment with the sequence-geometry model in all participants (post-training session; one-sample *t*-tests, all *p* < 0.05, Fig. 20e shows an example model fit in the hippocampus), but there was a main effect of Group in the hippocampus (*F*(2, 47) = 2.9, *p* = 0.062, *η*² = 0.112, trend-level) and its anterior subdivision (*F*(2, 47) = 3.82, *p* = 0.029, *η*² = 0.140; Fig.20f). Specifically, neural patterns during temporal order recognition were significantly more aligned with sequential trial order in the memory training group compared to passive controls in the hippocampus and its anterior subdivision after training (*p* = 0.049 and *p* = 0.025, respectively; Fig. 20g; but not compared to active controls, *p* > 0.05). We would like to note that it is not particularly surprising to find alignment with the sequence-geometry model in all participants. This is because of the temporal autocorrelation of the fMRI signal (trials closer in time will naturally be more similar to each other because of the BOLD signal correlation). However, we focus on group differences, showing stronger sequence-geometry alignment in the memory training group after training compared to controls. As above, no group differences were present before training (all *p* > 0.05; not shown).

As a robustness check, we also tested a Gaussian similarity kernel, which (like the reciprocal kernel) implements a monotonic decrease in neural pattern similarity with increasing word-location distance. The Gaussian includes a bandwidth parameter *σ* that controls the steepness of this decay: smaller *σ* values produce a sharp drop-off so that only neighbouring loci appear similar, whereas larger *σ* values lead to a slower drop-off such that distant loci still retain their similarity. In our data, the Gaussian kernel reproduced the abovementioned group differences (stronger model fit for the memory training group > passive controls, post-training) for smaller *σ* values in the anterior hippocampus (1.5, 2.5, and 3.5; all *p* < 0.05), confirming consistency with the reciprocal model, while the effect diminished at larger *σ* (4.5, *p* = 0.076, as is to be expected, indicating parameter-dependence).

### Trial-wise Intersubject Consistency (ISC) analysis

#### Method

To quantify whether experienced individuals processed unique word-location associations during word list encoding and temporal order recognition, we focused on the individuals in the athlete study. This is because memory athletes use their own loci routes that varied between individuals (i.e., the memory training group trained 1+3 loci routes: 1 route around the Max Planck Institute that was common to all, plus 3 self-constructed routes; because we did not have information on which loci routes participants utilised post-training, we focused on the memory athletes as we could be sure that the loci routes would differ between them).

As above, multivoxel patterns were extracted from the trial-wise beta estimates for each ROI and task (word list encoding: hippocampus, parahippocampal gyrus; temporal order recognition: hippocampus and its anterior/middle/posterior subdivisions, precuneus). For each trial, a voxel-by-subject matrix of beta estimates was constructed. Within each participant, voxel patterns were *z*-scored, and pairwise Pearson’s correlations were computed across participants to obtain a correlation matrix for each trial. Off-diagonal correlations were Fisher’s *z*-transformed and averaged across all other participants to yield a trial-level ISC value per subject. These values were subsequently averaged across all 72 word-location associations during word list encoding (we analysed all 72 because our focus was on the similarity across participants, irrespective of route transitions), and across 24 triplets during temporal order recognition. This was done for each ROI, resulting in one trial-wise ISC estimate per subject (Fig. 21a). Group differences in trial-wise ISC were then assessed by independent-samples t-tests between memory athletes and matched controls.

#### Results

During word list encoding, we found no significant group differences in trial-wise ISC in the hippocampus or parahippocampal gyrus after training (all *p* > 0.05, Fig.21b; no group differences were present before training, all *p* > 0.05; not shown). During temporal order recognition, memory athletes showed higher trial-wise ISC than matched controls in the precuneus (independent-samples *t*-test, *t(*31) = 4.42, *p* = 0.00011, *d* = 1.54, Fig.21c; again, no group differences were present before training, all *p* > 0.05; not shown). Put differently, neural patterns in the precuneus that were associated with different (triplets of) word-location associations were more similar between memory athletes than between matched controls during temporal order recognition. Neural patterns thus appeared shared between experienced individuals.

#### Summary & conclusion

First, we found that neural patterns in the (anterior) hippocampus were explained by sequence geometry during temporal order recognition. This was the case for the memory training group after training (compared to passive controls; but note that there was no significant difference compared to active controls – we speculate that these individuals may have come up with their own strategy to recall words in a sequence-like manner), suggesting that method of loci training may have promoted the mental reinstatement of loci routes in a hippocampal-entorhinal circuit that was shown to encode abstract relational and temporal information (Constantinescu et al., 2016; Eichenbaum, 2013; Garvert et al., 2017; Hsieh et al., 2014; Schapiro et al., 2013).

Second, we found increased trial-wise ISC (thus, increased trial-wise neural pattern similarity between individuals) in the precuneus of memory athletes after training (compared to matched controls). This is in line with theoretical accounts that link precuneus engagement to scene construction and context reinstatement (Diana et al., 2007; Ranganath & Ritchey, 2012), suggesting shared cognitive processes across different individuals.

Altogether, we found increased neural pattern similarity in the MTL and precuneus between experienced individuals, which may be related to the fact that they processed associative- and sequence-based memory content in a similar manner. We fully acknowledge that our additional analyses cannot disentangle the precise underlying mechanisms of method of loci processing (also keeping in mind the methodological limitations of fMRI-based research; Ramsey et al., 2010). Future work may build on our findings by employing methods with higher temporal resolution or direct neural recordings to further characterise the neural mechanisms of mental reinstatement during method of loci processing.

## Supplementary Figures

**Fig. S1.**
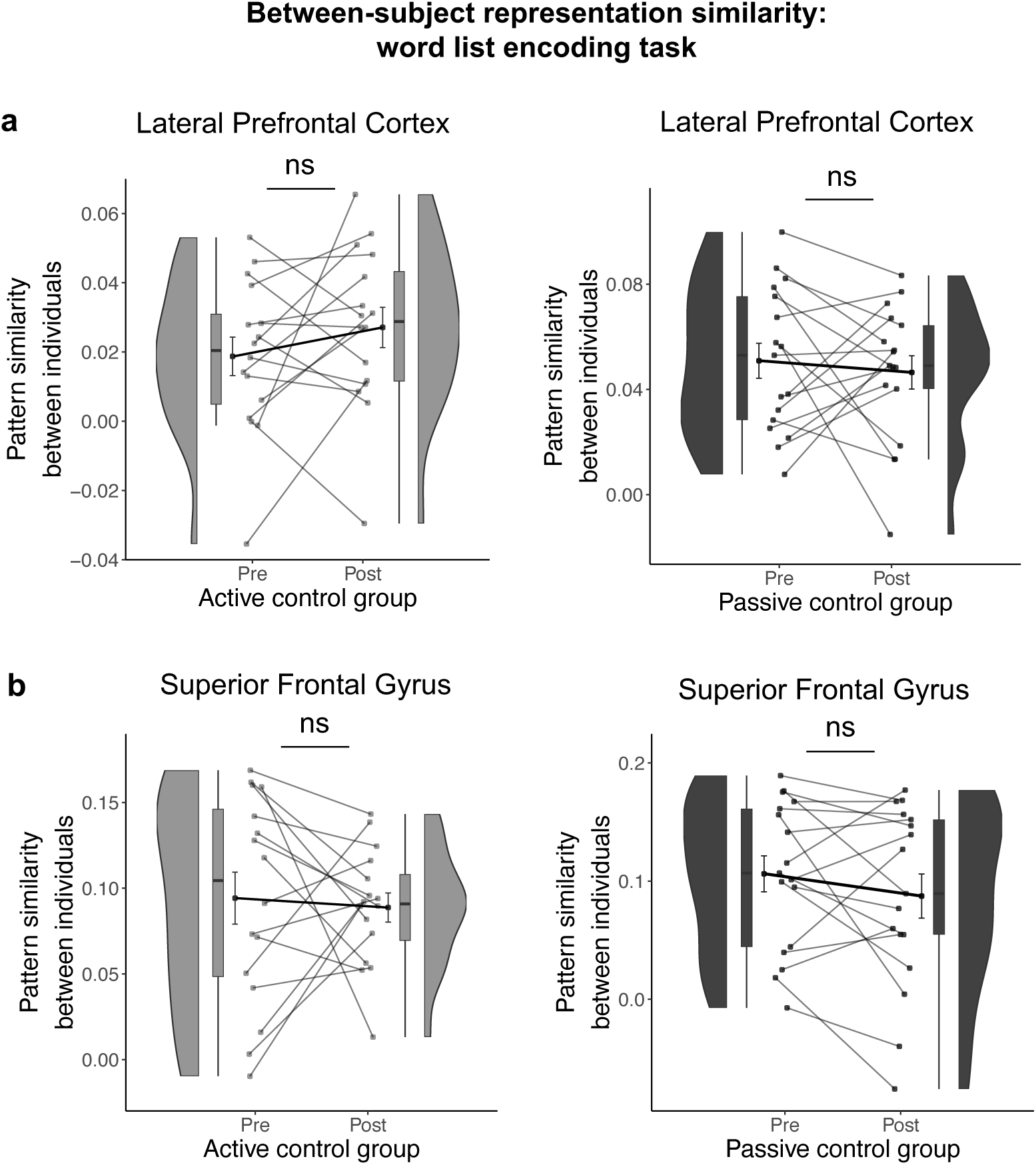
Results for between-subject ROI-based RSA during word list encoding, active and passive control groups. Regions-of-interest (ROIs) used in the between-subject RSA during word list encoding (i.e., active controls × active controls, passive controls × passive controls, analyzed separately for pre- and post-training sessions) included the (**a**) lateral prefrontal cortex and (**b**) superior frontal gyrus. Figure legend: Data points represent individual participants (active controls, *n* = 16; passive controls, *n* = 17); boxplots display the median and upper/lower quartiles, whiskers show 1.5 interquartile ranges above/below the quartiles; circles with error bars correspond to the mean ± standard error of the mean (S.E.M.); distributions illustrate the probability density function of individual data points. Abbreviations: ns, not significant. Pattern similarity reflects Fisher’s *z*-transformed Pearson’s correlations (*r*).

**Fig. S2.**
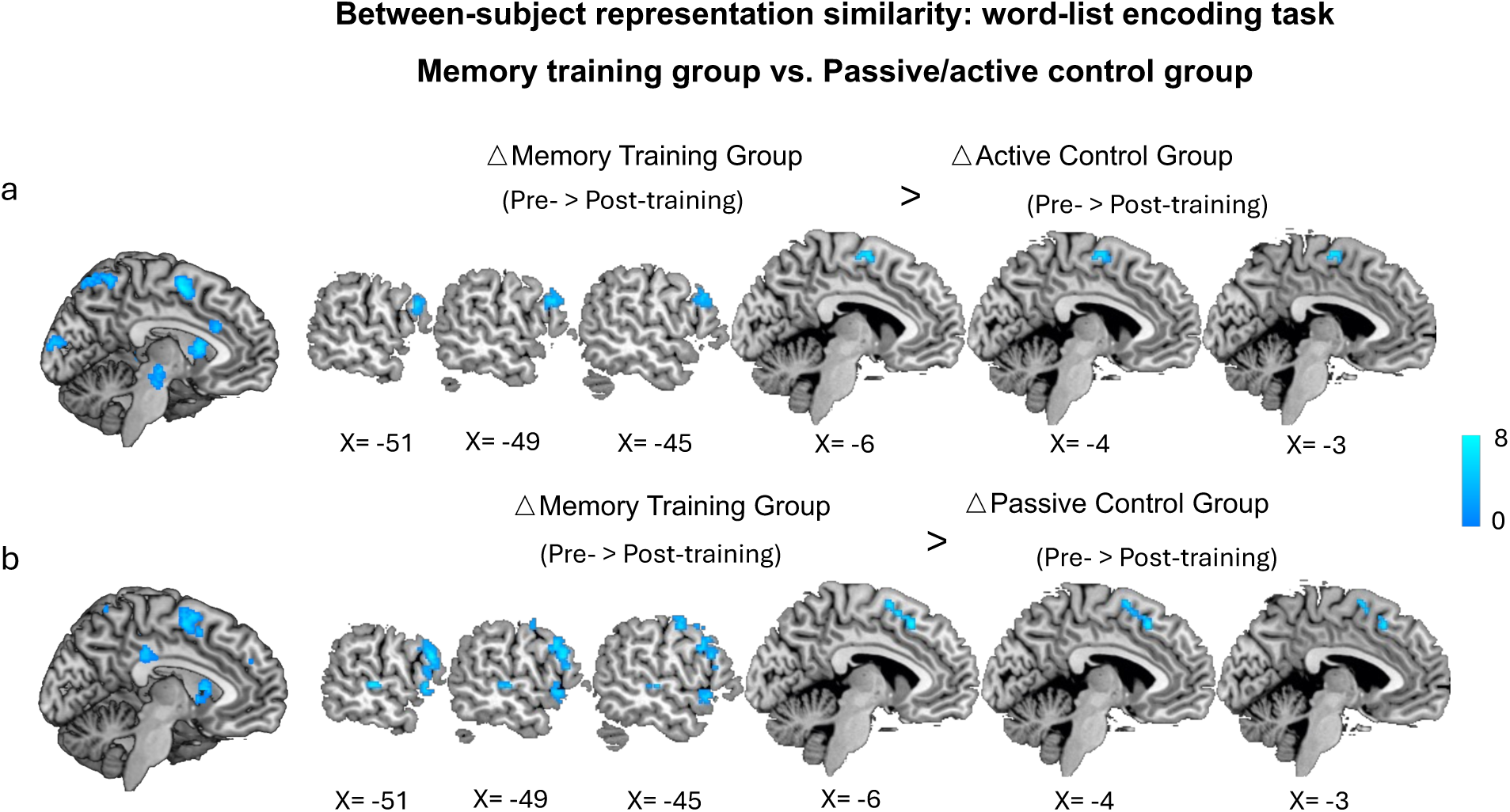
Method of loci leading to distinct neural representation among individuals is stronger than active and passive control groups. Searchlight RSA of the increased between-subject distinct neural representation (pre-minus post-training) in the memory training group after training during the word list encoding task, compared to (**a**) active or (**b**) passive control. Results are reported at *p* < 0.05, family-wise error (FWE)-corrected at the cluster level (cluster-defining threshold *p* < 0.001). Colour bar indicates *t*-values.

**Fig. S3.**
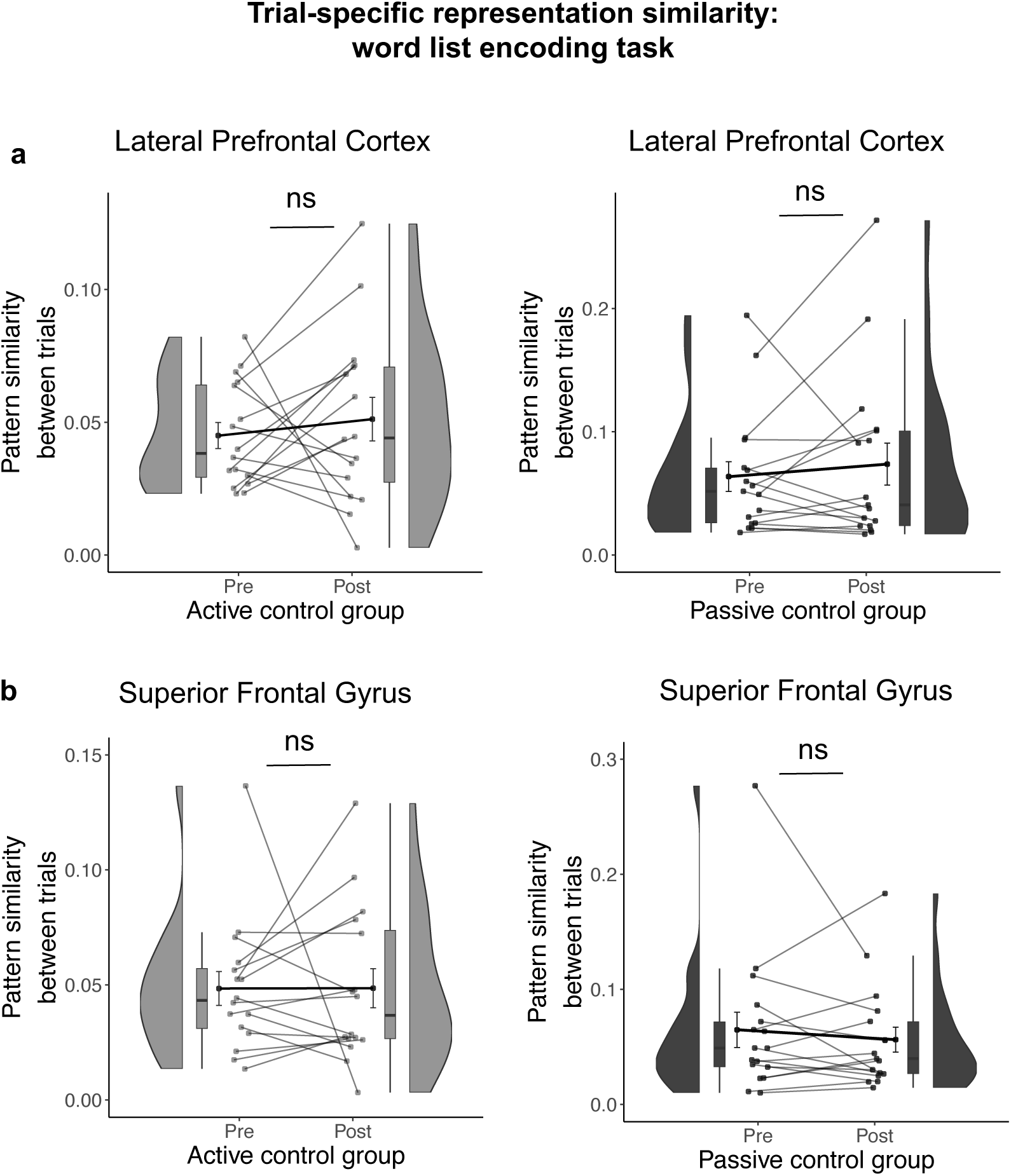
Results for trial-specific ROI-based RSA during word list encoding, active and passive control groups. Regions-of-interest (ROIs) used in the trial-specific RSA during word list encoding (i.e., active controls x active controls, passive controls × passive controls, analyzed separately for pre- and post-training sessions) included the (**a**) lateral prefrontal cortex and (**b**) superior frontal gyrus. Figure legend: Data points represent individual participants (active controls, *n* = 16; passive controls, *n* = 17); boxplots display the median and upper/lower quartiles, whiskers show 1.5 interquartile ranges above/below the quartiles; circles with error bars correspond to the mean ± standard error of the mean (S.E.M.); distributions illustrate the probability density function of individual data points. Abbreviations: ns, not significant. Pattern similarity reflects Fisher’s *z*-transformed Pearson’s correlations (*r*).

**Fig. S4.**
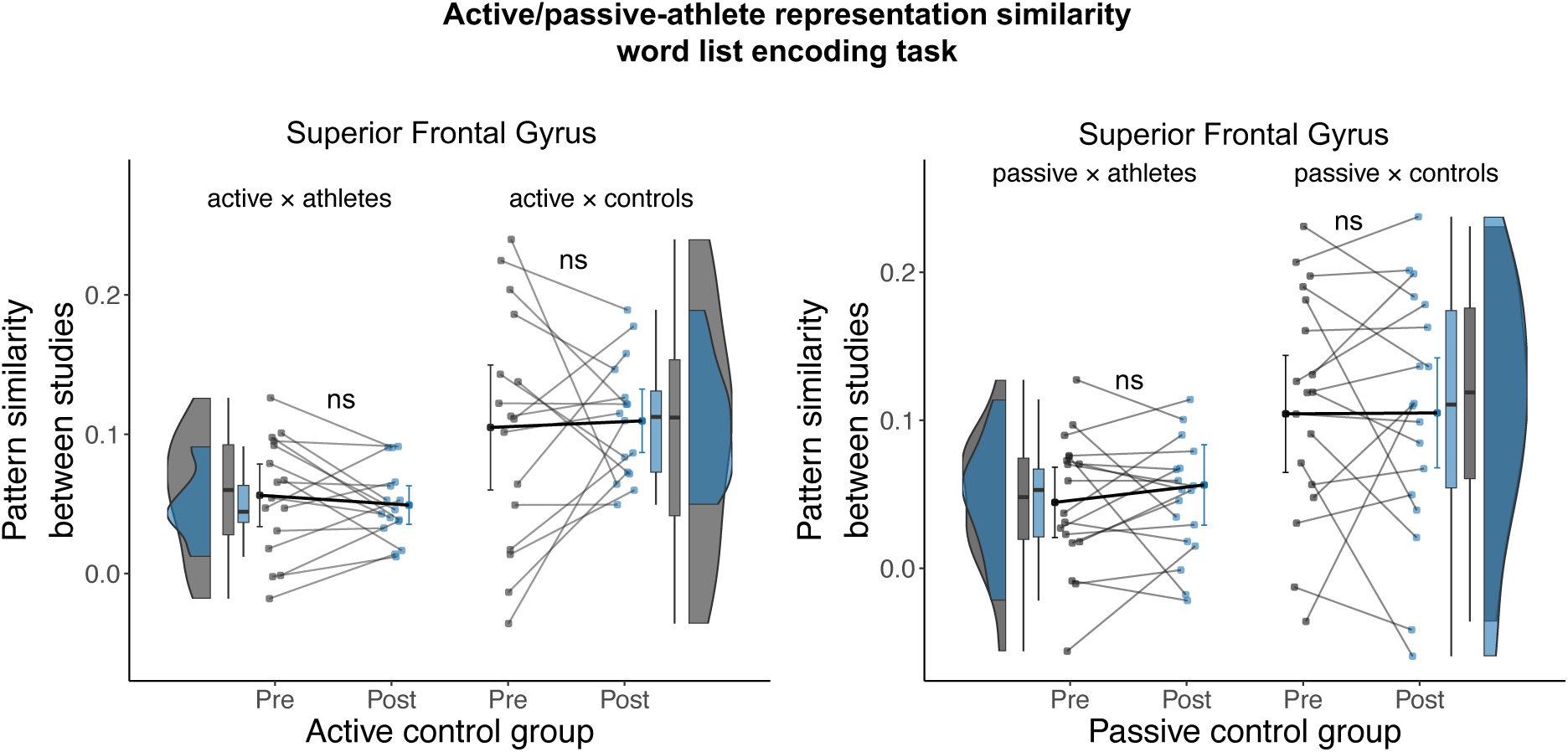
Results for trainee-athlete ROI-based RSA during word list encoding, active and passive control groups. Regions-of-interest (ROIs) are used in the trainee-athlete RSA during word list encoding, analyzed with repeat measures ANOVA of 2 × 2 factors: *time point* (pre-vs. post-training) and *group* (neural pattern similarity between active/passive controls × athletes vs. active/passive controls × matched controls). Figure legend: Data points represent individual participants (active controls, *n* = 16; passive controls, *n* = 17); boxplots display the median and upper/lower quartiles, whiskers show 1.5 interquartile ranges above/below the quartiles; circles with error bars correspond to the mean ± standard error of the mean (S.E.M.); distributions illustrate the probability density function of individual data points. Abbreviations: ns, not significant. Pattern similarity reflects Fisher’s *z*-transformed Pearson’s correlations (*r*).

**Fig. S5.**
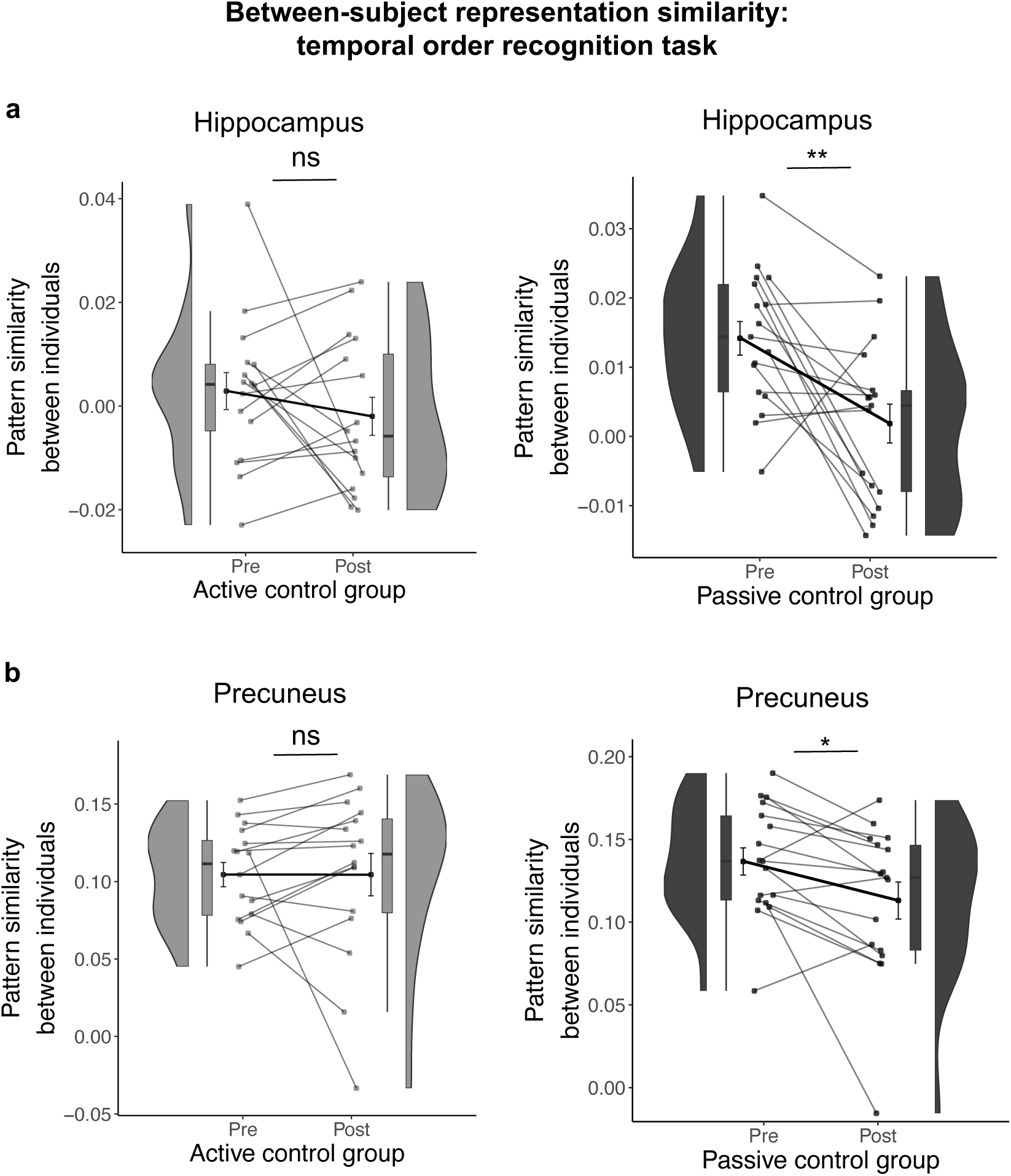
Results for between-subject ROI-based RSA during temporal order recognition, active and passive control groups. Regions-of-interest (ROIs) used in the between-subject RSA during temporal order recognition (i.e., active controls × active controls, passive controls × passive controls, analyzed separately for pre- and post-training sessions) included the (**a**) hippocampus and (**b**) precuneus. Figure legend: Data points represent individual participants (active controls, *n* = 16; passive controls, *n* = 17); boxplots display the median and upper/lower quartiles, whiskers show 1.5 interquartile ranges above/below the quartiles; circles with error bars correspond to the mean ± standard error of the mean (S.E.M.); distributions illustrate the probability density function of individual data points. Abbreviations: ns, not significant, * *p*<0.05, \*\**p*<0.01. Pattern similarity reflects Fisher’s *z*-transformed Pearson’s correlations (*r*).

**Fig. S6.**
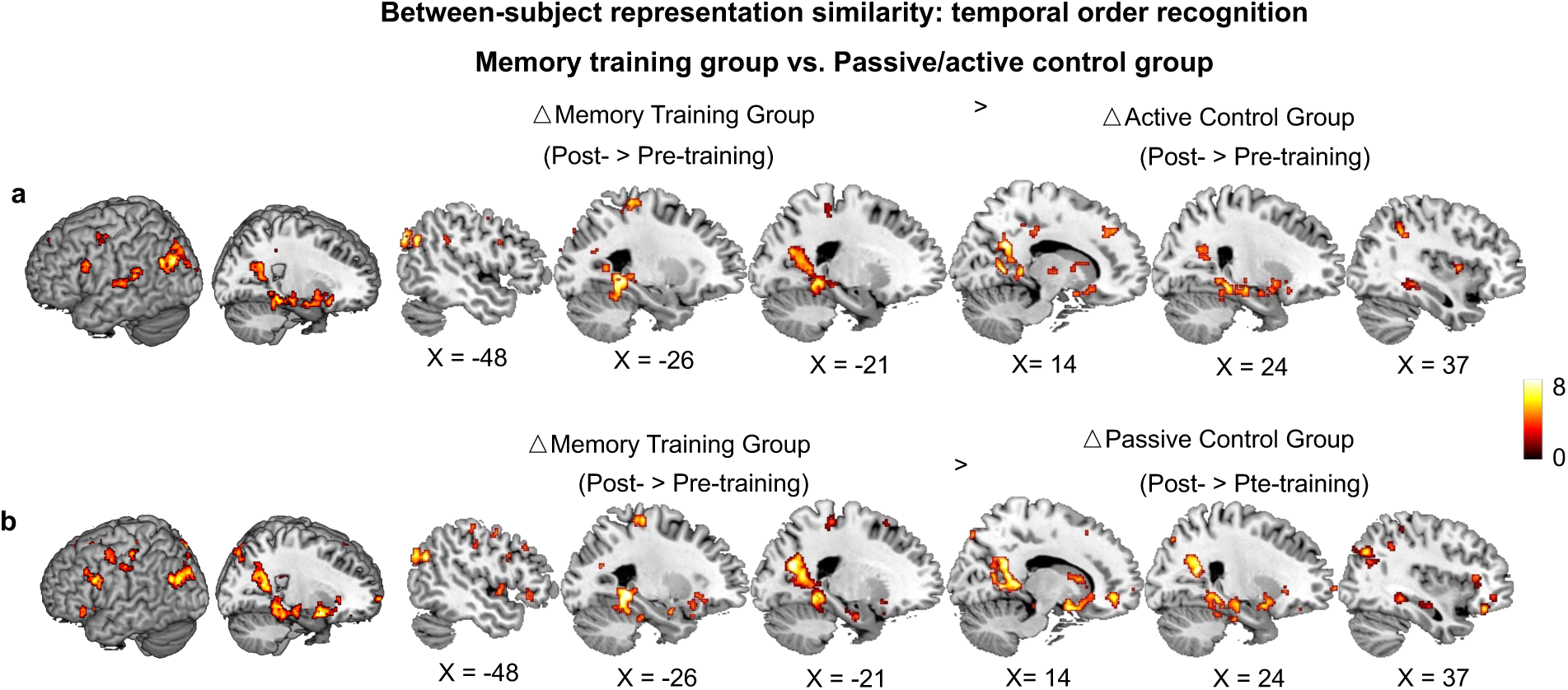
Method of loci increased pattern similarity between individuals stronger than active and passive control groups during temporal order recognition. (**a, b**) control analysis, showing whole-brain searchlight of increased pattern similarity between individuals (post-minus pre-training) in the memory training group after training, compared to (B) active or (C) passive control. Whole-brain results are reported at *p* < 0.05, family-wise error (FWE)-corrected at the cluster level (cluster-defining threshold *p* < 0.001). Colour bar indicates *t*-values.

**Fig. S7.**
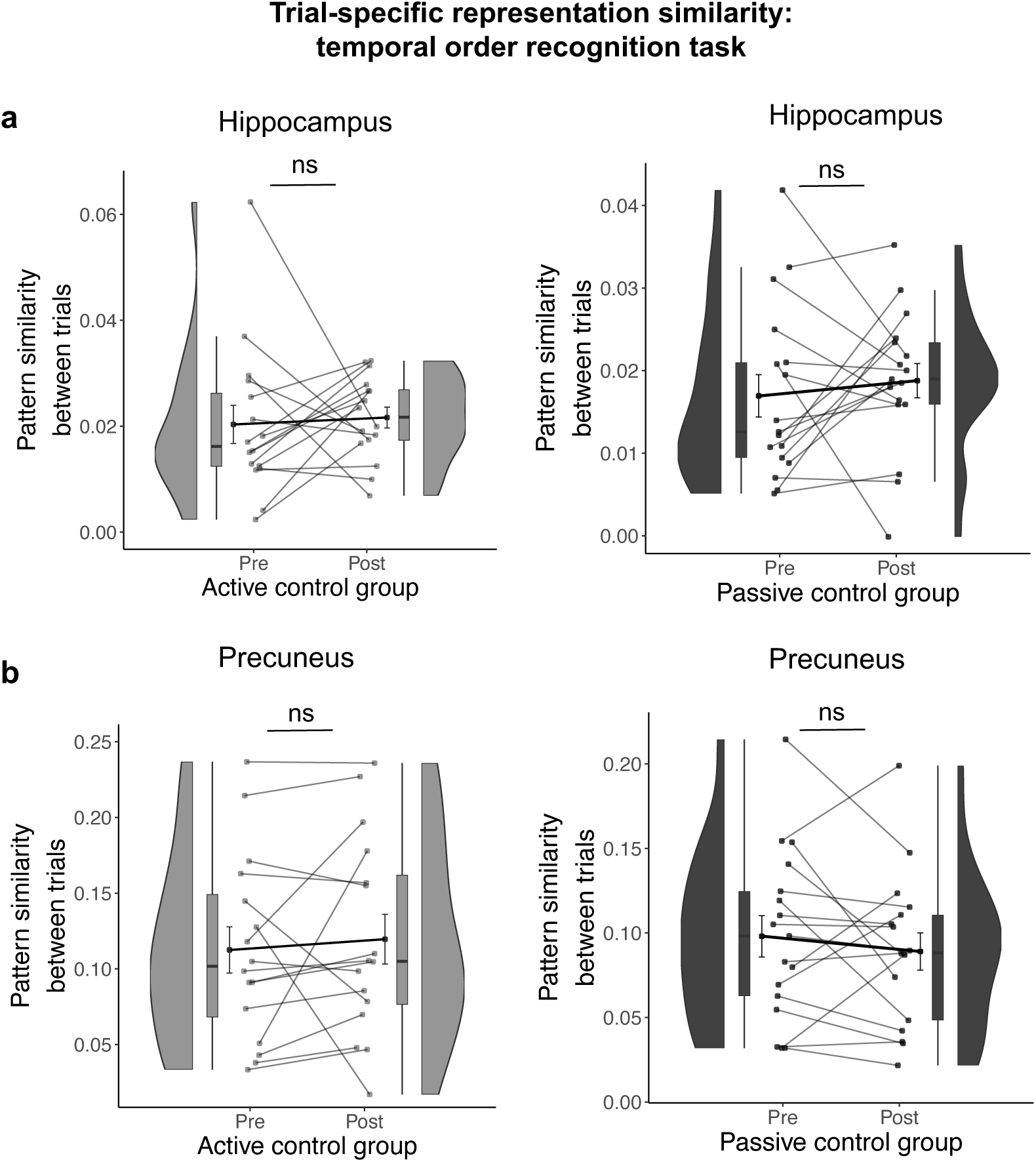
Results for trial-specific ROI-based RSA during temporal order recognition, active and passive control groups. Regions-of-interest (ROIs) used in the trial-specific RSA during temporal order recognition (i.e., active controls × active controls, passive controls × passive controls, analyzed separately for pre- and post-training sessions) included the (**a**) hippocampus and (**b**) precuneus. Figure legend: Data points represent individual participants (active controls, *n* = 16; passive controls, *n* = 17); boxplots display the median and upper/lower quartiles, whiskers show 1.5 interquartile ranges above/below the quartiles; circles with error bars correspond to the mean ± standard error of the mean (S.E.M.); distributions illustrate the probability density function of individual data points. Abbreviations: ns, not significant. Pattern similarity reflects Fisher’s *z*-transformed Pearson’s correlations (*r*).

**Fig. S8.**
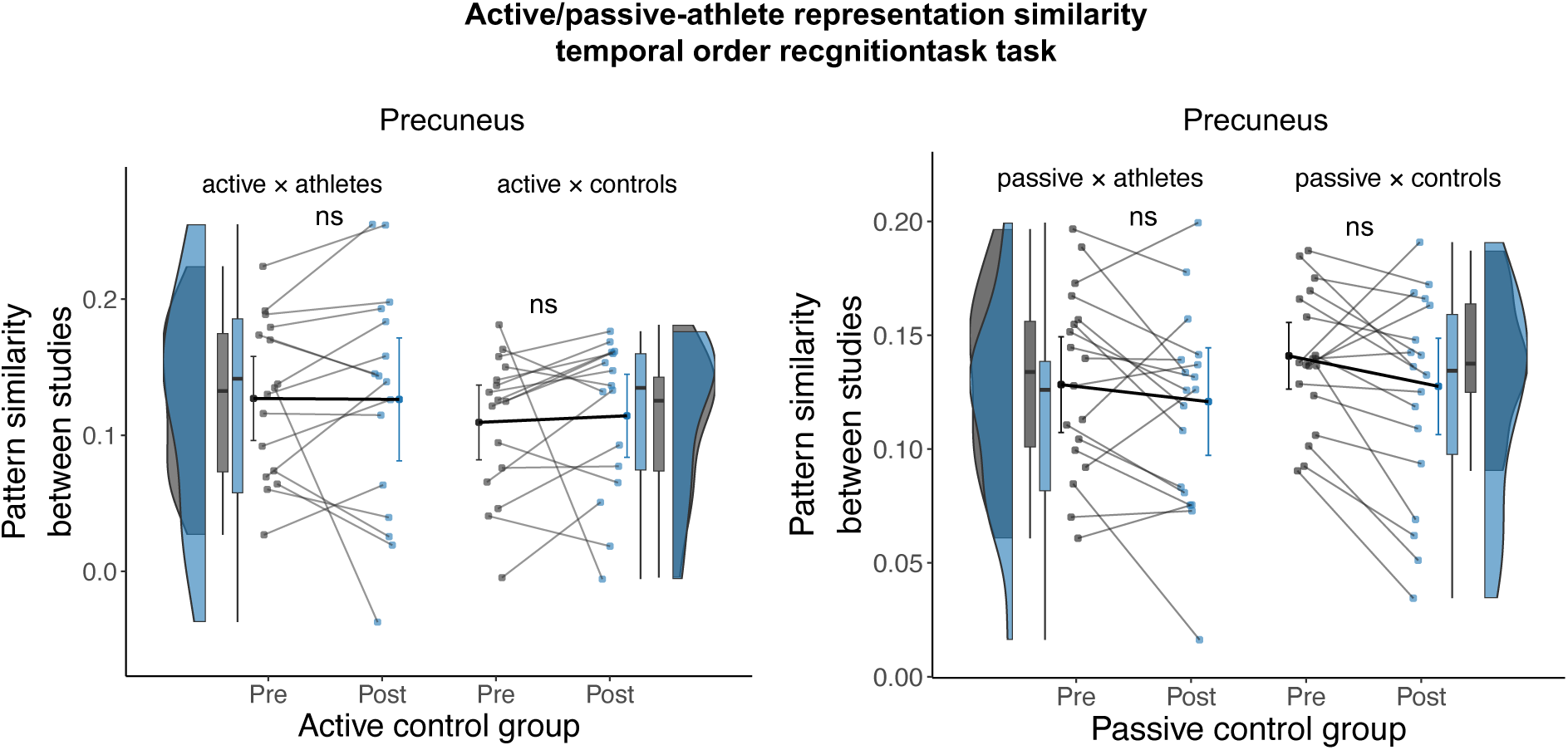
Results for trainee-athlete ROI-based RSA during temporal order recognition, active and passive control groups. Regions-of-interest (ROIs) are used in the between-group RSA during temporal order recognition, analyzed with repeat measures ANOVA of 2 × 2 factors: *time points* (pre-vs. post-training) and *group* (neural pattern similarity between active/passive controls × athletes vs. active/passive controls × matched controls). Figure legend: Data points represent individual participants (active controls, *n* = 16; passive controls, *n* = 17); boxplots display the median and upper/lower quartiles, whiskers show 1.5 interquartile ranges above/below the quartiles; circles with error bars correspond to the mean ± standard error of the mean (S.E.M.); distributions illustrate the probability density function of individual data points. Abbreviations: ns, not significant. Pattern similarity reflects Fisher’s *z*-transformed Pearson’s correlations (*r*).

**Fig. S9.**
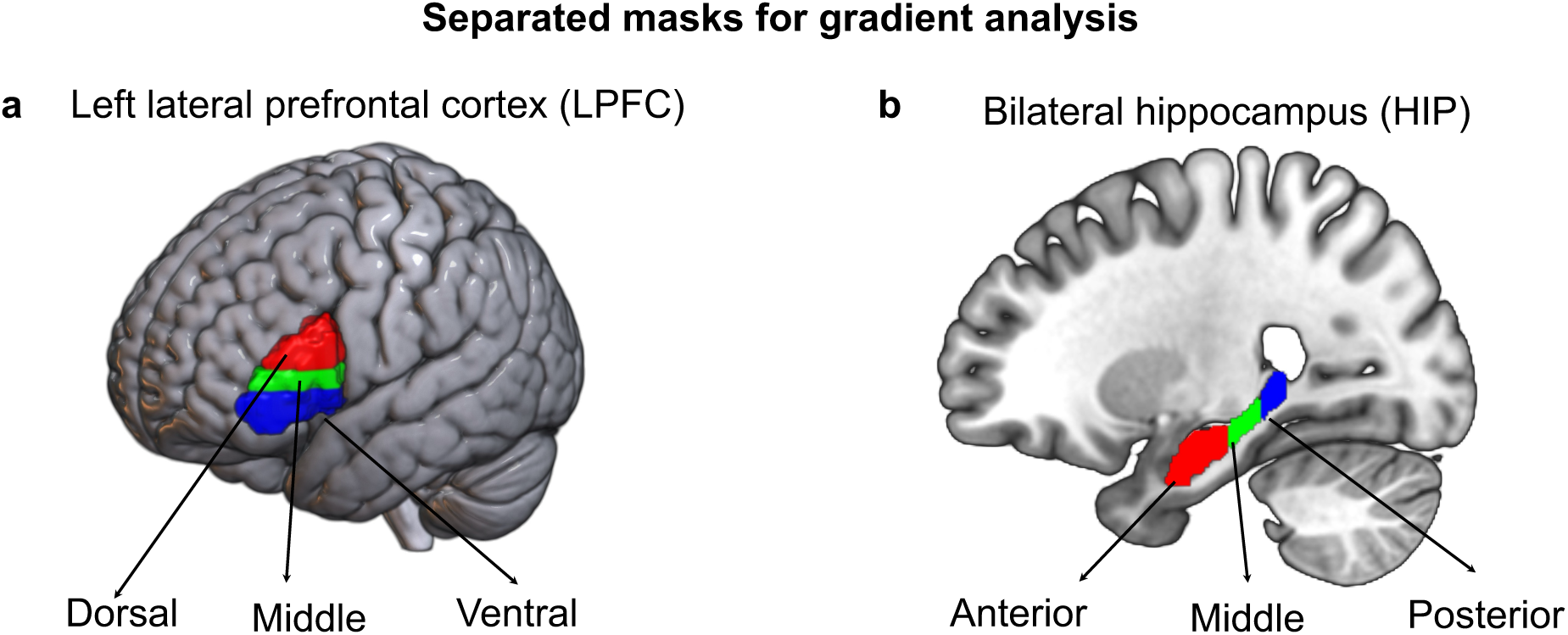
Gradient analysis of the lateral prefrontal cortex and hippocampus. Anatomical masks of (a) left lateral prefrontal cortex (LPFC) subregions based on dorsal (red), middle (green), and ventral (blue) partitions, and (b) bilateral hippocampus (HIP) subregions based on anterior (red), middle (red), and posterior (blue) partitions are shown on a template brain.

**Fig. S10.**
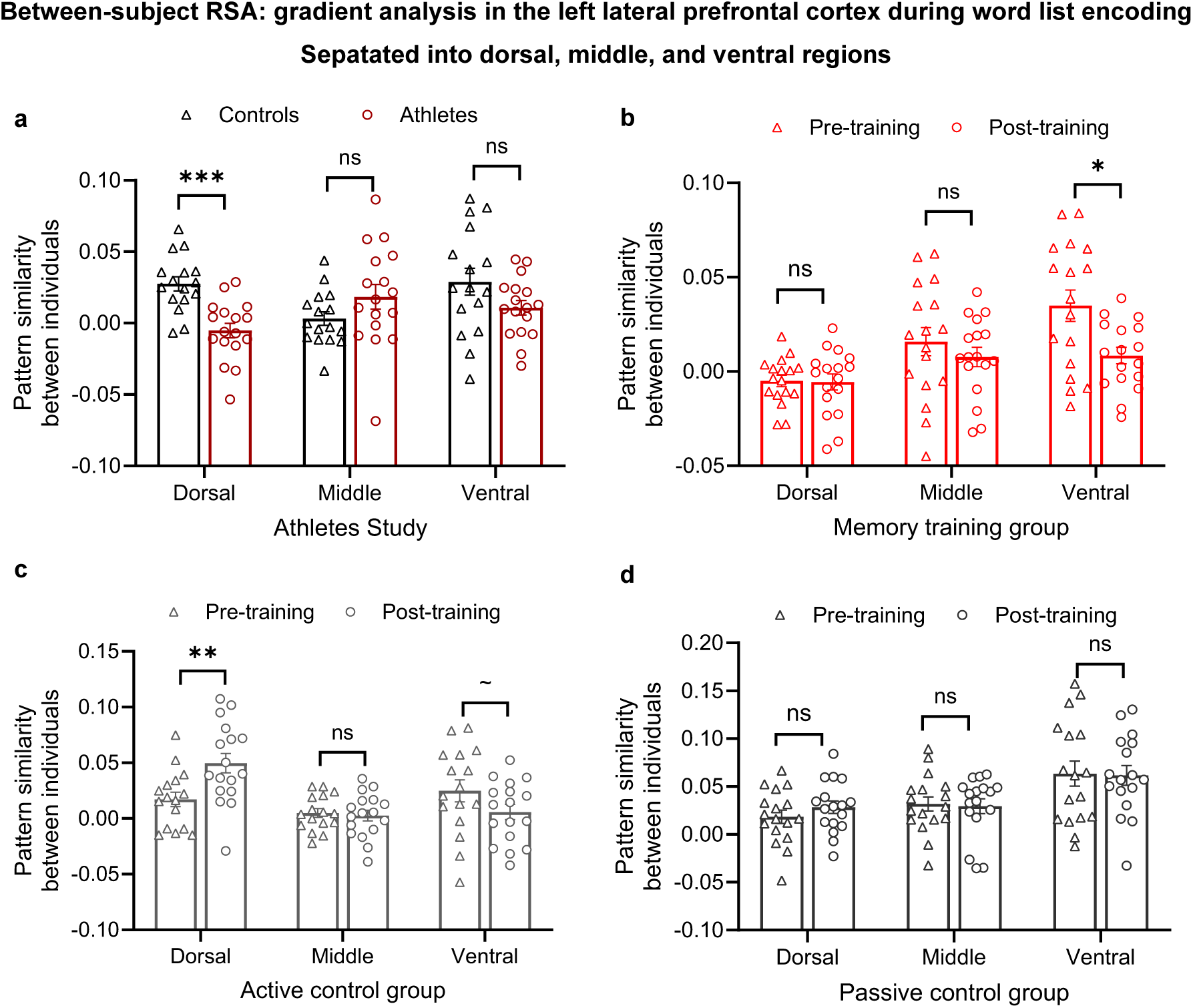
Method of loci training yields distinct neural representation between experienced individuals along the dorsal-ventral axis of the left lateral prefrontal cortex. (a, **b)** ROI results of the between-subject RSA for the memory athlete study and memory training group, and **(c, d)** analyses for the active and passive control groups. Data points represent individual participants in all panels (athlete study: *n* = 33; memory training group: *n* = 17; active control group: *n* = 16; passive control group: *n* = 17). \*\*\**p* < 0.001, \*\**p* < 0.01; \**p* < 0.017; results remained significant after false discovery rate (FDR) correction using the Benjamini– Hochberg procedure. ∼ denotes results that did not pass the correction but that exhibited a trend toward significance. *ns*, not significant. Pattern similarity reflects Fisher’s *z*-transformed Pearson’s correlations (*r*).

**Fig. S11.**
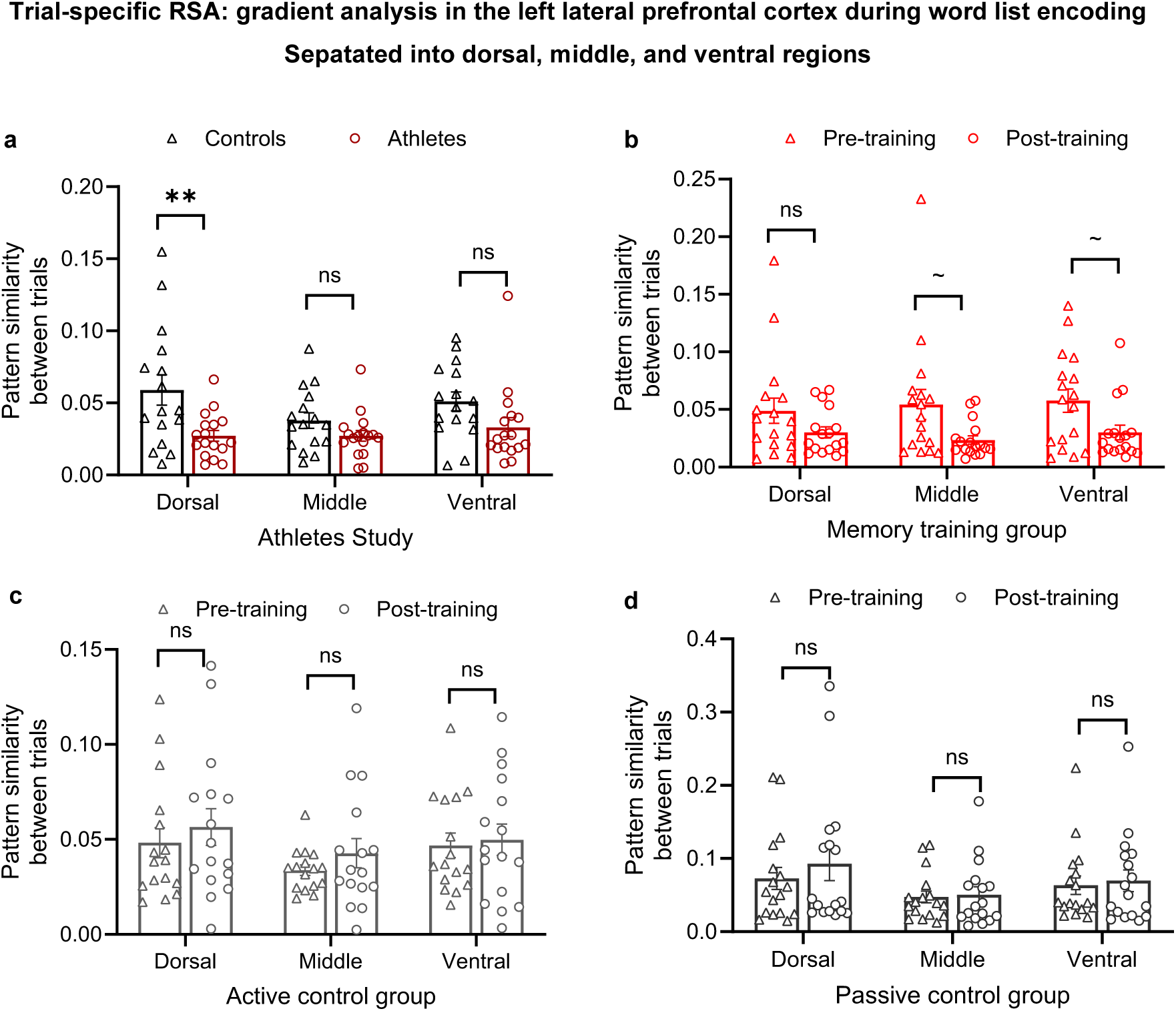
Method of loci training yields distinct neural representations between trials along the dorsal-to-ventral axis of the left lateral prefrontal cortex. (a, **b)** ROI results of the trial-specific RSA for the memory athlete study and memory training group, and **(c, d)** analyses for the active and passive control groups. \*\**p* < 0.01; results remained significant after false discovery rate (FDR) correction using the Benjamini-Hochberg procedure. ∼ denotes results that did not pass the correction but that exhibited a trend toward significance. *ns*, not significant. Pattern similarity reflects Fisher’s *z*-transformed Pearson’s correlations (*r*).

**Fig. S12.**
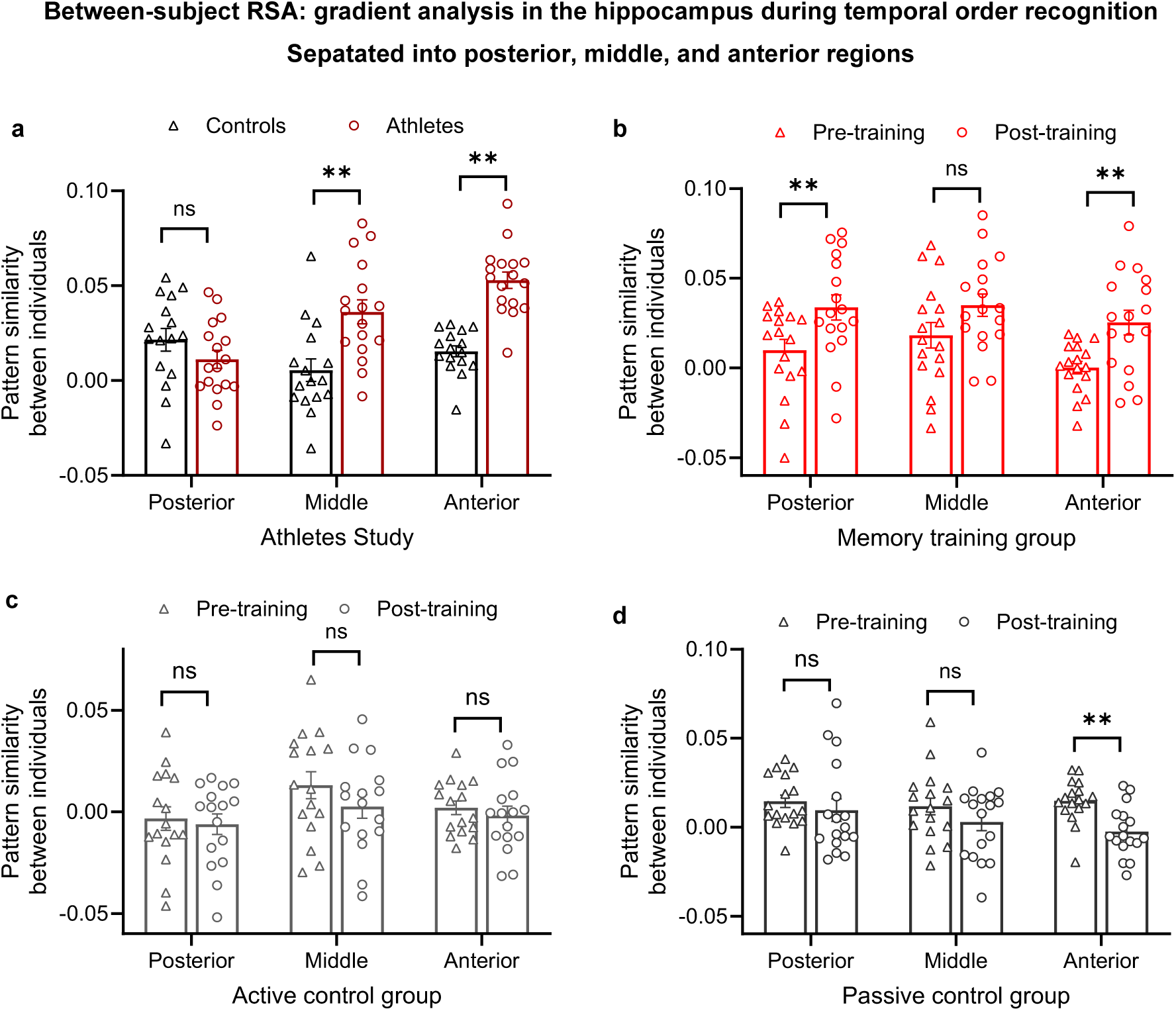
Method of loci training yields shared neural representations between experienced individuals along the posterior-to-anterior axis of the bilateral hippocampus. (a, **b)** ROI results of the between-subject RSA for the memory athlete study and memory training group, and **(c, d)** analyses for the active and passive control groups. * *p* < 0.05, \*\**p* < 0.01, ns, not significant; results remained significant after false discovery rate (FDR) correction using the Benjamini-Hochberg procedure. Pattern similarity reflects Fisher’s *z*-transformed Pearson’s correlations (*r*).

**Fig. S13.**
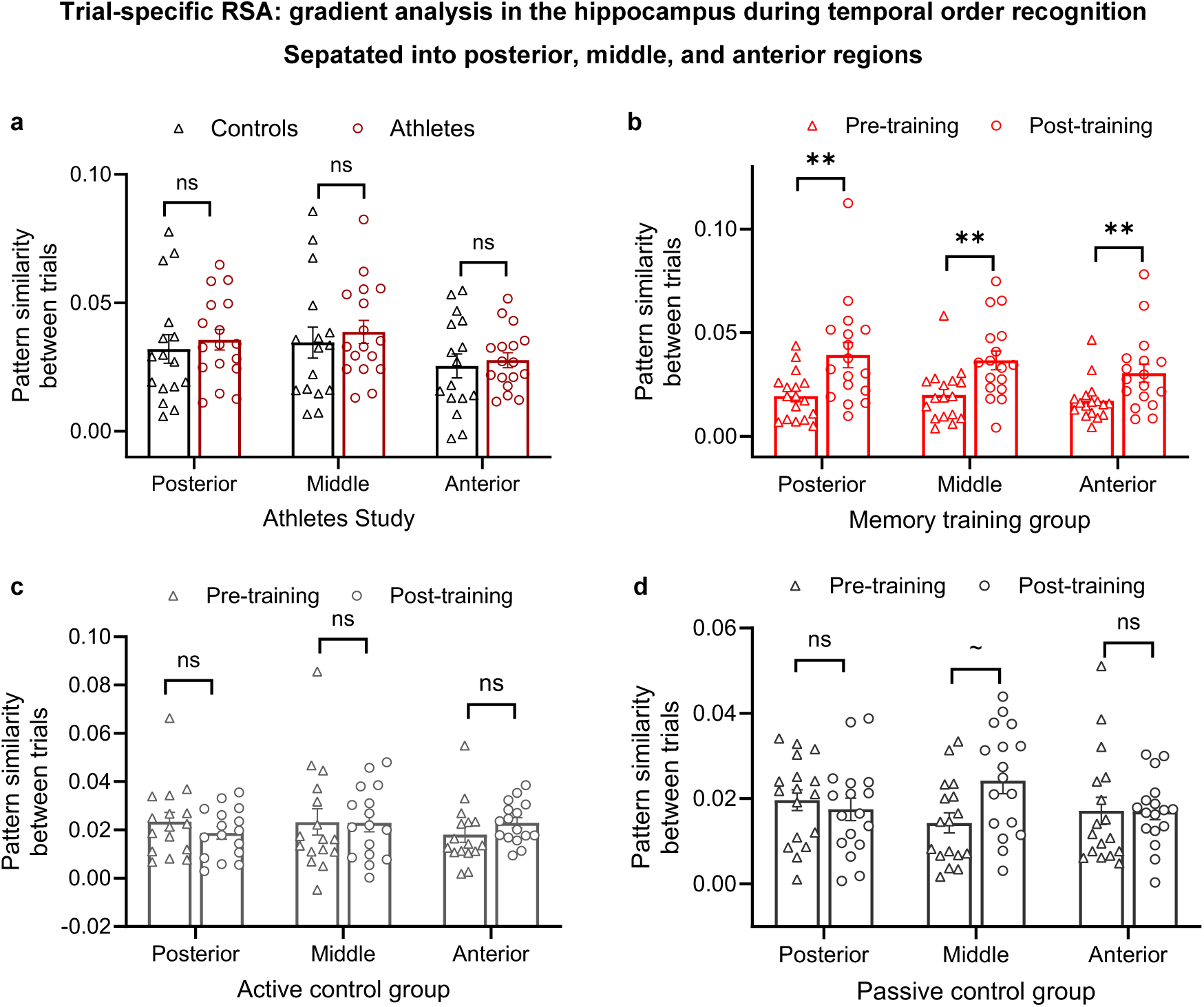
Method of loci training yields shared neural representations between trials along the posterior-to-anterior axis of the bilateral hippocampus. (a, **b)** ROI results of the trial-specific RSA for the memory athlete study and memory training group, and **(c, d)** analyses for the active and passive control groups. \*\**p* < 0.01, *ns*, not significant; results remained significant after false discovery rate (FDR) correction using the Benjamini-Hochberg procedure. ∼ denotes results that did not pass the correction but that exhibited a trend toward significance. Pattern similarity reflects Fisher’s *z*-transformed Pearson’s correlations (*r*).

**Fig. S14.**
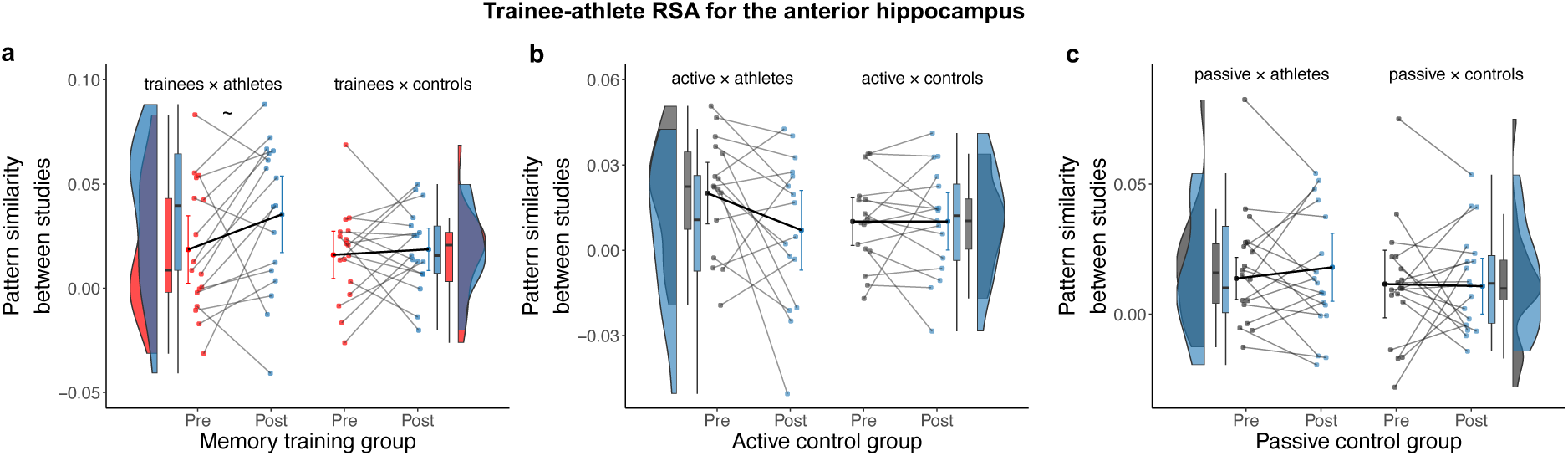
Method of loci training yields shared neural representations between trainees and athletes in the bilateral anterior hippocampus during temporal order recognition. ROI results of the comparison between trainees and memory athletes pertaining to the anterior hippocampus for **(a)** memory training group, **(b)** active control group, and **(c)** passive control group. Data points represent individual participants (athlete study, *n* = 33, memory training group, *n* = 17; active control group, *n =* 16; passive control group, *n* = 17). Results remained significant after false discovery rate (FDR) correction using the Benjamini-Hochberg procedure. ∼ denotes results that did not pass the correction but that exhibited a trend toward significance. Pattern similarity reflects Fisher’s *z*-transformed Pearson’s correlations (*r*).

**Fig. S15.**
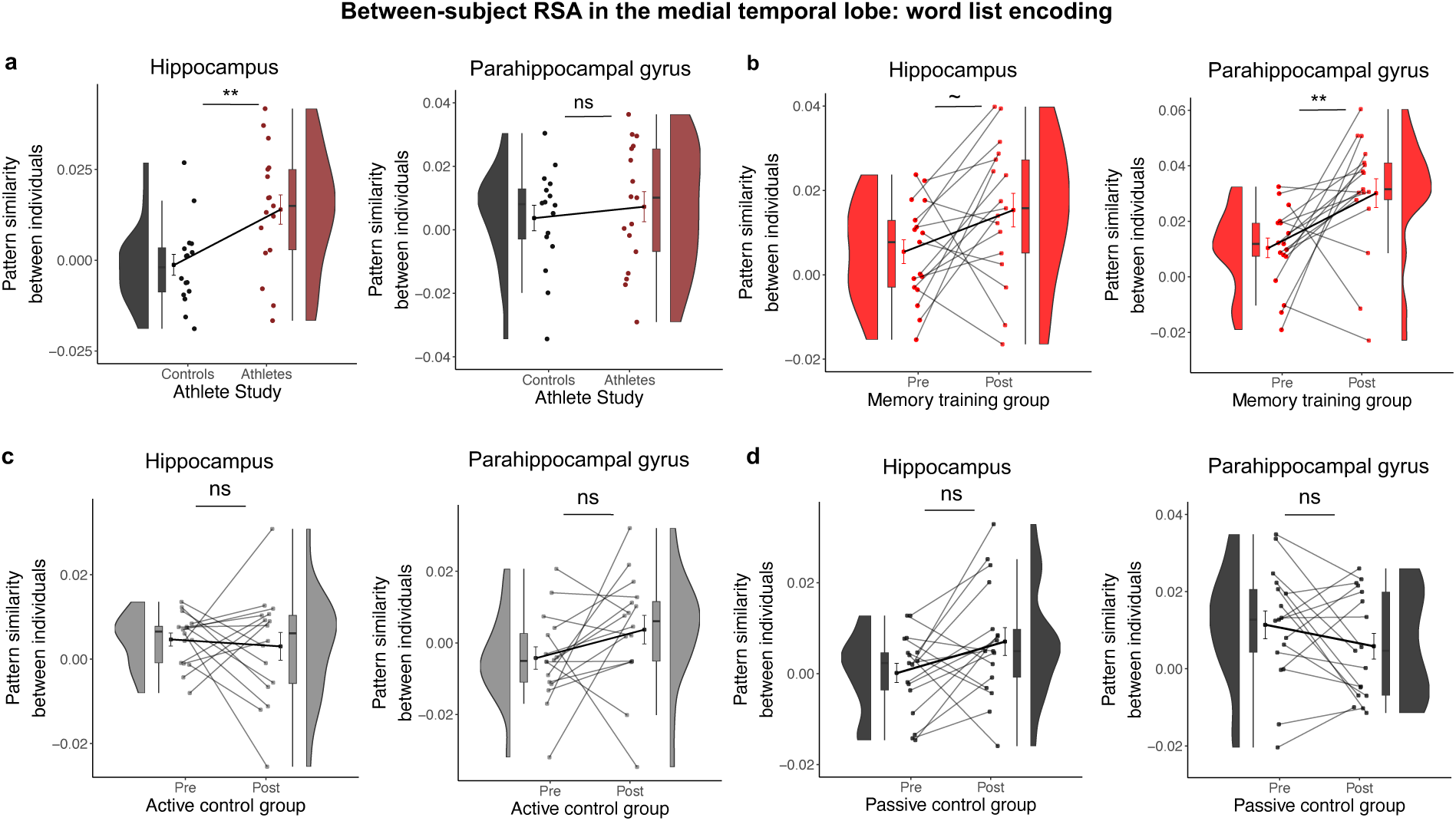
Method of loci training yields shared neural representations in the medial temporal lobe between experienced individuals during word list encoding. ROI results of the between-subject RSA pertaining to the (**a**) athlete study and (**b**) the training study, and analyses for **(c, d)** the active and passive control groups. Data points represent individual participants (athlete study, *n* = 33, memory training group, *n* = 17, active control group, *n* = 16, passive control group, *n* = 17). \*\**p* < 0.01; Results remained significant after false discovery rate (FDR) correction using the Benjamini–Hochberg procedure. ∼ denotes results that exhibited a trend toward significance. Pattern similarity reflects Fisher’s *z*-transformed Pearson’s correlations (*r*).

**Fig. S16.**
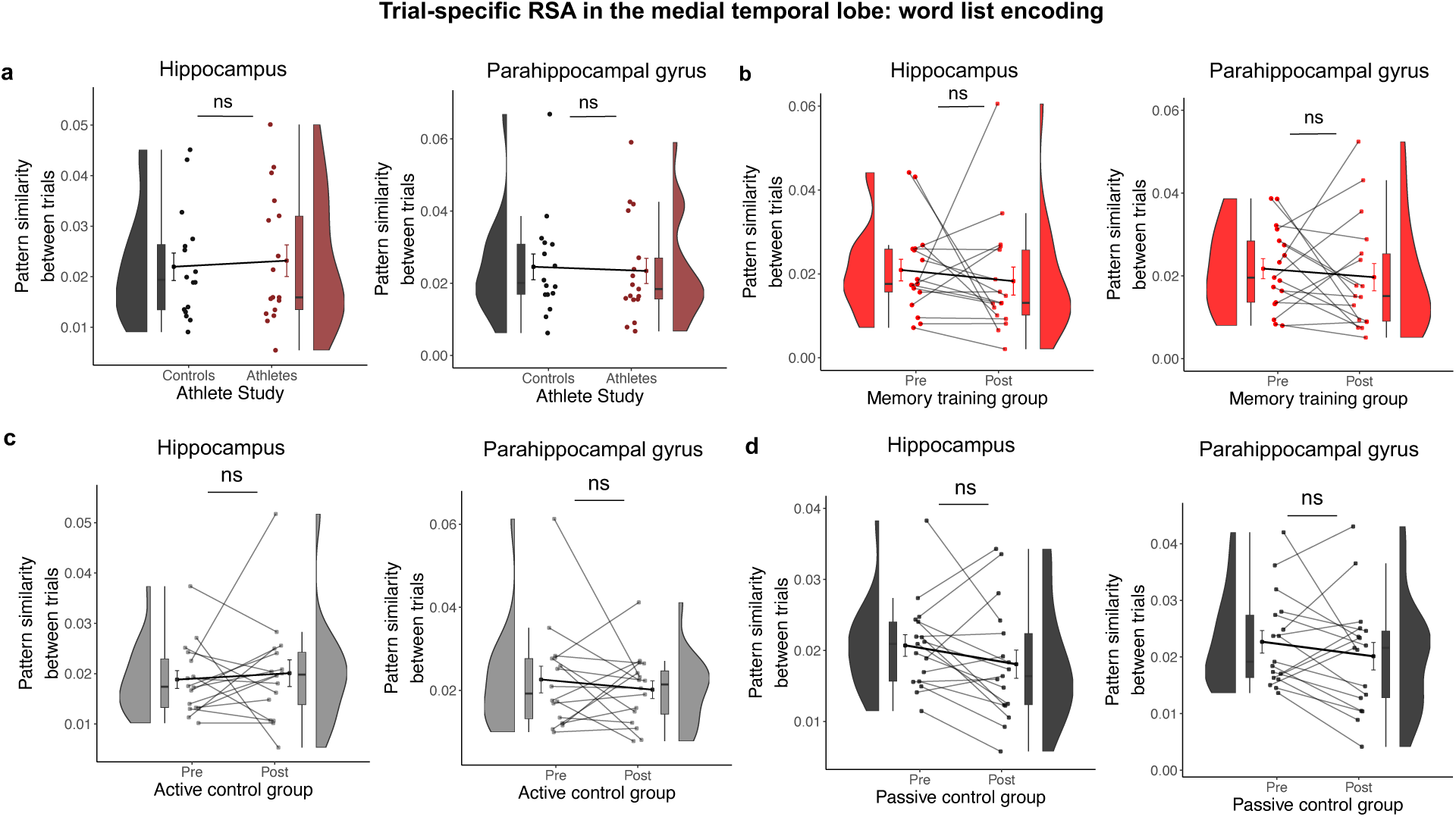
Method of loci training does not induce shared neural representations between encoded items in the medial temporal lobe during word list encoding. ROI results of the trial-specific RSA pertaining to the (**a**) athlete study and (**b**) the training study, and analyses for **(c, d)** the active and passive control groups. Data points represent individual participants (athlete study, *n* = 33, memory training group, *n* = 17, active control group, *n* = 16, passive control group, *n* = 17). *ns*, not significant. Pattern similarity reflects Fisher’s *z*-transformed Pearson’s correlations (*r*).

**Fig. S17.**
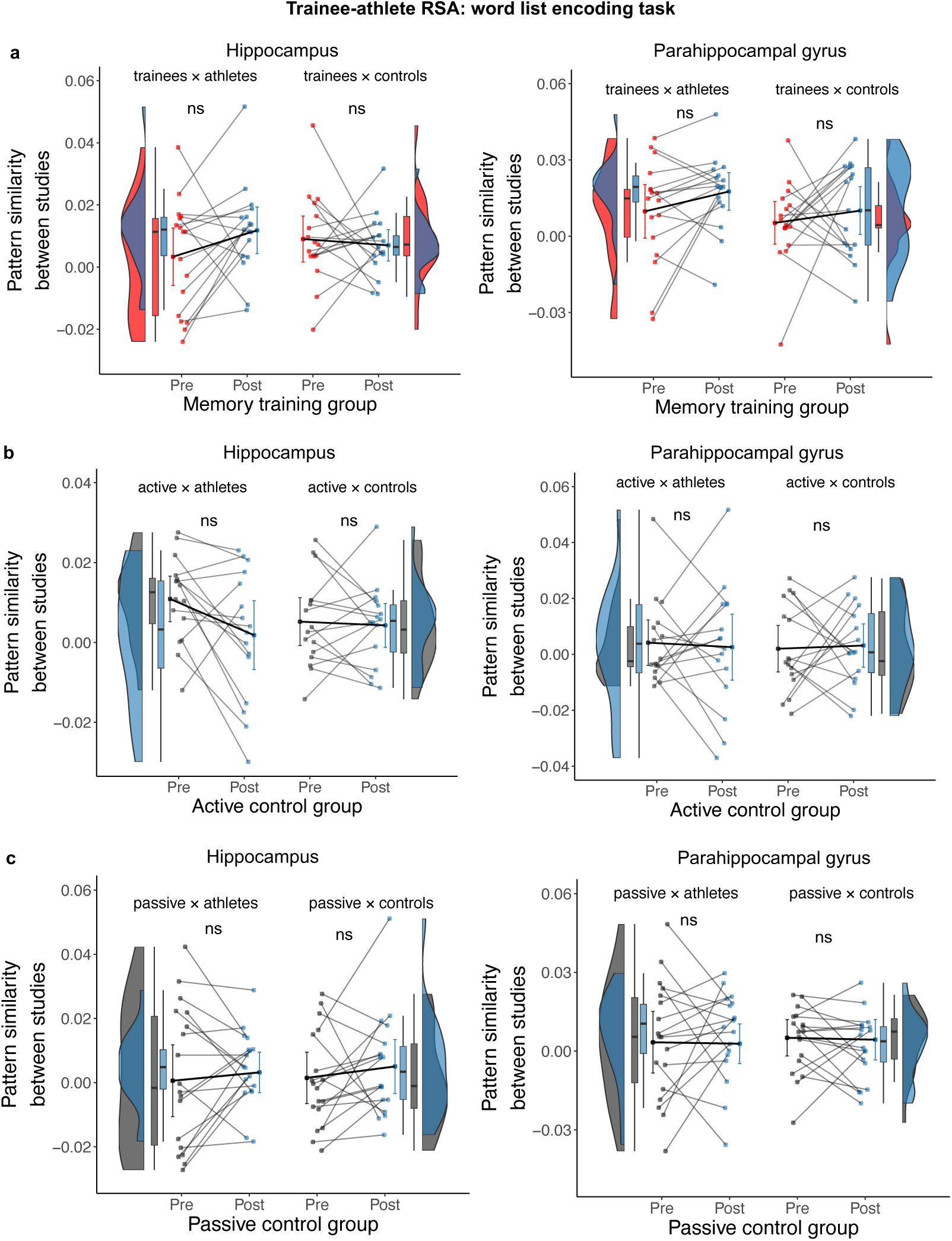
No significant effect of method of loci training on neural representations in the medial temporal lobe between trainees and memory athletes during word list encoding. ROI results of the comparison between trainees and memory athletes pertaining to the hippocampus and parahippocampal gyrus for **(a)** memory training group, **(b)** active control group, and **(c)** passive control group. Data points represent individual participants (athlete study, *n* = 33, memory training group, *n* = 17; active control group, *n =* 16; passive control group, *n* = 17). *ns*, not significant. Pattern similarity reflects Fisher’s *z*-transformed Pearson’s correlations (*r*).

**Fig. S18.**
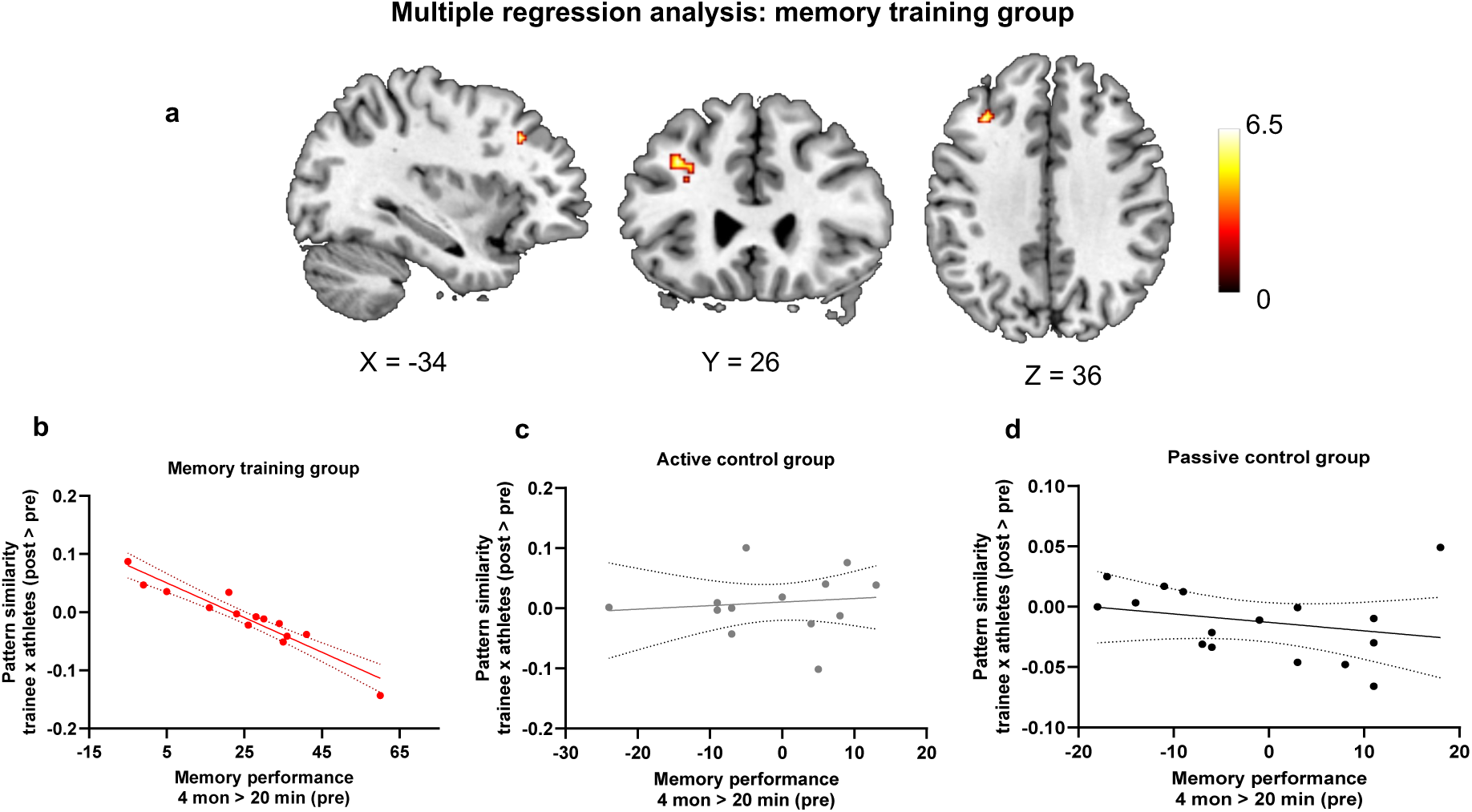
Training-induced increase in neural similarity to memory athletes was negatively associated with memory performance in the memory training group. **(a)** Voxel-wise multiple regression analysis showing contrast difference maps (Post > Pre), with individual memory improvement (number of words freely recalled at the 4-months retest minus 20-minute free recall during the pre-training session) included as a covariate of interest. A significant cluster was observed specifically in the memory training group, located in the left lateral prefrontal cortex (peak MNI coordinates: x = −34, y = 26, z = 36; cluster size = 30 voxels; *p* < 0.05, FWE-corrected at the cluster level), using a cluster-defining threshold of *p* < 0.001 and a minimum cluster size of 27 voxels. **(b-d)** Scatter plots illustrating the correlation between training-induced increases in neural similarity to memory athletes and memory improvement per group, extracted from the lateral prefrontal cortex cluster shown in (a). Plots are shown for visualization purposes only. memory training group, *n* = 14, active control group, *n* = 14, passive control group, *n* = 16. Pattern similarity reflects Fisher’s *z*-transformed Pearson’s correlations (*r*).

**Fig. S19.**
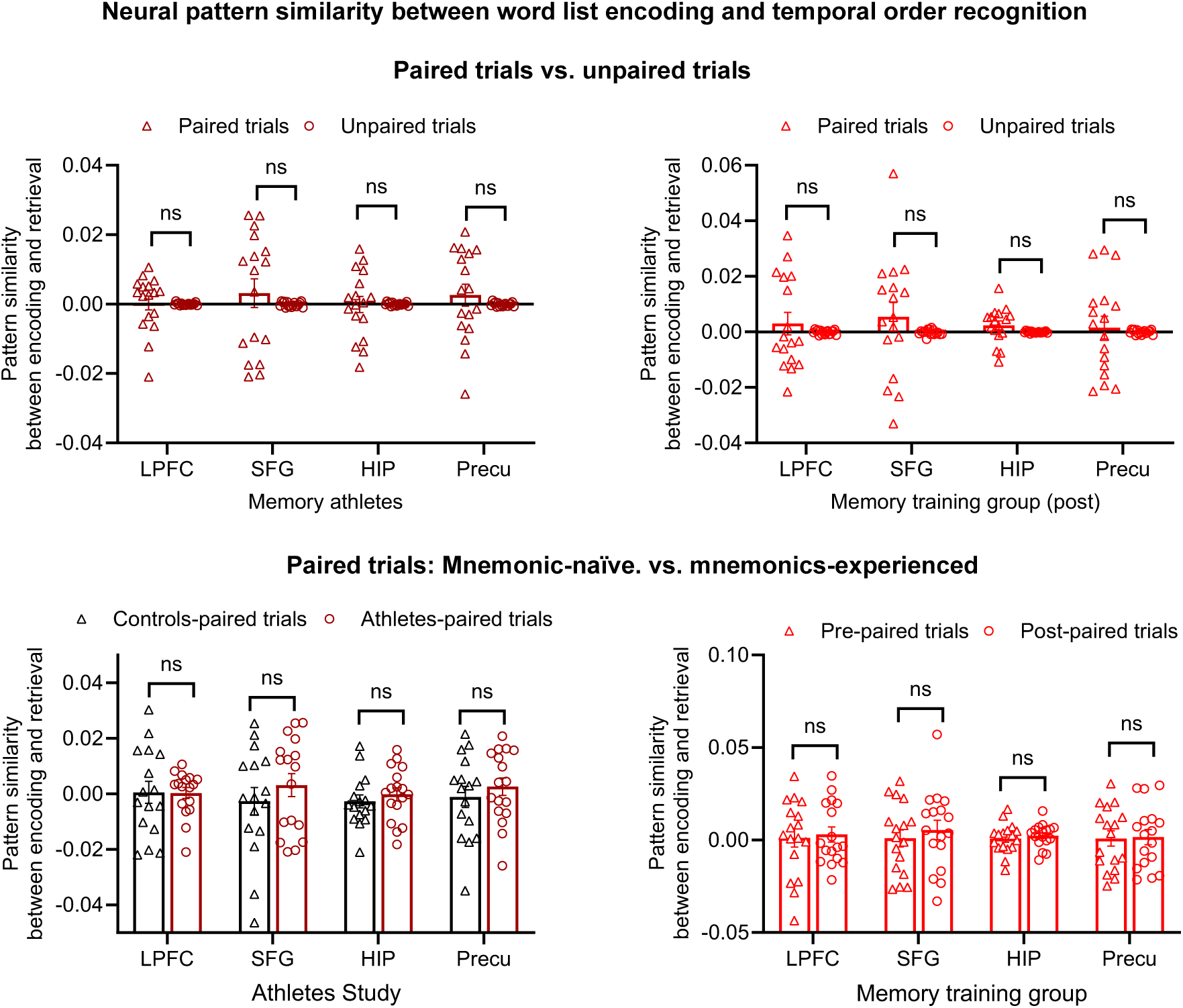
Trial-specific encoding-retrieval similarity analyses. (a, **b)** Neural pattern similarity between encoding and retrieval activity patterns for paired and unpaired trials. **(c, d)** Neural pattern similarity of paired trials across pre- and post-training sessions in the memory training group and across groups in the athlete study (memory athletes vs. matched controls). **(a-d)** data points represent individual participants (memory athlete *n* = 17; matched control *n* = 16; memory training group: *n* = 17). *ns*, not significant. Pattern similarity reflects Fisher’s *z*-transformed Pearson’s correlations (*r*).

**Fig. S20.**
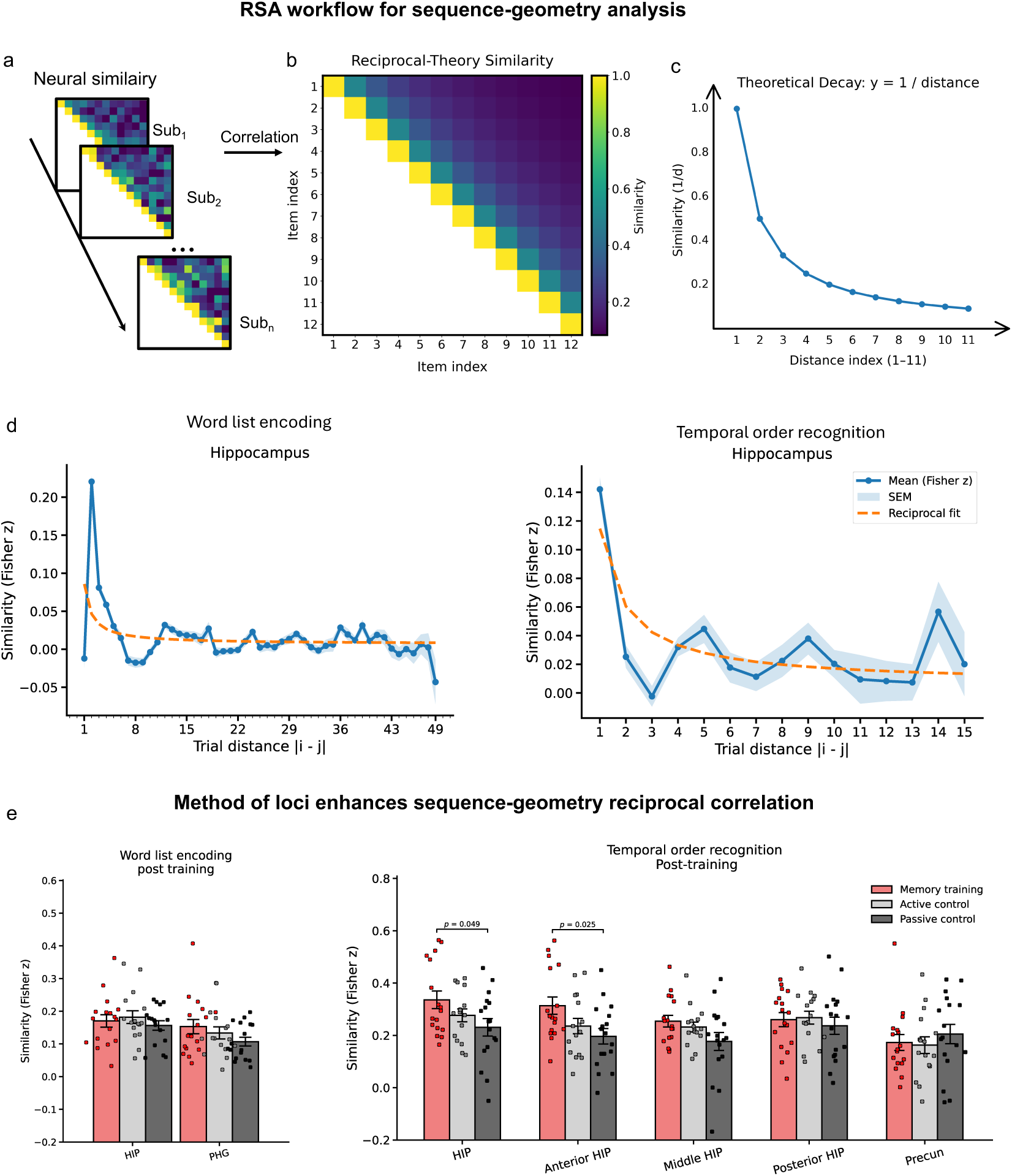
Method of loci enhances sequence-geometry of neural patterns during temporal order recognition. (**a-c**) Workflow for sequence-geometry RSA. (**a**) Neural model construction with trial-by-trial pattern similarity based on each trials’ position in the sequence (Sub, subject). (**b**) Theoretical sequence-geometry model with reciprocal distance. (**c**) Illustration of the reciprocal distance function with decreasing similarity at increasing trial distance. (**d**) Neural pattern similarity (hippocampus) as a function of trial distance in the memory training group after training during word list encoding (left) and temporal order recognition (right). (**e**) Results of the sequence-geometry analysis pertaining to word list encoding (left) and temporal order recognition (right). Error bars show mean ± standard error of the mean (SEM). Sample sizes: memory training group, *n* = 17; active controls, *n* = 16; passive controls, *n* = 17. Reported *p*-values are uncorrected for multiple comparisons; none of the effects survived false discovery rate (FDR) correction (Benjamini–Hochberg). ROIs: HIP, hippocampus; anterior, middle, and posterior HIP subdivisions; Precun, precuneus; PHG, parahippocampal gyrus.

**Fig. S21.**
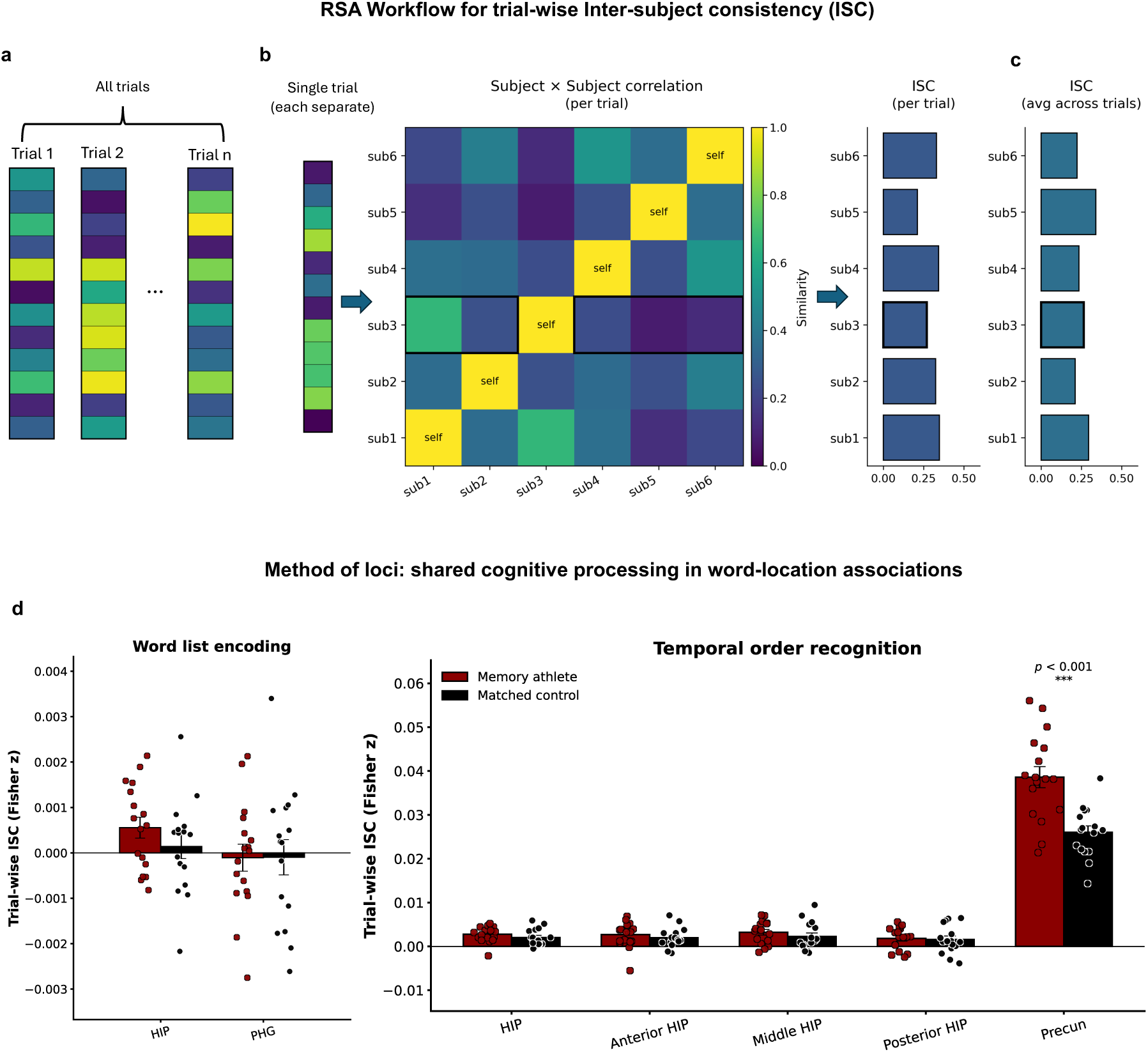
Method of loci enhances trial-wise neural pattern similarity between experienced individuals during temporal order recognition. (**a-c**) Workflow for trial-wise Intersubject Similarity (ISC) RSA. (**a**) Trial-wise beta estimates for all trials within a given ROI. (**b**) For each trial, subject, and ROI, we constructed a voxels-by-subject matrix and computed the correlation between each subject’s neural pattern with the average of all other participants, and Fisher’s *z*-transformed it, yielding a trial-level ISC value. (**c**) Trial-level ISC values were averaged across trials to produce one ISC estimate per subject and ROI. (**d**) Results of the trial-wise ISC analysis pertaining to word list encoding (left) and temporal order recognition (right). Error bars show mean ± standard error of the mean (SEM). Reported *p*-values are uncorrected for multiple comparisons; *** indicates results that remained significant after false discovery rate (FDR) correction (Benjamini-Hochberg procedure). Error bars represent mean ± SEM. Sample sizes: memory athletes, *n* = 17; matched controls, *n* = 16. ROIs: HIP, hippocampus; anterior HIP, anterior hippocampus; middle HIP, middle hippocampus; posterior HIP, posterior hippocampus; Precun, precuneus; PHG, parahippocampal gyrus.

## Supplementary Tables

### General note regarding tables displaying fMRI results

Unless stated otherwise, significance of all MRI analyses was assessed using cluster inference with a cluster-defining threshold of *p* < 0.001 and a cluster probability of *p* < 0.05 family-wise error (FWE) corrected for multiple comparisons. The corrected cluster size (i.e., the spatial extent of a cluster that is required in order to be labeled as significant) was calculated using the SPM extension “CorrClusTh.m” and the Newton-Raphson search method (script provided by Thomas Nichols, University of Warwick, United Kingdom, and Marko Wilke, University of Tübingen, Germany; http://www2.warwick.ac.uk/fac/sci/statistics/staff/academicresearch/nichols/scripts/spm/). MNI coordinated represent the location of peak voxels. We report the first local maximum within each cluster. Anatomical nomenclature for all tables was obtained from the Laboratory for Neuro Imaging (LONI) Brain Atlas (LBPA40, http://www.loni.usc.edu/atlases/, 54). We report all contrasts that yielded significant results. L, left; R, right.

**Table S1.** Demographic details. . Note that these data were presented previously (Dresler et al., 2017; Wagner et al., 2021).

|  | Athlete study |  | Training study |  |  |
| --- | --- | --- | --- | --- | --- |
|  | Memory athletes | Matched controls | Memory training | Active controls | Passive controls |
| *N* | 17 | 16 | 17 | 16 | 17 |
| Males | 9 | 9 | 17 | 16 | 17 |
| Age (years), mean ± standard deviation (SD) | 24.6 ± 4.3 | 25.4 ± 3.9 | 23.7 ± 2.7 | 24.3 ± 2.5 | 24.4 ± 3.8 |
| Age (years), range | 19-32 | 20-35 | 20-29 | 20-29 | 18-30 |
| Fluid reasoning, mean ± SD | 128.1 ± 9.6 | 128.4 ± 10.8 | 117.4 ± 12.7 | 116.4 ± 14.6 | 118.2 ± 13.2 |
| Memory abilities, mean ± SD | Not Tested | 104.6 ± 27.8 | 103.3 ± 13.3 | 100.8 ± 21.9 | 101.8 ± 16.2 |
| Left-handers | 3 | 3 | 0 | 0 | 0 |
| Smokers | 1 | 1 | 0 | 0 | 0 |

**Table. S2.** Searchlight RSA, between-subject, athlete study, word list encoding. Analysis comprised an independent *t*-test comparing memory athletes (*n* = 17) to matched controls (*n* = 16). Contrast: memory athletes < matched controls during the word list encoding > fixation baseline. Critical cluster size: 36 voxels.

| Contrast & brain regions | MNI |  |  | Z value | Cluster size |
| --- | --- | --- | --- | --- | --- |
|  | x | y | z |  |  |
| *Memory athletes < matched controls* |  |  |  |  |  |
| L superior frontal gyrus | -8 | 12 | 48 | 6.08 | 273 |
| L superior frontal gyrus | -6 | -2 | 58 | 5.02 | 93 |
| R superior frontal gyrus | 10 | 24 | 46 | 5.83 | 36 |
| L middle frontal gyrus | -38 | 18 | 34 | 5.16 | 104 |
| L middle frontal gyrus | -16 | 58 | -12 | 4.77 | 52 |
| R inferior frontal gyrus | 42 | 26 | -4 | 4.56 | 101 |
| L inferior frontal gyrus | -58 | 18 | 24 | 4.4 | 81 |
| L inferior temporal gyrus | -54 | -52 | -14 | 5.29 | 90 |
| R superior parietal gyrus | 26 | -58 | 40 | 4.86 | 58 |
| L superior parietal gyrus | -16 | -60 | 12 | 4.75 | 36 |
| L superior parietal gyrus | -26 | -68 | 50 | 4.64 | 46 |
| L precentral gyrus | -50 | 6 | 22 | 4.91 | 142 |
| L precentral gyrus | -42 | -12 | 48 | 4.81 | 54 |

**Table. S3.** Searchlight RSA, between-subject, memory training study, word list encoding. Analysis comprised an independent *t*-test comparing the memory training group (*n* = 17) pre- to post-training. Contrast: post-training < pre-training during the word list encoding > fixation baseline. Critical cluster size: 30 voxels.

| Contrast & brain regions | MNI |  |  | Z value | Cluster size |
| --- | --- | --- | --- | --- | --- |
|  | x | y | z |  |  |
| *Memory training group (post-training < pre-training)* |  |  |  |  |  |
| L superior frontal gyrus | -2 | -2 | 58 | 6.51 | 472 |
| R superior frontal gyrus | 4 | 50 | 28 | 4.75 | 80 |
| L middle frontal gyrus | -38 | 38 | 32 | 4.54 | 33 |
| L middle frontal gyrus | -22 | 22 | 38 | 4.63 | 41 |
| L inferior frontal gyrus | -38 | 20 | 2 | 4.04 | 72 |
| R postcentral gyrus | 44 | -24 | 50 | 5.21 | 182 |
| L postcentral gyrus | -58 | -12 | 44 | 4.49 | 31 |
| R precentral gyrus | 54 | 0 | 38 | 4.35 | 40 |
| R postcentral gyrus | 16 | -30 | 44 | 5.09 | 72 |
| L superior temporal gyrus | -52 | -2 | -4 | 4.47 | 123 |
| R superior temporal gyrus | 54 | -34 | 8 | 4.36 | 43 |
| R superior temporal gyrus | 60 | -14 | 8 | 4.82 | 42 |
| L middle temporal gyrus | -46 | -10 | -18 | 4.77 | 64 |
| R supramarginal gyrus | 60 | -38 | 48 | 4.66 | 34 |
| R supramarginal gyrus | 68 | -20 | 34 | 4.6 | 49 |
| R superior parietal gyrus | 22 | -58 | 36 | 4.28 | 52 |
| R precuneus | 6 | -58 | 60 | 5.09 | 137 |
| L superior occipital gyrus | -18 | -84 | 40 | 5.09 | 38 |
| L cuneus | -6 | -98 | 16 | 67 | 34 |
| L cingulate gyrus | -12 | -48 | 26 | 4.32 | 31 |
| L fusiform gyrus | -46 | -56 | -18 | 5.05 | 65 |
| cerebellum | 30 | -74 | -20 | 5.13 | 83 |
| L putamen | -16 | 10 | -6 | 4.93 | 45 |
| L caudate | -10 | 12 | 6 | 4.84 | 79 |
| brainstem | -10 | -20 | -10 | 4.47 | 67 |

**Table. S4.** Searchlight RSA, trainee-athlete, memory training study, word list encoding. The analysis used a repeated-measures ANOVA to compare neural pattern similarity between the memory training group (*n* = 17) and the memory athlete study, examining pre- and post-training effects across two factors: *time point* (pre-vs. post-training) and *group* (neural pattern similarity between trainees × athletes vs. trainees × matched controls). The interaction contrasts during word list encoding. Critical cluster size: 38 voxels.

| Contrast & brain regions | MNI |  |  | Z value | Cluster size |
| --- | --- | --- | --- | --- | --- |
|  | x | y | z |  |  |
| Word list encoding |  |  |  |  |  |
| L inferior frontal gyrus | -56 | 24 | -2 | 5.24 | 154 |
| L superior frontal gyrus | -2 | -2 | 58 | 4.22 | 42 |
| L angular gyrus | -28 | -60 | 38 | 4.57 | 42 |
| cerebellum | 14 | -54 | -16 | 4.42 | 43 |

**Table. S5.** Searchlight RSA, between-subject, athlete study, temporal order recognition. Analysis comprised an independent *t*-test comparing memory athletes (*n* = 17) to matched controls (*n* = 16). Contrast: memory athletes > matched controls during the temporal order recognition > fixation baseline. Critical cluster size: 35 voxels.

| Contrast & brain regions | MNI |  |  | Z value | Cluster size |
| --- | --- | --- | --- | --- | --- |
|  | x | y | z |  |  |
| *Memory athletes > matched control* |  |  |  |  |  |
| L parahippocampal gyrus | -26 | -38 | -14 | 6.81 | 2571 |
| R precuneus | 2 | -70 | 62 | 6.19 | 133 |
| L angular gyrus | -42 | -62 | 18 | 5.61 | 41 |
| R superior frontal gyrus | 10 | 50 | 32 | 5.59 | 70 |
| R postcentral gyrus | 68 | -16 | 22 | 5.52 | 105 |
| R inferior frontal gyrus | 54 | 10 | 4 | 5.44 | 103 |
| R lateral orbitofrontal gyrus | 36 | 36 | -12 | 5.14 | 43 |
| R middle frontal gyrus | 26 | 28 | 32 | 5.05 | 89 |
| R superior frontal gyrus | 20 | 34 | 48 | 4.82 | 41 |
| R supramarginal gyrus | 54 | -30 | 22 | 4.77 | 55 |
| R superior frontal gyrus | 4 | 68 | -4 | 4.75 | 97 |
| L middle frontal gyrus | -28 | 24 | 44 | 4.9 | 43 |
| L middle frontal gyrus | -28 | 36 | 48 | 4.38 | 42 |
| L inferior frontal gyrus | -46 | 26 | -6 | 5.26 | 65 |
| L inferior frontal gyrus | -46 | 4 | 12 | 4.07 | 57 |
| R superior temporal gyrus | 50 | 6 | -10 | 4.73 | 128 |
| R superior temporal gyrus | 50 | 24 | -18 | 4.6 | 42 |
| L inferior temporal gyrus | -32 | -4 | -26 | 4.31 | 39 |
| L superior parietal gyrus | -16 | -56 | 52 | 4.56 | 56 |
| R superior parietal gyrus | 36 | -42 | 50 | 4.35 | 45 |
| brainstem | 0 | -18 | -20 | 4.55 | 96 |
| L postcentral gyrus | -62 | -16 | 14 | 4.23 | 39 |
| R postcentral gyrus | 62 | -8 | 38 | 5.37 | 84 |
| R precentral gyrus | 38 | -10 | 42 | 4.37 | 40 |
| L middle occipital gyrus | -26 | -82 | 0 | 4.35 | 84 |
| R middle occipital gyrus | 56 | -70 | 8 | 5.17 | 63 |

**Table. S6.** Searchlight RSA, between-subject, memory training study, temporal order recognition. Analysis comprised an independent *t*-test comparing the memory training group (*n* = 17) pre- to post-training. Contrast: post-training > pre-training during the temporal order recognition> fixation baseline. Critical cluster size: 31 voxels.

| Contrast & brain regions | MNI |  |  | Z<br>value | Cluster<br>size |
| --- | --- | --- | --- | --- | --- |
|  | x | y | z |  |  |
| *Memory training group (post-training > pre-training)* |  |  |  |  |  |
| L fusiform gyrus | -28 | -46 | -16 | 6.06 | 2868 |
| L parahippocampal gyrus | -18 | -10 | -26 | 4.62 | 51 |
| R parahippocampal gyrus | 18 | -28 | -16 | 5.06 | 424 |
| L angular gyrus | -48 | -66 | 28 | 4.74 | 219 |
| R angular gyrus | 40 | -48 | 42 | 4.06 | 58 |
| R angular gyrus | 56 | -46 | 56 | 4.47 | 61 |
| L superior parietal gyrus | -14 | -80 | 50 | 3.78 | 32 |
| L precentral gyrus | -34 | -10 | 56 | 4.76 | 51 |
| L postcentral gyrus | -52 | -26 | 44 | 4.96 | 48 |
| L postcentral gyrus | -24 | -34 | 62 | 4 | 44 |
| R caudate | 16 | 18 | 6 | 4.9 | 429 |
| R caudate | 8 | 4 | 6 | 4.45 | 73 |
| L superior frontal gyrus | -12 | 10 | 54 | 4.66 | 52 |
| L superior frontal gyrus | -12 | 42 | 34 | 4.52 | 42 |
| R superior frontal gyrus | 2 | 40 | -8 | 4.11 | 44 |
| L superior frontal gyrus | -12 | 26 | 26 | 3.87 | 31 |
| L inferior frontal gyrus | -56 | 16 | 30 | 4.46 | 99 |
| R lateral orbitofrontal gyrus | 38 | 38 | -16 | 4.15 | 61 |
| brainstem | -4 | -22 | -16 | 5.11 | 125 |

**Table. S7.** Searchlight RSA, trainee-athlete, memory training study, temporal order recognition. The analysis used a repeated-measures ANOVA to compare neural pattern similarity between the memory training group (*n* = 17) and the memory athlete study, examining pre- and post-training effects across two factors: *time point* (pre-vs. post-training) and *group* (neural pattern similarity between trainees × athletes vs. trainees × matched controls). The interaction contrasts during temporal order recognition. Critical cluster size: 37 voxels.

| Contrast & brain regions | MNI |  |  | Z value | Cluster size |
| --- | --- | --- | --- | --- | --- |
|  | x | y | z |  |  |
| R parahippocampal gyrus | 22 | -32 | -14 | 5.32 | 128 |
| L middle frontal gyrus | -14 | 46 | 22 | 4.76 | 42 |
| cerebellum | -2 | -60 | -18 | 4.36 | 42 |

**Table S8.**
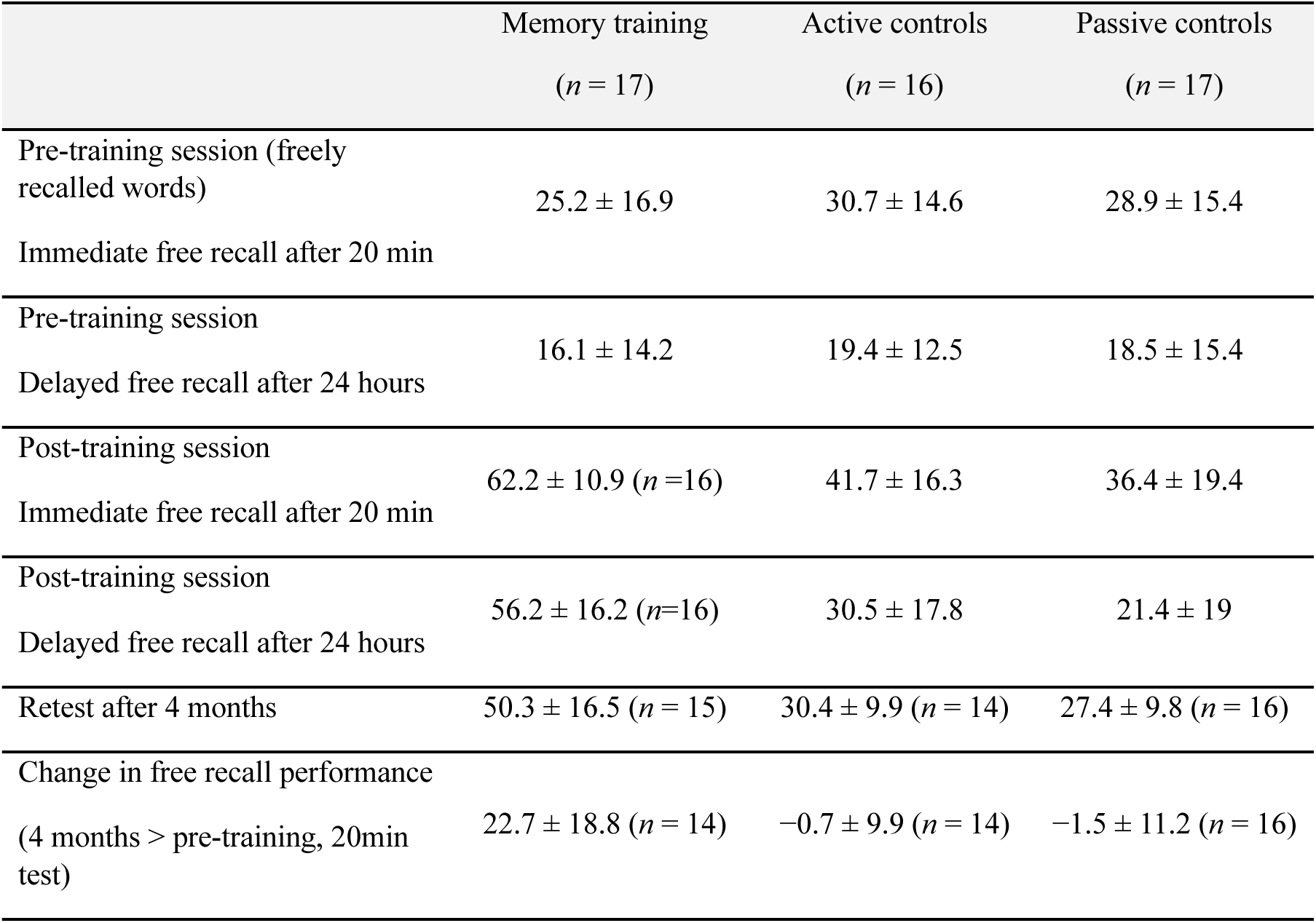
Memory performance of pre-and post-training for training study. Values represent the average number of freely recalled words ± standard deviation (SD). Note that these data were presented previously (Dresler et al., 2017; Wagner et al., 2021).

**Table. S9.**
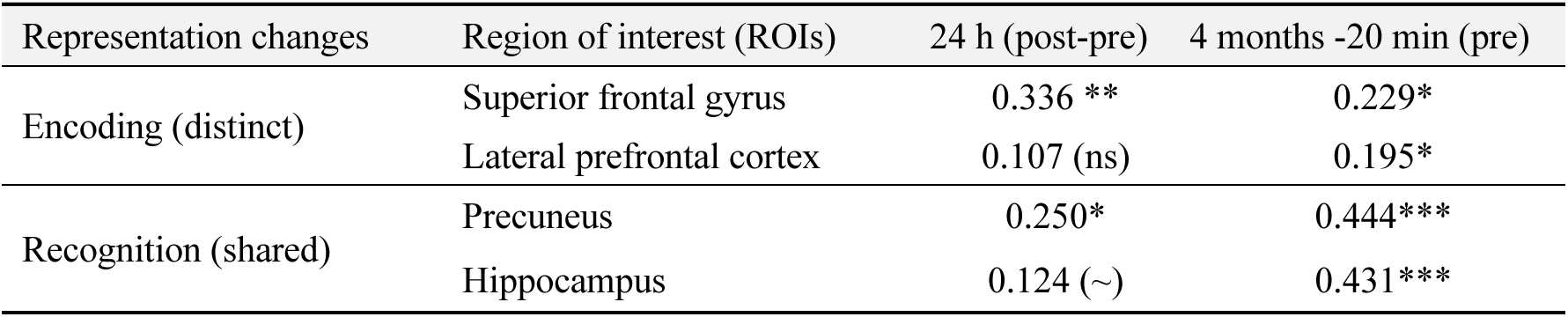
Training-related changes in neural representations are associated with better long-term memory performance after 24 hours and 4 months. Correlations of observed memory performance improved (i.e., assessed in 24 hours (*n* = 49), and 4 months (*n* = 44)), with predicted memory outcome improved from machine learning prediction analysis based on ROIs (details see Methods), both on word list encoding and temporal order recognition tasks. Notes: ∼*p* < 0.10; \**p*<0.05; \*\**p*<0.01; \*\*\**p*<0.001; ns not significant; *n* represents the sample size in corresponding calculation.

| Representation changes | Region of interest (ROIs) | 24 h (post-pre) | 4 months -20 min (pre) |
| --- | --- | --- | --- |
| Encoding (distinct) | Superior frontal gyrus | 0.336 ** | 0.229* |
|  | Lateral prefrontal cortex | 0.107 (ns) | 0.195* |
| Recognition (shared) | Precuneus | 0.250* | 0.444*** |
|  | Hippocampus | 0.124 (~) | 0.431*** |

**Table S10.** Summary of the results from the LPFC gradient analysis. All results remained significant after false discovery rate (FDR) correction using the Benjamini-Hochberg procedure. ∼ denotes results that did not pass the correction but exhibited a trend toward significance. ↑ indicates an increase in neural pattern similarity, ↓ indicates a decrease in neural pattern similarity. \*\**p* < 0.01, \**p* < 0.017.

| RSA Method | Group (Post vs. Pre) | Dorsal LPFC | Middle LPFC | Ventral LPFC |
| --- | --- | --- | --- | --- |
| **Between-subject** | Athlete vs Control | ↓*** |  |  |
|  | Memory Training |  |  | ↓* |
|  | Active Control | ↑** |  | ↓~ |
|  | Passive Control |  |  |  |
| **Trial-specific** | Athlete vs Control | ↓** |  |  |
|  | Memory Training |  | ↓~ | ↓~ |
|  | Active Control |  |  |  |
|  | Passive Control |  |  |  |

**Table S11.** Summary of the results for hippocampus gradient analysis. All results remained significant after false discovery rate (FDR) correction using the Benjamini-Hochberg procedure. ∼ denotes results that did not pass the correction but exhibited a trend toward significance. **↑** indicates an increase in neural pattern similarity, ↓ indicates a decrease in neural pattern similarity. \*\**p* < 0.01, \*\*\**p* < 0.001.

| Method | Group (Post vs. Pre) | Anterior Hippocampus | Middle Hippocampus | Posterior Hippocampus |
| --- | --- | --- | --- | --- |
| **Between-subject** | Athlete vs Control | ↑** | ↑** |  |
|  | Memory Training | ↑** |  | ↑** |
|  | Active Control |  |  |  |
|  | Passive Control |  |  | ↓** |
| **Trial-specific** | Athlete vs Control |  |  |  |
|  | Memory Training | ↑** | ↑** | ↑** |
|  | Active Control |  |  |  |
|  | Passive Control |  |  |  |
| **Trainee–Athlete** | Memory Training | ↑~ |  |  |
|  | Active Control |  |  |  |
|  | Passive Control |  |  |  |

**Table S12.** Per-group correlations with memory performance. Regions-of-interest (ROIs): LPFC, lateral prefrontal cortex, SFG, superior frontal gyrus, HIP, hippocampus, Prec, Precuneus, neural pattern similarity extracted during word list encoding (LPFC, SFG) and temporal order recognition (HIP, Prec) tasks (see also Methods section). Pearson correlation (*r*) values are given for per-group correlations between neural pattern similarity and memory performance at the 4 months retest (4 months minus free recall performance at the 20-min test during the pre-training, PRE, session), along with *p*-values.

| 4mo-20min (PRE) |  | Memory training | Active control | Passive control |
| --- | --- | --- | --- | --- |
| ROI | N | 14 | 14 | 16 |
| LPFC | Pearson Correlation | -0.406 | 0.353 | 0.072 |
|  | Sig. (2-tailed) | 0.150 | 0.215 | 0.790 |
| SFG | Pearson Correlation | 0.219 | -0.284 | 0.244 |
|  | Sig. (2-tailed) | 0.452 | 0.326 | 0.361 |
| HIP | Pearson Correlation | 0.278 | 0.363 | 0.492 |
|  | Sig. (2-tailed) | 0.335 | 0.202 | 0.053 |
| Prec | Pearson Correlation | 0.113 | 0.405 | 0.485 |
|  | Sig. (2-tailed) | 0.700 | 0.151 | 0.057 |

**Table S13.** Recognition performance (*d*-prime) per group. Performance values obtained from the temporal order recognition task during the pre- and post-training sessions, per group, showing the means (M) and standard errors of the mean (SEM).

| Training study | Pre-training session | Post-training session |
| --- | --- | --- |
|  | *d*-prime (M±SEM) | *d*-prime (M±SEM) |
| Memory training (*n* = 17) | 1.3 ± 1.3 | 2.5 ± 0.6 |
| Active controls (*n* = 16) | 1.6 ± 0.5 | 2.1 ± 1.3 |
| Passive controls (*n* = 17) | 1.6 ± 1.1 | 2.3 ± 0.9 |

**Table S14.** Word lists used during the word list encoding task.

| Word # | List 1 |  | List 2 |  | Word # | List 1 |  | List 2 |  |
| --- | --- | --- | --- | --- | --- | --- | --- | --- | --- |
|  | German | English | German | English |  | German | English | German | English |
| 1 | Bienenstock | Beehive | Diskette | Floppy disk | 37 | Apfel | Apple | Regal | Shelf |
| 2 | Holz | Wood | Ring | Ring | 38 | Steckdose | Power outlet | Zitrone | Lemon |
| 3 | Roller | Scooter | Abfall | Trash | 39 | Bett | Bed | Hund | Dog |
| 4 | Elefant | Elephant | Diamant | Diamond | 40 | Kellner | Waiter | Kiste | Crate |
| 5 | Griff | Handle | Blut | Blood | 41 | Tastatur | Keyboard | Statue | Statue |
| 6 | Vorhang | Curtain | Porto | Postage | 42 | Gold | Gols | Schloss | Castle |
| 7 | Handtasche | Handbag | Eisenbahn | Railway | 43 | Teller | Plate | Retter | Rescuer |
| 8 | Eis | Ice cream | Meer | Sea | 44 | Lkw | Truck | Lehrerin | Teacher |
| 9 | Wirbel | Whirl | Ritter | Knight | 45 | Netz | Net | Tür | Door |
| 10 | Marine | Navy | Gemälde | Painting | 46 | Beleg | Receipt | Fahne | Flag |
| 11 | Ball | Ball | Brief | Letter | 47 | Schiedsrichter | Referee | Radio | Radio |
| 12 | Palast | Palace | Schlange | Snake | 48 | Weg | Path | Nacht | Night |
| 13 | Industrie | Industry | Museum | Museum | 49 | Note | Musical note/grade | Tiger | Tiger |
| 14 | Geld | Money | Stadt | City | 50 | Pyramide | Pyramid | Limonade | Lemonade |
| 15 | Patient | Patient | Flasche | Bottle | 51 | Geweih | Antler | Tulpe | Tulip |
| 16 | Lama | Lama | Deckel | Lid | 52 | Wasser | Water | Gruppe | Group |
| 17 | Familie | Family | Kilometer | Kilometer | 53 | Schlaf | Sleep | Brot | Bread |
| 18 | Kanne | Jug | Wanze | Bug | 54 | Vogel | Bird | Reifen | Tire |
| 19 | Landschaft | Landscape | Sammlung | Collection | 55 | Zebra | Zebra | Hopfen | Hops |
| 20 | Drache | Dragon | Klebstoff | Glue | 56 | Zeitschrift | Magazine | Kaffee | Coffee |
| 21 | Farbe | Whirl | Küche | Kitchen | 57 | Teig | Dough | Schmied | Blacksmith |
| 22 | Lachs | Dishes | Storch | Stork | 58 | Popcorn | Popcorn | Schnecke | Snail |
| 23 | Spinne | Car | Bluse | Blouse | 59 | Sänger | Singer | Garten | Garden |
| 24 | Zimmer | Room | Küste | Coast | 60 | Schrott | Scrap | Biss | Bite |
| 25 | Knopf | Button | Dieb | Thief | 61 | Kirmes | Fair | Gabel | Fork |
| 26 | Schachtel | Box | Lappen | Rag | 62 | Wetter | Weather | Himmel | Sky |
| 27 | Papier | Paper | Nase | Nose | 63 | Schach | Chess | Kuh | Cow |
| 28 | Hirsch | Deer | Fax | Fax | 64 | Tinte | Ink | Wanne | Bathtub |
| 29 | Bohne | Bean | Wolle | Wool | 65 | Karte | Map | Wohnung | Apartment |
| 30 | Boden | Floor | Bühne | Stage | 66 | Hemd | Shirt | Maus | Mouse |
| 31 | Zaun | Fence | Stall | Stable | 67 | Geschirr | Dishes | Traktor | Tractor |
| 32 | Weste | Vest | Dose | Can | 68 | Auto | Car | Musik | Music |
| 33 | Sekretärin | Secretary | Krankenschwester | Nurse | 69 | Rauch | Smoke | Stuhl | Chair |
| 34 | Späne | Wood chips | Zollstock | Yardstick | 70 | U-Bahn | Metro | Bronze | Bronze |
| 35 | Lineal | Ruler | Dudelsack | Bagpipe | 71 | Frau | Woman | Mann | Man |
| 36 | Bach | Stream | Fluss | River | 72 | Socken | Sock | Knöchel | Ankle |

